# The microbiome shapes immunity in a sex-specific manner in mouse models of Alzheimer’s disease

**DOI:** 10.1101/2024.05.07.593011

**Authors:** John W. Bostick, T. Jaymie Connerly, Taren Thron, Brittany D. Needham, Matheus de Castro Fonseca, Rima Kaddurah-Daouk, Rob Knight, Sarkis K. Mazmanian

## Abstract

**INTRODUCTION**: Preclinical studies reveal that the microbiome broadly affects immune responses and deposition and/or clearance of amyloid-beta (Aβ) in mouse models of Alzheimer’s disease (AD). Whether the microbiome shapes central and peripheral immune profiles in AD models remains unknown.

**METHODS**: We examined adaptive immune responses in two mouse models containing AD-related genetic predispositions (3xTg and 5xFAD) in the presence or absence of the microbiome.

**RESULTS**: T and B cells were altered in brain-associated and systemic immune tissues between genetic models and wildtype mice, with earlier signs of immune activity in females. Systemic immune responses were modulated by the microbiome and differed by sex. Further, the absence of a microbiome in germ-free mice resulted in reduced cognitive deficits, primarily in females.

**DISCUSSION**: These data reveal sexual dimorphism in early signs of immune activity and microbiome effects, and highlight an interesting interaction between sex and the microbiome in mouse models of AD.

## 1 BACKGROUND

Alzheimer’s disease (AD) is the most prevalent neurodegenerative disease, and its incidence is expected to increase further as the global population ages [1]. Considerable progress has been made in understanding cellular and molecular contributors to AD; however, much remains unknown about its development and pathological mechanisms. Approximately two-thirds of AD cases in the United States are in females, prompting the question of how sex influences susceptibility [1]. Furthermore, although many people accumulate amyloid-β (Aβ) plaques, a key pathological hallmark of AD, only a subset will develop AD. Genetics cannot completely explain these differences [1,2]. Recent research has implicated lifestyle and environmental factors, including diet and the microbiome: the collection of microorganisms that colonize the human body [3–8]. Animal models, such as transgenic rodents, provide an opportunity to explore these questions.

AD is characterized by the accumulation of Aβ plaques and tau-mediated neurofibrillary tangles, leading to neuroinflammation, impaired synaptic communication, neuronal death, and decline in cognitive function. Two widely used models of AD are the 3x transgenic (3xTg) and 5x familial Alzheimer’s disease (5xFAD) mouse models [9,10]. These preclinical models employ overexpression of gene variants with known AD association to translate human pathophysiology into mice. Both models develop AD-associated Aβ aggregates, with a well-characterized time course. The 3xTg model, but not the 5xFAD model, also expresses the tau protein, with fibrils accumulating by 12 months [11]. While the immune response in the central nervous system (CNS) of both animal models has been extensively characterized, less is known about immune responses in peripheral tissues outside the CNS or the effect of the microbiome on immunity.

Prior evidence from AD models has revealed inflammation associated with Aβ and tau accumulation in the CNS [12]. Inflammatory responses by astrocytes and microglia, two cell types that interact closely with neurons and promote their function, may contribute to initial innate immune responses [13]. These cells are important components of protective Aβ clearance pathways. The adaptive immune response is directed at specific molecular targets that activate T and B cell receptors in an antigen-specific manner. As pathology progresses, the adaptive immune system is activated, although the initiating events, molecular targets, and timing are not well understood. Nonetheless, T and B cell responses can be protective in AD animal models, as genetic depletion of these cells results in greater accumulation of plaques and increased cytokine production [14]. Paradoxically, eliminating B cells alone is protective against AD-like progression in mice, indicating a B cell contribution to pathogenic processes [15]. Thus, the protective and pathogenic contributions of T and B cells to AD require further study.

The intestinal microbiome influences outcomes related to AD [7,16]. In AD mouse models, germ-free (GF) conditions or antibiotic treatment reduces Aβ aggregates in the brain [3–5]. In addition, the microbiome regulates inflammatory responses in AD [6,7]. Microbes and microbe-produced factors may affect AD in several ways, but one important aspect is activating immune cells [17,18]. Previous studies have identified that the microbiome may influence AD outcomes in a sex-specific manner and could involve hormone-microbiome-AD pathology interactions [19–24]. However, understanding the mechanisms by which the microbiome exerts these influences on AD requires further investigation.

Herein, we investigated immune responses in mice with AD-associated genetic landscapes in the presence or absence of a microbiome by comparing GF mice to conventionally raised specific pathogen-free (SPF) mice, i.e., those with a normal laboratory microbiota, with the same genotype. We found that adaptive immune responses were elevated in multiple tissues and varied by sex and microbiome status. Specifically, we show that the time of onset of the adaptive immune response in brain-draining lymph nodes is sex-dependent, occurring earlier in female mice. In the 3xTg model, immune responses were dramatically elevated when males, but not females, were kept germ-free. In contrast, the 5xFAD model showed increased immune activity in females compared to males in SPF conditions. Importantly, the absence of a microbiome attenuated immune responses and improved cognitive function, primarily in female mice in both models. Our results demonstrate that the microbiome and sex interact to influence immunity and cognition in mouse models of AD.

## 2 METHODS

### 2.1 Mice

Wildtype C57BL/6J (The Jackson Laboratory, Cat#000664) mice were obtained from Jackson Laboratory at 8 weeks of age. Homozygous 3xTg [B6;129-Tg(APPSwe,tauP301L)1Lfa *Psen1^tm1Mpm^*/Mmjax] and hemizygous 5xFAD [B6.Cg- Tg(APPSwFlLon,PSEN1*M146L*L286V)6799Vas/Mmjax] (The Jackson Laboratory, Cat#004807 and Cat#034848) mice were maintained in a colony in the laboratory of S.K.M at the California Institute of Technology (Caltech). 5xFAD hemizygous mice and wildtype littermates were produced by crossing transgenic and C57BL/6J mice. C57BL/6J, 3xTg, and 5xFAD mice were rederived as germ-free (GF) in the Caltech gnotobiotic facility. After weaning, mice were housed together with littermates until experiments. All experiments were performed with both female and male mice. For behavioral experiments, investigators were not blinded to group.

Experimental mice were housed in sterilized microisolator cages and maintained on ad libitum autoclaved 5010 PicoLab Rodent Diet (LabDiet, Cat#5010) and sterilized water. Germ-free mice were maintained on water containing antibiotics during behavioral tests [ampicillin (1 mg/mL); vancomycin (0.5 mg/mL); neomycin (1 mg/mL); 1% sucrose]. Specific pathogen-free (SPF) control mice were provided water containing 1% sucrose during behavioral tests. Animals were group housed (2–5 mice per cage) unless otherwise specified. Conditions in the animal housing facilities were maintained at 21-24 °C, 30-70% humidity, with a cycle of 13 hours light, 11 hours dark. All experiments were performed with approval from the Institutional Animal Care and Use Committee (IACUC) of Caltech (Protocol IA20-1798).

### 2.2 Immune cell isolation and profiling

#### 2.2.1 Intestinal lamina propria immune cell isolation

For isolation of intestinal lamina propria cells, the small and large intestines were dissected and placed immediately into ice-cold phosphate-buffered saline (PBS). After mesenteric fat and Peyer’s patches (small intestine) were removed, the intestines were longitudinally opened, and luminal contents were washed out with cold PBS. Tissue pieces were washed for 10 min. in 1 mM dithiothreitol (DTT)/PBS at room temperature on a rocker to remove mucus, followed by a wash for 25 min. in 10 mM EDTA/30 mM HEPES/PBS at 37 °C on a platform shaker (180 rpm) to remove epithelium. After a 2 min. wash in complete RPMI, the tissue was digested in a six-well plate for 1.5 hr. in complete RPMI with 150 U/mL (small intestine) or 300 U/mL (large intestine) collagenase VIII (Sigma-Aldrich) and 150 µg/mL DNase (Sigma-Aldrich) in a cell culture incubator (5% CO_2_). Tissue digests were passed through a 100 μm cell strainer and separated by centrifugation (1250 ×g for 20 min.) using a 40/80% Percoll gradient. Immune cells were collected at the 40/80% interface and washed with HBSS wash buffer [1X HBSS without phenol red, Ca^2+^, or Mg^2+^; 10mM HEPES buffer, DNase I (50 µg/ml), 1 mM MgCl_2_, and BSA Fraction V (2.5 mg/ml)] before surface staining, fixation (eBioscience Foxp3 / Transcription Factor Staining Buffer Set), and intracellular staining.

#### 2.2.2 Spleen and lymph node immune cell isolation

For the spleen and lymph nodes, the tissue was passed through a 100 μm cell strainer and washed with HBSS wash buffer. The spleen suspension was centrifuged at 500 ×g for 5 min., resuspended in red blood cell lysis buffer (Sigma-Aldrich), and incubated for 8 min. at room temperature. Spleen and lymph node cell suspensions were washed with 0.5% BSA/PBS before surface staining and fixation (eBioscience Foxp3 / Transcription Factor Staining Buffer Set).

#### 2.2.3 Cell counting and normalization

Single cell suspensions of tissues were washed and cells resuspended in 1 mL MACs buffer. 10 µL of this volume was diluted in 190 µL MACS buffer. Flow cytometry was performed on a Beckman Coulter CytoFLEX S flow cytometer. Each sample was run for 30 s at a set flow rate of 60 µL/min., so that 30 µL was used for cell counting. To calculate cells counts, FlowJo was utilized to gate cells and exclude potential debris. The gated cell count was used to calculate the total cells for each tissue with the following formula:

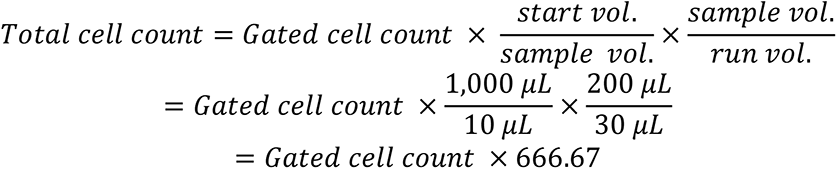

#### 2.2.4 Immune cell profiling by flow cytometry

CD16/32 antibody (eBioscience) was used to block non-specific binding to Fc receptors before surface staining at 4 °C for 30 min. Immune cells were stained with antibodies (eBioscience, BioLegend) for surface markers (1:200 dilution): CD19 (FITC), CD3e (PE), CD4 (APC; BV510), CD45.2 (BV421), CD8a (APC-e780), and TCRβ (PerCP-Cy5.5) and intracellular markers (1:100 dilution): Foxp3 (APC), GM-CSF (PE), IFNγ (FITC), IL-17A (PE), and IL-4 (PE-Cy7). Live and dead cells were discriminated by Live/Dead Fixable Violet or Aqua Dead Cell Stain Kit (Invitrogen). Mean fluorescence intensity (MFI) was calculated as the median value of the channel gated on the positive cell population.

### 2.3 Behavior Testing

Noldus EthoVision software (Wageningen, Netherlands) was used to record and analyze animal behavior.

#### 2.3.1 Y Maze

Spatial memory was assessed by spontaneous alternation measurement in a three-arm open maze as described previously [25]. Mice were acclimated in the behavior room for thirty minutes prior to testing. Each arm was assigned a letter (A, B, or C). Mice were placed into Arm C and then allowed a period of five minutes to explore the maze. At the end of five minutes, the mouse was removed from the maze. The maze was thoroughly cleaned with 0.5% hydrogen peroxide disinfectant and allowed to dry to remove olfactory cues. A camera mounted above the center of the maze was utilized to identify the center point of each mouse. Arm entries and re-entries were counted automatically with EthoVision software. An arm entry was counted if the center point of the mouse crossed the threshold between the arm and center. If a mouse entered an arm momentarily, returned to the center of the maze, and then entered the same arm again, this was considered as two entries to the same arm. Spontaneous alternations were classified as successive entries into three successive arms without error. Percent alternations was calculated with the following equation:

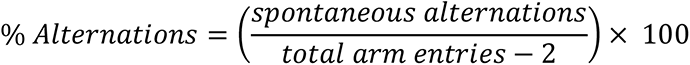

Mice were moved into the behavior room thirty minutes prior to assessment on day one to acclimate. Mice were assessed over two days and housed overnight in the behavior room between assessment days. Mice were placed into a box with two similar objects, either two 8 x 4 inch colored blocks, or two cell culture flasks filled with sand, and allowed to explore the box for ten minutes as described previously [26]. The initial objects were referred to as the “familiar objects.” Half the mice were familiarized to colored blocks and half to flasks to avoid object bias. The following day, one of the familiar objects was replaced with a “novel object” of the opposite type and the mice were allowed to explore the field and objects for an additional ten minutes. Time spent at each object was recorded with EthoVision software. The following formula was used to measure novel object recognition:

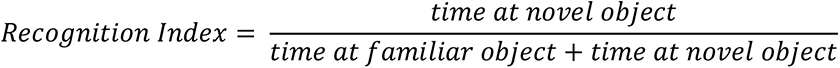

#### 2.3.2 Barnes Maze

We followed the shortened Barnes Maze protocol outlined in Attar et al. [27]. The procedure included a habituation period, four training periods, and a final test period. All mice were habituated for a minimum of thirty minutes in the behavior room to allow for adjustment to their surroundings. We maintained a minimum of one hour between the habituation period and training periods. Each cohort completed a habituation period and two training periods on day one. Two more training sessions were completed on day two. Mice were rested on day three and tested on day four. The Barnes Maze has eighteen equally spaced holes around the perimeter of a circular maze. The goal box was always aligned in the same position relative to visual cues in the room. Four reference objects, in addition to the natural asymmetry of the behavior room, were used during all periods of testing to orient the mice in the room. In the habituation phase, mice were guided to a dark goal box located beneath the target hole. Aversive stimuli, including bright diffuse light, fan-induced air currents, and white noise provided encouragement for the mice to find the goal box during the trainings and test. During each training, mice were allowed 180 seconds to locate and enter the goal box. When the mouse entered the goal box, the aversive noise machine was turned off via remote control. Mice were allowed twenty seconds to further acclimate to the goal box before being returned to their home cage. If the mouse did not enter the goal box on their own within 180 seconds, they were guided to the goal box and allowed to acclimate for twenty seconds without the aversive noise. They were then returned to their home cage. On the test day, the goal box was removed, and the mice were allowed to roam for 120 seconds. We assessed the latency in seconds required for mice to reach the goal box during the trainings or the location of the removed goal box on test day.

### 2.4 Statistical Methods

Unless otherwise noted, statistical analyses were performed with the non-parametric Mann-Whitney U test for behavior assays and two-way ANOVA for immune response data on individual biological samples with GraphPad Prism 10.0. Two-way ANOVA analyses were corrected for multiple comparisons. Unless otherwise noted, only statistical comparisons with p<0.05 are shown. *p<0.05, **p<0.01, ***p<0.001, ****p<0.0001, n.s. not significant.

## 3 RESULTS

### 3.1 Adaptive immunity is enhanced in early life and differentially by sex in mouse models of AD

To evaluate the contribution of the microbiome to immune responses in Alzheimer’s disease (AD), we examined immune responses in two well-established mouse models (3xTg and 5xFAD) [9,10] in the presence or absence of the microbiome, utilizing specific pathogen-free (SPF; i.e., laboratory microbiota) and germ-free (GF) mice (**Figure S1**). We analyzed both local (tissue-specific draining lymph nodes) and systemic (spleen) immune responses in male and female mice at ages corresponding to previously reported onset and progression of AD-related pathology: 7, 12, and 15 months for the 3xTg model and 5 and 8 months for the 5xFAD model.

In the superficial and deep cervical lymph nodes (CLNs), which are the primary draining lymph nodes for the lymphatics associated with the brain, head, and neck, we found that CD4^+^ T cell frequencies and numbers were elevated in female 3xTg mice at the earliest age measured (7 months), whether the mice were raised under GF or SPF conditions (**Figure 1A and B**; **Figure S2A and B**). This increase in CD4^+^ T cells in brain-associated lymph nodes occurred before the onset of behavioral deficits previously reported at 12 months of age [21,28]. T cell increases in the CLNs were driven by CD4^+^ (helper) T cells, (**Figure 1A and 1B**) as well as CD8^+^ (cytotoxic) T cells in the deep CLNs (**Figure S2D**). Notably, there were no significant increases in CD4^+^ T cell frequencies in male 3xTg mice at the same age, indicating sexual dimorphism in immune responses in the 3xTg model at this early age (**Figure 1A and B; Figure S2A-D**). This parallels previously described differences in the onset of Aβ plaque accumulation in the brains of 3xTg mice [11,21,29], with female mice showing plaques in the hippocampus and cortex and impaired cognitive function at earlier ages than males [11,30]. We found decreased B cell numbers in the superficial CLNs of GF and SPF female 3xTg mice (**Figure S3A**), indicating differential rates of accumulation or egress between T and B cells in the lymph nodes. No changes were observed in the deep CLNs (**Figure S3B**).

**Figure 1.**
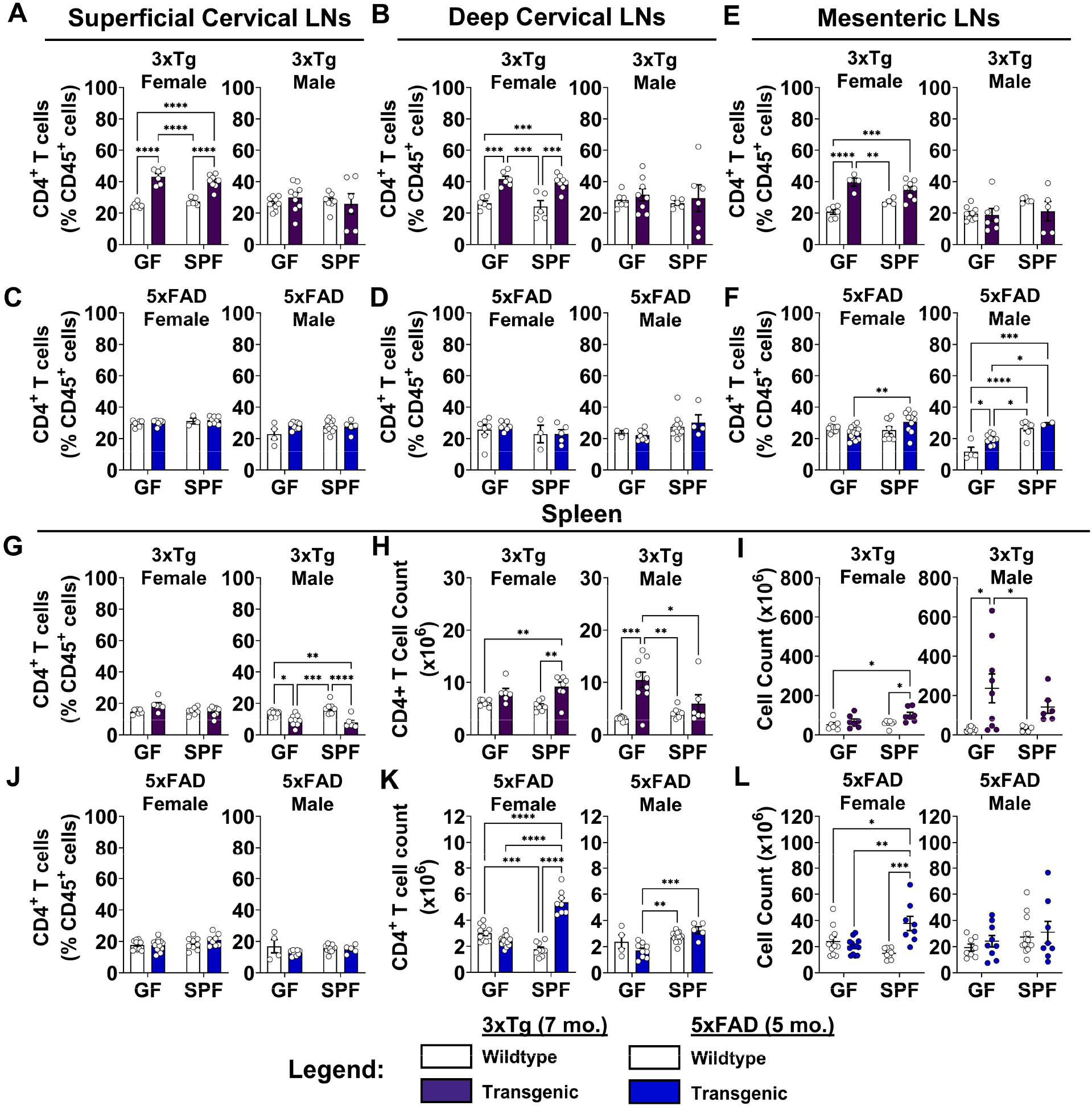
Adaptive immunity is activated early and differentially by sex in mouse models of AD. (A) Quantification of CD4^+^ T cell frequencies in the superficial cervical lymph nodes of female (Left) and male (Right) 3xTg mice compared to wildtype controls in germ-free (GF) and specific pathogen-free (SPF) conditions at 7 months of age. (B) Deep cervical lymph nodes. Data are pooled from 2 independent experiments (n = 4 to 8 per group); two-way ANOVA. (C) Superficial cervical lymph nodes of 5xFAD mice at 5 months of age. (D) Deep cervical lymph nodes. Data are pooled from 2 independent experiments (n = 4 to 10 per group); two-way ANOVA. (E) Mesenteric lymph nodes of 3xTg and (F) 5xFAD mice. Data are pooled from 2 independent experiments (n = 4 to 10 per group); two-way ANOVA. (G) Spleens of 3xTg mice. (H) Quantification of 3xTg spleen CD4^+^ T cell counts. (I) Cell counts in the spleens of 3xTg mice. Data are pooled from 2 independent experiments (n = 6 to 9 per group); two-way ANOVA. (J) Quantification of CD4^+^ T cell frequencies in the spleens of 5xFAD mice. (K) Quantification of 5xFAD spleen CD4^+^ T cell counts. (L) Cell counts in the spleens of 5xFAD mice. Data are pooled from 2 independent experiments (n = 4 to 12 per group); two-way ANOVA. Error bars indicate SEM. *P < 0.05, **P < 0.01, ***P < 0.001, ****P < 0.0001.

In contrast, in the 5xFAD mouse model, we observed no changes in CD4^+^ T cell frequency in the CLNs in female or male mice at the earliest age measured (5 months) (**Figure 1C and D**).

However, decreases in total T cell numbers (**Figure S2E and F**) and B cell numbers (**Figure S3E, F, I, and J**) were observed in the CLNs and intestines, though not all comparisons followed this pattern.

In the mesenteric lymph nodes (MLNs), which drain the small and large intestines, we observed elevated T and B cell numbers in both SPF female and male 3xTg mice at 7 months of age, but not in GF mice, indicating increased adaptive immune response in the intestines that is microbiome dependent (**Figures S2G and S3C**). CD4^+^ T cell proportions in the MLNs increased in both GF and SPF female, but not male, mice (**Figure 1E**). CD8^+^ T cell numbers in the MLNs were unaffected by sex, genotype, or microbiome status (**Figure S2H**).

In female 5xFAD mice at 5 months of age, there was no significant difference in CD4^+^ T cell frequencies from wildtype (WT) mice in the MLNs, but a slight increase was observed between GF and SPF conditions (**Figure 1F**). SPF male 5xFAD mice displayed an increase in CD4^+^ T cell frequencies regardless of genotype (**Figure 1F**). B cell numbers were elevated in female 5xFAD mice under GF conditions, while SPF mice paradoxically showed lower B cell counts (**Figure S3G**).

We next assessed systemic immune responses by examining the spleen. CD4^+^ T cell frequencies were decreased in male, but not female, 3xTg mice under both microbiome conditions (**Figure 1G**). However, total CD4^+^ T cell numbers were elevated in both female and male 3xTg mice, with the greatest increase in GF male mice (**Figure 1H**). These increased CD4^+^ T cell numbers tracked with an increased overall number of cells in the spleen, consistent with previous observations of increased immune cell numbers in the spleen in this model (**Figure 1I**) [31,32]. In female 3xTg mice, spleen size, quantified by cell count, doubled in SPF conditions (**Figure 1I**). In male mice, spleen cell counts increased even more remarkably by 8- and 4-fold in GF and SPF conditions, respectively (**Figure 1I**). The greater increase in spleen cellularity in GF male mice was unexpected and suggests a sex-dependent role for the microbiome in systemic immune responses in 3xTg mice. 5xFAD mice showed no changes in CD4^+^ T cell frequencies and slight increases in B cell frequencies (**Figures 1J** **and S2I**) but, again, significant increases in total CD4^+^ T cell and B cell numbers, an effect which was most pronounced in SPF female mice (**Figure 1K and L; Figure S3H**).

Together, these results show elevated adaptive immune responses both in locally draining lymph nodes (brain and intestine) and systemically (spleen) at the earliest ages examined in both models. In 3xTg mice, CD4^+^ T cell numbers in CLNs were elevated in females, but not males, regardless of microbiome status. Systemic immune responses were modulated by the microbiome and differed by sex, with increased immune cell numbers in GF males. In 5xFAD mice, increases in T and B cell numbers were restricted to the spleen and MLNs, suggesting less pronounced immune responses than in 3xTg mice. These data reveal sexual dimorphism in both immunity and the effects of the microbiome in these mouse models. It is not clear whether the elevated CD4^+^ T cells and B cells play protective or pathogenic roles.

### 3.2 The immune response in 3xTg mice is characterized by increased cytokine responses in males, but attenuated cytokine responses in females

To characterize the quality of T cell responses, we examined cytokine production and profiled effector and regulatory T cells in both the 3xTg and 5xFAD models. The cytokine interleukin (IL)-17A has been implicated in the pathogenesis of neurodegenerative diseases including stroke, multiple sclerosis, AD, and Parkinson’s disease [33]. IL-17A is expressed at barrier and mucosal surfaces, including the skin, gastrointestinal and urogenital tracts, and is induced by specific bacterial taxa of the microbiome that colonize environmentally exposed surfaces of the body [33,34]. The functional role of IL-17A in the CNS is under active investigation, but data suggest that this cytokine promotes neuroinflammation [30,33,35,36]. In 12-month-old male 3xTg mice, we observed elevated IL-17A^+^ CD4^+^ T cells in the brain-associated superficial and deep CLNs (**Figure 2A-D**), despite unchanged overall CD4^+^ T cell frequencies (**Figure S4A-D**). Both IL-17A^+^ T cell frequencies (**Figure 2A-D**), and IL-17A mean fluorescence intensity (MFI; **Figure 2E-H**) — a measurement of average cytokine production per cell — trended higher in SPF compared to GF conditions, consistent with IL-17 induction by the microbiome. Notably, IL-17A MFI in female mice was lower compared to male mice (**Figure 2E-H**), indicating low IL-17A production by T cells in female mice. In peripheral lymphoid tissues outside of the CNS, such as the spleen and MLNs, there were decreased frequencies of IL-17A-producing T cells and lower MFI in both male and female SPF 3xTg compared to SPF wildtype mice (**Figure 2A, B, E, and F**). Overall, female 3xTg mice did not have statistically significant increases in the frequency of cytokine-producing T cells or average cytokine production as determined by MFI (**Figure 2A-H**), despite showing elevated CD4^+^ T cell frequencies (**Figure S4A-D)**. Together, these data indicate that T cell proportions are elevated in 3xTg female mice, but IL-17A production is attenuated, whereas the opposite effect occurs in males, particularly in the brain-associated lymph nodes.

**Figure 2.**
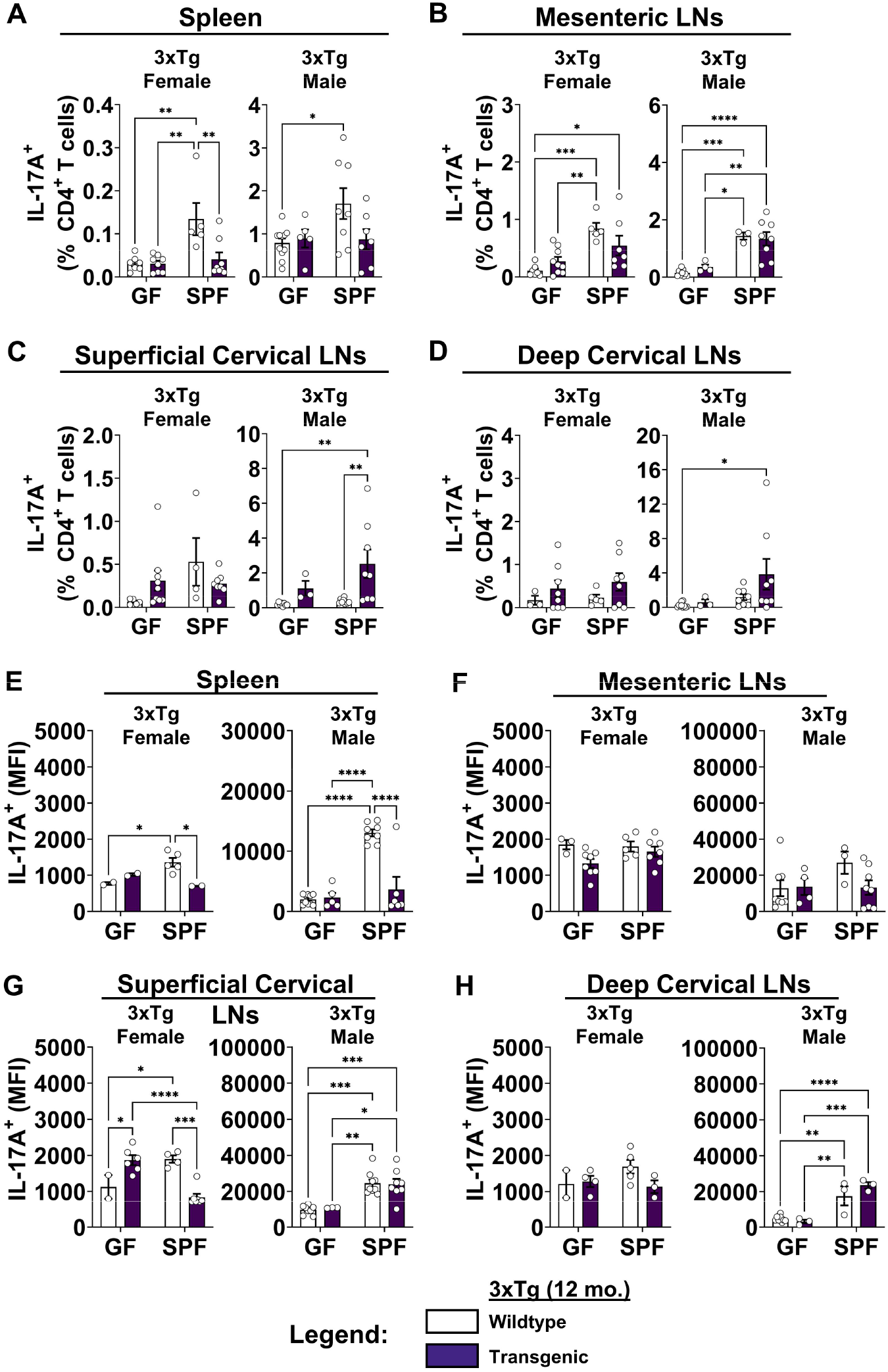
**The immune response in 3xTg mice is characterized by increased cytokine responses in males, but attenuated cytokine responses in females.** (A) Quantification of IL-17A^+^ T cell frequencies in the spleens of female (Left) and male (Right) 3xTg mice compared to wildtype controls in germ-free (GF) and specific pathogen-free (SPF) conditions at 12 months of age. (B) Mesenteric lymph nodes. (C) Superficial cervical lymph nodes. (D) Deep cervical lymph nodes. (E) Quantification of IL-17A mean fluorescence intensity (MFI) expressed by CD4^+^ T cells in the spleen. (F) Mesenteric lymph nodes. (G) Superficial cervical lymph nodes. (H) Deep cervical lymph nodes. Data are pooled from 2 independent experiments (n = 3 to 10 per group); two-way ANOVA. Error bars indicate SEM. *P < 0.05, **P < 0.01, ***P < 0.001, ****P < 0.0001.

Interferon gamma (IFNγ) is a signature cytokine in neuroinflammatory environments [37]. Similar to IL-17A, IFNγ-producing T cells were increased in male, but not female 3xTg mice, particularly in the superficial and deep CLNs (**Figure S5A-D**). IFNγ-producing T cells were elevated in MLNs of male 3xTg mice, although not statistically significantly (**Figure S5B)**. IFNγ MFI increased significantly between GF and SPF conditions in the MLNs and deep CLNs (**Figure S5F and H**) and was elevated in the spleen and superficial CLNs (**Figure S5E and G**) of wildtype female mice. This was not observed in male mice, except in the spleen, indicating a sex-dependent microbiome effect on IFNγ production in the lymph nodes (**Figure S5E-H**).

Notably, in SPF female mice, IFNγ MFI decreased comparing wildtype to transgenic mice, suggesting a sex-dependent regulation of cytokine production (**Figure S5E-H**). In the spleen, we observed decreased frequencies of IFNγ -producing T cells and lower MFI in both male and female SPF 3xTg compared to SPF wildtype mice (**Figure S5A and E**). These results are congruent with the IL-17A results and show an increase in IFNγ^+^ T cell frequencies in the CLNs of male 3xTg mice, but a decrease in IFNγ^+^ T cell frequencies and cytokine production in the spleens of female and male 3xTg SPF mice, highlighting differences between local and systemic immune responses.

In inflamed environments, regulatory T cell numbers increase with expanding cytokine-producing effector T cells and act to limit immune responses [38,39]. A major subset of CD4^+^ regulatory T cells is marked by the forkhead box transcription factor P3 (Foxp3). Foxp3^+^ T cell frequencies were not significantly changed between transgenic and WT mice but were reduced in SPF compared to GF 3xTg mice in most tissues and both sexes (**Figure S6A-D**). This indicates a potentially unbalanced regulatory response by Foxp3^+^ T cells in 3xTg mice.

### 3.3 The immune response in 5xFAD mice is characterized by increased IL-17A-producing T cells

Consistent with the role of the microbiome in promoting IL-17A production [33,34], we found that IL-17A^+^ T cells and IL-17A MFI in the intestines of 5xFAD mice were increased by the presence of the microbiome (i.e., SPF conditions) (**Figure 3A, B, F and G)**.

**Figure 3.**
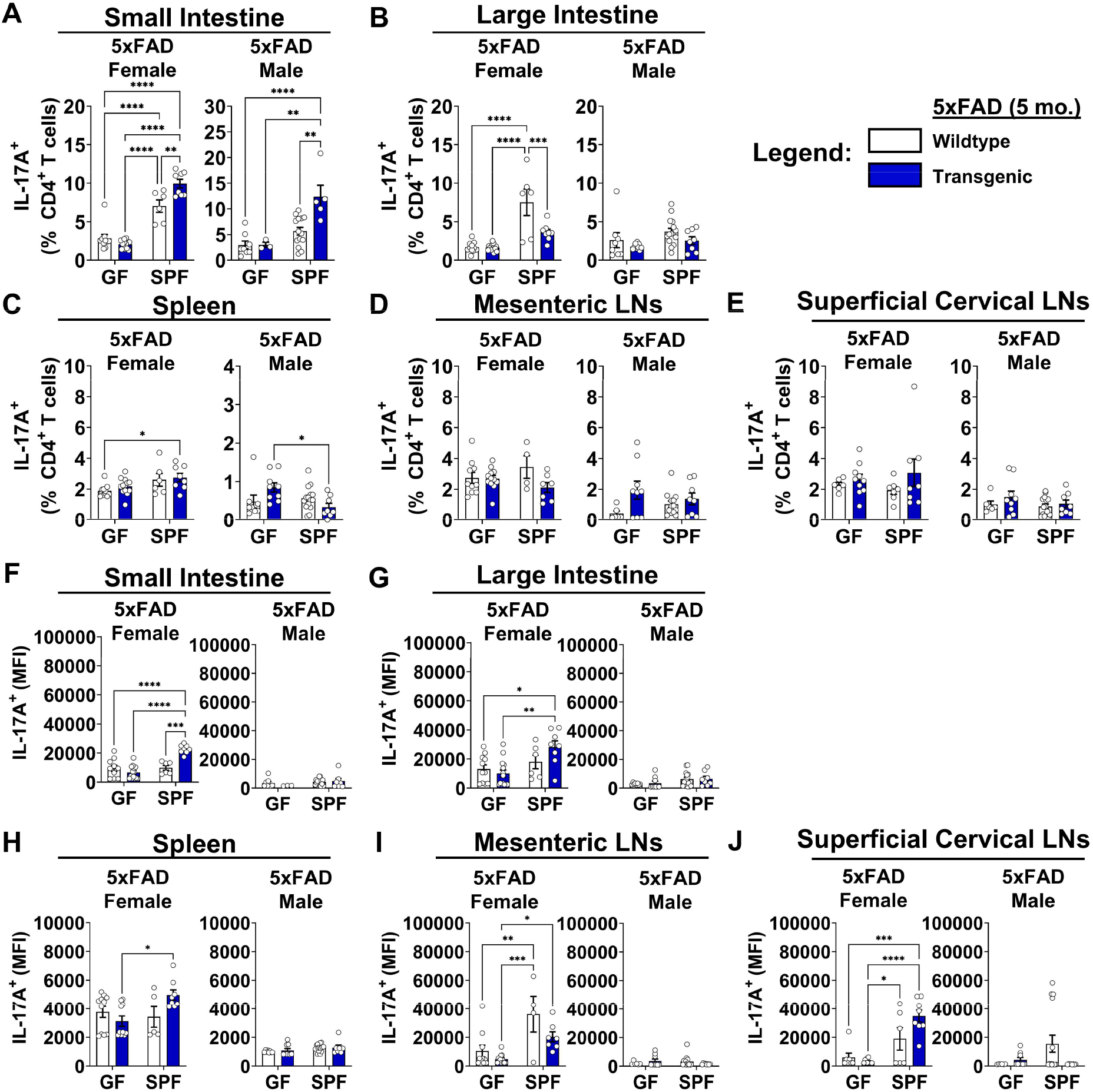
**The immune response in 5xFAD mice is characterized by increased IL-17A-producing T cells.** (A) Quantification of IL-17A^+^ T cell frequencies in the small intestines of female (Left) and male (Right) 5xFAD mice compared to wildtype controls in germ-free (GF) and specific pathogen-free (SPF) conditions at 5 months of age. (B) Large intestine. (C) Spleen. (D) Mesenteric lymph nodes. (E) Superficial cervical lymph nodes. (F) Quantification of IL-17A mean fluorescence intensity (MFI) expressed by CD4^+^ T cells in the small intestine. (G) Large intestine. (H) Spleen. (I) Mesenteric lymph nodes. (J) Superficial cervical lymph nodes. Data are pooled from 2 independent experiments (n = 4 to 14 per group); two-way ANOVA. Error bars indicate SEM. *P < 0.05, **P < 0.01, ***P < 0.001, ****P < 0.0001.

Importantly, and unexpectedly, we found IL-17A-producing CD4^+^ T cells were further increased in the small intestines of SPF 5xFAD transgenic female and male mice compared to wildtype controls at 5 months of age (**Figure 3A**). Previous reports indicated reduced IL-17A production in the gut-associated immune cell populations of 5xFAD transgenic mice, specifically in the mesenteric lymph nodes and Peyer’s patches [40]. Indeed, we observed lower IL-17A^+^ T cell frequencies in the large intestine, spleen, and MLNs of SPF transgenic mice (**Figure 3B-D**) and lower IL-17A MFI in the MLNs (**Figure 3H and I**). We found no changes in IL-17A^+^ T cell frequencies or counts in the superficial CLNs (**Figures 3E** **and S7A**), however, we observed an increase in IL-17A MFI in SPF transgenic female mice (**Figure 3J**). The increase in the IL-17A^+^ T cell populations in the small intestine tissue (lamina propria) (**Figure 3A**) vs. the large intestine and MLNs (**Figure 3B and D**) indicates the environment of the small intestine is specifically conducive to either IL-17A^+^ T cell differentiation, proliferation, and/or retention. In accordance with the observed increase in spleen cell counts (see **Figure 1L**), the total number of IL-17A^+^ T cells dramatically increased in the spleens of SPF female 5xFAD mice (**Figure S7B**). We observed a moderate increase in IL-17A^+^ T cell numbers in the MLNs of GF male transgenic mice (**Figure S7C**). The observed increase of IL-17A^+^ T cells in the 5xFAD transgenic mice suggests increased cytokine production that is modulated by gut bacteria [35,36], and may be induced by an alteration in microbiome localization or composition in the intestines, or both.

We observed largely unchanged IFNγ-producing T cell frequencies in 5xFAD compared to wildtype control mice in most tissues (**Figure S8A-E**). However, IFNγ^+^ T cell frequencies increased in the spleen and superficial CLNs of female SPF 5xFAD mice (**Figure S8C and E**), consistent with a previous study that noted increased transcription of the IFNγ-induced chemokine Cxcl10 in the brains of female 5xFAD mice at 5 months of age [41]. IFNγ MFI remained unchanged in most tissues (**Figure S8F-G**), apart from an increase in the MLNs and superficial CLNs of SPF female mice (**Figure S8I and J**).

We examined two additional cytokines associated with activated immune responses, granulocyte-macrophage colony-stimulating factor (GM-CSF) and interleukin (IL)-4. GM-CSF produced by lymphocytes activates myeloid cells, such as macrophages and microglia, and participates in the communication between the two subsets of immune cells [37]. IL-4 is a key cytokine in the induction of Type 2 helper T cells (Th2) that mediate allergy, asthma, and wound repair, and in addition, acts as an anti-inflammatory cytokine that antagonizes Type 1 (IFNγ) immune responses [42]. Although only statistically significant in the small intestine, we observed an overall trend of increased GM-CSF^+^ T cell frequencies in GF 5xFAD transgenic mice compared to GF wildtype controls, but decreased frequencies in SPF mice (**Figure S9A-E**). GM-CSF MFI was significantly increased in the small intestine, MLNs, and superficial CLNs of SPF compared to GF female mice, but not in other tissues or male mice (**Figure S9F-J**). We generally observed low and highly variable frequencies of IL-4^+^ T cells in all tissues examined (**Figure S10A-E**).

Remarkably, IL-4 MFI was increased in all tissues examined in SPF transgenic compared to SPF wildtype female and male mice, possibly indicating a regulatory response to Type 1 immunity (**Figure S10F-J)**.

We observed increased Foxp3^+^ regulatory T cell frequencies and counts in the spleens, MLNs and deep CLNs of SPF female, but not male, 5xFAD mice (**Figure S11A, B, and D**). Male mice showed lower proportions of Foxp3^+^ T cells in the MLNs, superficial and deep CLNs, and large intestines, regardless of microbiome status (**Figure S11B-E**). Together, these data indicate increased immune responses that are balanced by the induction of regulatory T cells in female, but not male, 5xFAD mice.

### 3.4 Longitudinal immune responses differ between 3xTg mice and 5xFAD mice

The most significant risk factor for AD is aging [1]. To determine how immune responses progressively change in mouse models, we examined mice at multiple ages. We found that systemic immune responses increased with age in 3xTg mice, especially in SPF mice, with increased T and B cell numbers in the spleen. However, local immune responses in the lymph nodes remained the same or declined with age. The CD4^+^ T cell counts that we observed at 7 months of age in SPF female 3xTg animals remained elevated in the spleens and MLNs but declined by half in the CLNs from 7 to 15 months of age (**Figure 4A-D)**. In contrast to SPF female mice, CD4^+^ T cell numbers remained the same or declined in GF female mice (**Figure S12A-D**). Both GF and SPF male mice showed progressive increases in CD4^+^ T cells in the spleen, with minimal change in other tissues as animals aged (**Figure S12E-L**). These data indicate that changes in systemic spleen CD4^+^ T cell responses with age are influenced by microbiome status to a greater extent in female than male mice. B cell numbers mirrored CD4^+^ T cell numbers in female (**Figure 4E-H, S13A-D**) and male (**S13E-L**) 3xTg mice.

**Figure 4.**
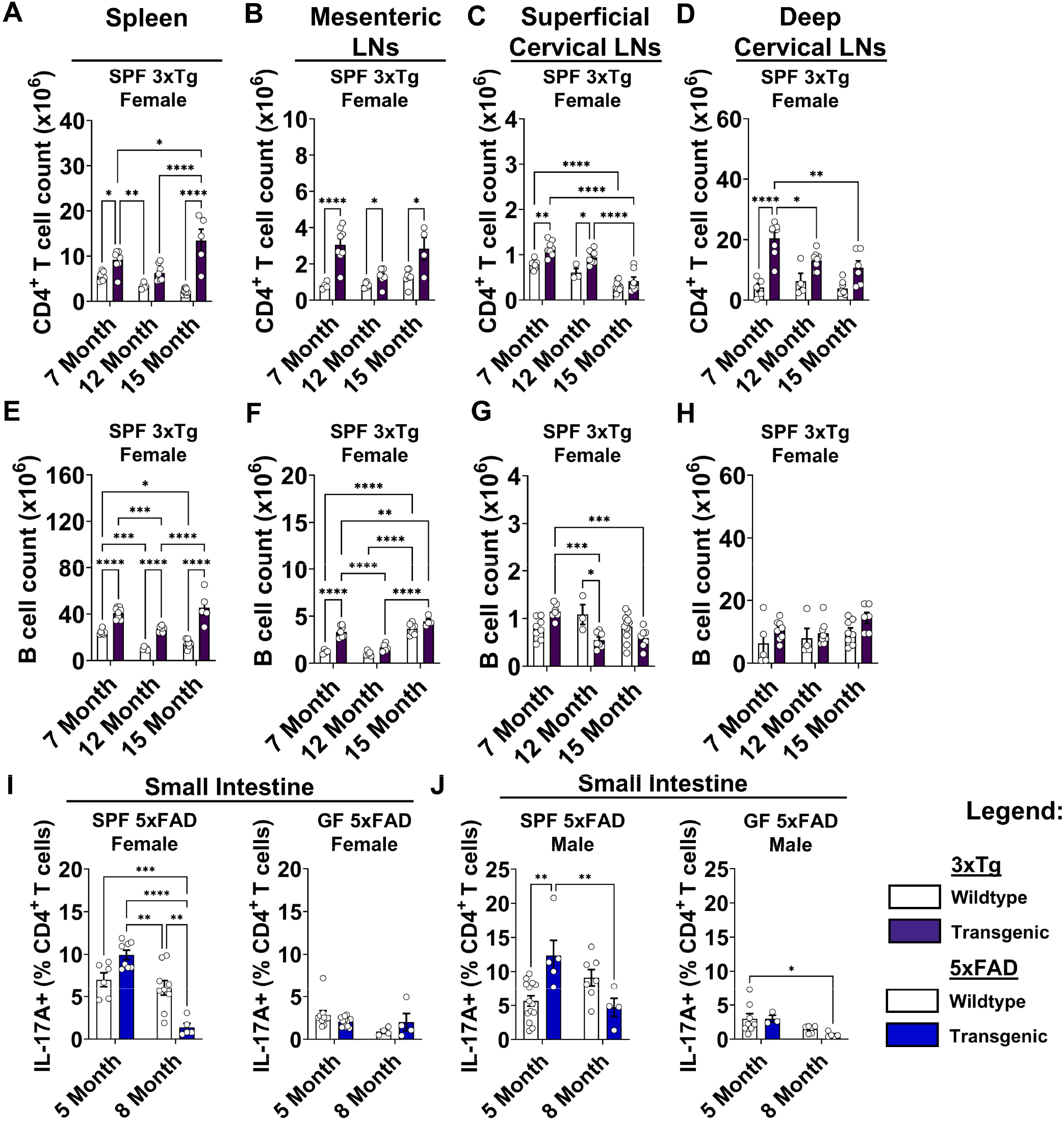
Longitudinal immune responses differ between 3xTg mice and 5xFAD mice. (A) Quantification of CD4^+^ T cell counts in the spleens of female 3xTg mice compared to wildtype controls in specific pathogen-free (SPF) conditions at 7-, 12-, and 15-month ages. (B) Mesenteric lymph nodes. (C) Superficial cervical lymph nodes. (D) Deep cervical lymph nodes. Data are pooled from 5 independent experiments (n = 3 to 8 per group); two-way ANOVA. (E) Quantification of B cell counts in the spleens of female 3xTg mice. (F) Mesenteric lymph nodes. (G) Superficial cervical lymph nodes. (H) Deep cervical lymph nodes. Data are pooled from 5 independent experiments (n = 3 to 11 per group); two-way ANOVA. For clarity, only significant comparisons of the same mouse group between ages or between transgenic and control mice at the same age are shown. (I) Quantification of IL-17A^+^ T cell frequencies in the small intestines of female 5xFAD mice compared to wildtype controls in SPF (Left) and GF (Right) conditions at 5- and 8-month ages. (J) Small intestines of male 5xFAD mice. Data are pooled from 5 independent experiments (n = 4 to 14 per group); two-way ANOVA. Error bars indicate SEM. *P < 0.05, **P < 0.01, ***P < 0.001, ****P < 0.0001.

In SPF 5xFAD mice, IL-17A^+^ T cells were attenuated in the small intestine from 5 to 8 months in both female and male mice (**Figure 4I and J**). However, B cell numbers remained elevated in the spleens of SPF female, but not SPF male, mice from 5 to 8 months of age, indicating sustained systemic B cells in SPF but not GF mice (**Figure S14A and B**). In the MLNs and CLNs, B cell numbers largely remained the same under SPF and GF conditions, with some variability (**Figure S14D, F, and H**), and GF male mice showed a decline in B cell counts with age in the MLNs (**Figure S14D**). We found an age-dependent increase in B cells that was independent of disease in the superficial and deep CLNs and MLNs of SPF female mice (**Figure S14C, E, and G**). The differences in immune responses between 3xTg and 5xFAD mice as animals age, especially in GF conditions, suggest differing mechanisms of immune activation and resolution. Overall, these data reveal that increased T and B cell numbers and CD4^+^ T cell cytokine production develop early in some models of AD, are largely microbiome-dependent, and can be durable or can resolve as animals age depending on variables such as sex and microbiome status.

### 3.5 Both 3xTg and 5xFAD female mice show improved learning and cognition in the absence of a microbiome

We next sought to determine whether differences in immune response correlate with cognitive outcomes. It has been previously reported that AD mouse models display reduced amyloid plaques and improved cognitive function under GF conditions or after antibiotic treatment, compared to mice with an intact microbiome [3–5]. However, reported behavioral outcomes in these two preclinical models have varied widely with regard to presence of a phenotype, age, behavioral paradigm tested, and examination of the influence of the gut microbiota [28,29]. To obtain a broad view of cognitive performance in both models across aging, SPF and GF mice were evaluated via the modified Barnes maze, Y maze, and novel object recognition tests.

Overall, phenotypes were subtle, consistent with several previous studies [28,29,43]. However, we observed the most robust differences in cognitive function between transgenic and wildtype mice in the modified Barnes maze, with differences in both training learning and test performance (**Figure 5** **and S15**).

**Figure 5.**
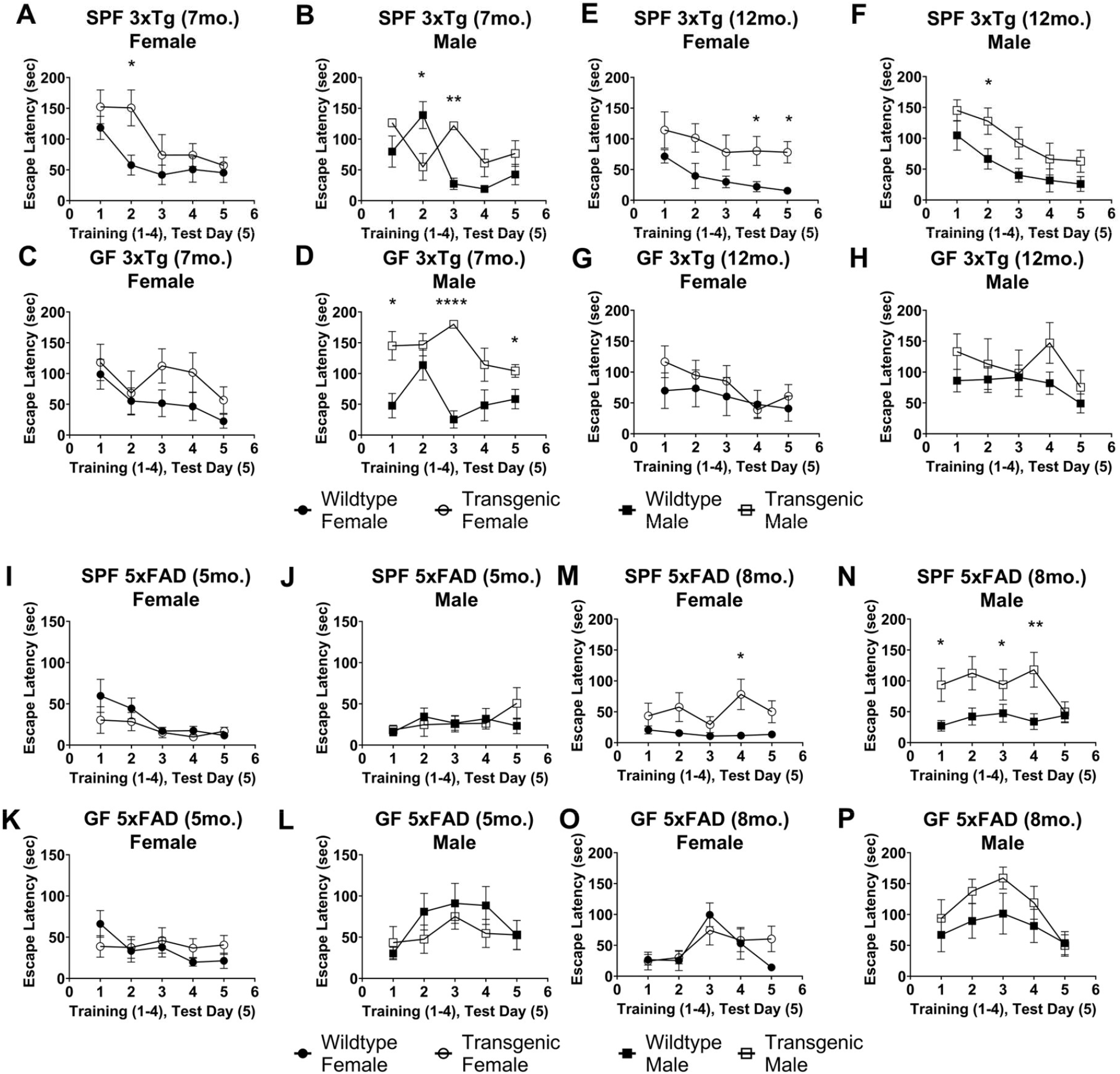
**Both 3xTg and 5xFAD mice show improved learning and cognition in the absence of a microbiome.** (A) Training (1-4) and test (5) day mean escape latency in Barnes Maze for 7-month-old female 3xTg mice compared to wildtype controls in specific pathogen-free (SPF) conditions. (B) Male mice in SPF conditions. (C) Female mice in germ-free (GF) conditions. (D) Male mice in GF conditions. Data are pooled from 2 independent experiments (n = 6 to 11 per group); Mann-Whitney U test comparing transgenic to wildtype mice. (E) 12-month-old female 3xTg mice in SPF conditions. (F) Male mice in SPF conditions. (G) Female mice in GF conditions. (H) Male mice in GF conditions. Data are pooled from 2 independent experiments (n = 5 to 11 per group). (I) 5-month-old female 5xFAD mice in SPF conditions. (J) Male mice in SPF conditions. (K) Female mice in GF conditions. (L) Male mice in GF conditions. Data are pooled from 2 independent experiments (n = 8 to 14 per group); Mann-Whitney U test comparing transgenic to wildtype mice. (M) 8-month-old female 5xFAD mice in SPF conditions. (N) Male mice in SPF conditions. (O) Female mice in GF conditions. (P) Male mice in GF conditions. Data are pooled from 2 independent experiments (n = 5 to 13 per group); Mann-Whitney U test comparing transgenic to wildtype mice. Error bars indicate SEM. *P < 0.05, **P < 0.01, ****P < 0.0001.

Cognitive deficits, as measured by increased latencies to escape the maze, were more apparent in SPF mice compared to their GF counterparts, implicating the microbiome in this behavioral outcome. We observed cognitive deficits in both sexes in SPF 3xTg mice at 7 and 12 months of age (**Figure 5A, B, E, and F**) and SPF 5xFAD mice at 8 months of age (**Figure 5M and N**).

Female mice showed improved performance under GF conditions (**Figure 5C, G, K, and O**), whereas 7-month-old GF 3xTg males maintained cognitive deficits (**Figure 5D**). In addition to comparing wildtype vs. transgenic mice, we also assessed Barnes maze performance between GF and SPF conditions (**Figure S15A-P**). As already noted, 7-month-old GF 3xTg males showed poorer performance than their SPF counterparts (**Figure S15D**). In the 5xFAD experiment, wildtype males showed poorer performance in GF conditions, but the differences were only statistically significant at 5 months of age (**Figure S15J and N**).

Spatial memory testing with the Y maze did not show significant cognitive decline in male or female 3xTg mice at 7 or 12 months of age in either GF or SPF conditions, with the exception of 7-month-old GF 3xTg males (**Figure S16A-C**). Seven and 12-month-old SPF female 3xTg mice performed better than wildtype mice (**Figure S16A and B**), consistent with a previous report [43]. 5xFAD mice showed no deficits compared to wildtype mice in Y maze at 5 or 8 months of age (**Figure S16D and E**). We did not observe any deficits in novel object recognition in male or female 3xTg or 5xFAD mice (**Figure S17**), except for SPF 3xTg females at 7 months of age (**Figure S17A**) and GF 5xFAD males at 8 months of age (**Figure S17E**). Together, these data highlight deficits in spatial and working memory in the 3xTg and 5xFAD mouse models, with negligible differences in recognition memory. Our results support the notion that age, immune state, sex, and the microbiome impact outcomes of cognitive performance, with the most significant effects by gut bacteria being on the interaction between sex and immune responses.

## 4 DISCUSSION

The microbiome is an important contributor to immunity, and has recently been implicated in neurodegenerative disorders such as AD [44–46]. Whether and how gut bacteria actively contribute to AD pathophysiology remains an open question. In this study, we examined immune and cognitive profiles in two transgenic mouse models of AD in the presence or absence of gut bacteria to understand how the microbiome shapes immune responses associated with AD. We found that the microbiome marginally influences immune cell proportions in the brain-draining lymph nodes in both mouse models, but strongly modulates systemic immune responses, the cytokine response in local lymphoid tissues, and cognitive deficits. Notably, the immune state in these mice was modulated by sex, and this sex effect differed between local and systemic lymphoid tissues and between mice with an intact and absent microbiome.

In the 3xTg and 5xFAD mouse models, we found two contrasting immune environments: (1) in 3xTg mice, increased T and B cell numbers and cytokine production were evident throughout secondary lymphoid tissues, including those associated with the intestine and brain, as well as systemically; (2) in 5xFAD mice, changes in immune cell numbers and cytokine production were more limited and, while somewhat systemic, largely localized to intestinal tissue and brain-draining lymph nodes. In both models, responses were driven by activated adaptive immunity, with elevated CD4^+^ T cells and B cells. However, as evident from the grossly elevated spleen cell counts in 3xTg mice compared to the more mildly elevated spleen cell counts of 5xFAD mice, 3xTg mice had a more pronounced immune response. The most significant genetic difference between the models is expression of the tau protein, in addition to amyloid precursor protein and presenilin, in 3xTg mice, which results in tau aggregation (tauopathy) as well as Aβ deposition [11]. Ectopic expression of human tau in mice has been shown to increase adaptive immune responses and brain atrophy in another AD model and may be a factor in the widespread immune responses we observed in 3xTg mice [47].

Previous studies have shown that adaptive immunity participates in both protective and pathogenic processes in AD mouse models [48]. Complete abrogation of T and B cell responses in the 5xFAD model by genetic knockout of *Rag2 and Il2rg* results in increased amyloid aggregation and decreased brain expression of genes related to immunoglobulin antibody production and associated signaling, indicating a protective role for the adaptive immune system [14]. Paradoxically, the loss of B cells alone results in reduced amyloid aggregation and improved cognitive function in 3xTg mice [15]. Studies indicate that B cell-produced autoantibodies are present in both 3xTg and 5xFAD mouse models [31,49]. Chronic inflammatory states may contribute to pathogenic mechanisms that promote cellular stress, death, and the progression of AD. T and B cells, among other immune cells, have been shown to promote chronic inflammation through antibody production, activation of additional immune cells, and inflammatory cytokine production [15,48,50,51]. We observed that adaptive immune responses were largely unchanged with age in 3xTg mice, in contrast to the attenuation of immune responses in 5xFAD mice. This indicates a chronic elevation of immune responses in 3xTg mice, which may contribute to more severe pathological outcomes.

Here, we find that immune cell proportions in both the 3xTg and 5xFAD models are influenced by sex, with female mice showing earlier activation of adaptive immunity. This corresponds with a previously reported early rise in Aβ in the brains of female 3xTg and 5xFAD mice [11,52]. The cause of this accelerated pathology in female mice is not well understood. Previous studies suggested that pathology in the 3xTg and 5xFAD models is influenced by sex hormones, as alterations of sex hormones during early development and adulthood in 3xTg mice, or modulation of estrogen receptor activity in 5xFAD mice, can modify Aβ accumulation [21–24]. Notably, an early study in humans identified that estrogen-based hormone therapy reduces the risk of AD in women [53].

We observed elevated T and B cell counts and cytokine production in the draining lymph nodes of both the brain and intestine, indicating that immune responses may be activated and/or maintained by both brain and peripheral amyloid or tau pathology. There is significantly more (25x) Aβ peptide in the circulation of 3xTg compared to 5xFAD mice, which might contribute to the increased immune cell numbers in the spleen and liver of 3xTg mice observed in this study and others [31,32,54,55]. The liver, kidneys, spleen, and small intestine are important sites for the clearance of circulating Aβ and may act as hubs for activation of systemic immunity [56,57]. Previous work demonstrated reduced amyloid concentration in the brains of GF or antibiotic-treated mice [3–5]; therefore, our expectation was to find reduced systemic inflammation.

Remarkably, while 3xTg female mice showed attenuated systemic immune responses under GF conditions, 3xTg male mice had elevated T and B cell numbers. A previous study noted that male 3xTg mice develop a more robust autoimmune-like response, with serum antibodies against cell nuclei [58]. In 5xFAD mice, we observed reduced T and B cell frequencies in GF compared to SPF conditions. These findings suggest a sex- and microbiome-mediated effect on systemic immune responses which requires further study.

Increased immune cell numbers and CD4^+^ T cell cytokine production corresponded with cognitive deficits in 3xTg and 5xFAD mice, an effect most pronounced in 3xTg female mice at early ages and in SPF conditions. Of note, GF 3xTg male mice had significant cognitive deficits, which corresponded with an increased systemic inflammatory response, suggesting that severe inflammatory responses may be associated with cognitive function, although the mechanism is not clear. 5xFAD mice indicated cognitive deficits at only the last time point measured (8 months), with deficits more pronounced in SPF than GF conditions.

Limitations of this work include scope of the immune response survey, restricted mainly to secondary lymphoid tissues and the adaptive immune response, and variability in the results of cognitive testing. Discrepancies in reported cognitive deficits exist across studies [28,30], perhaps due to technical differences between testing conditions, different microbiomes between animal facilities, and other variables. In future studies, metagenomic sequencing of the microbiome may help determine causes of variability. For immune response evaluation, we chose secondary lymphoid tissues because they represent hubs of immune activation. In future studies, local tissue responses may provide additional insight into the mechanisms of immune activation and resolution. Amyloid beta (Aβ) and tau analysis were not performed in this study to assess brain pathology differences between groups, which is a feature of these animal models.

In conclusion, we found that the microbiome influences both immune and cognitive outcomes in a sex-dependent manner in two mouse models of AD. Increased adaptive immune responses were observed in both female and male mice and were attenuated in germ-free conditions.

Unexpectedly, GF male 3xTg mice maintained elevated adaptive immune and cytokine responses that correlated with cognitive deficits, indicating a protective role for the microbiome in male mice in this model. These data demonstrate a sex and microbiome interaction in the immune response and cognitive performance of AD mouse models. Future studies may focus on microbiome-influenced mechanisms that intersect with sex hormone signaling and immune responses to determine their effects on AD-like outcomes in mice, and perhaps in humans.

## Supporting information

Supplementary Representative Flow Cytometry Plots

## ACKNOWLEDGMENTS

We thank the Mazmanian laboratory for helpful suggestions on this work and Catherine Oikonomou for assistance with manuscript editing. We thank the Caltech Flow Cytometry and Cell Sorting Facility for flow cytometry services and assistance.

## CONSENT STATEMENT

Informed consent was not applicable to this study.

---

**Supplementary Figure 1.**
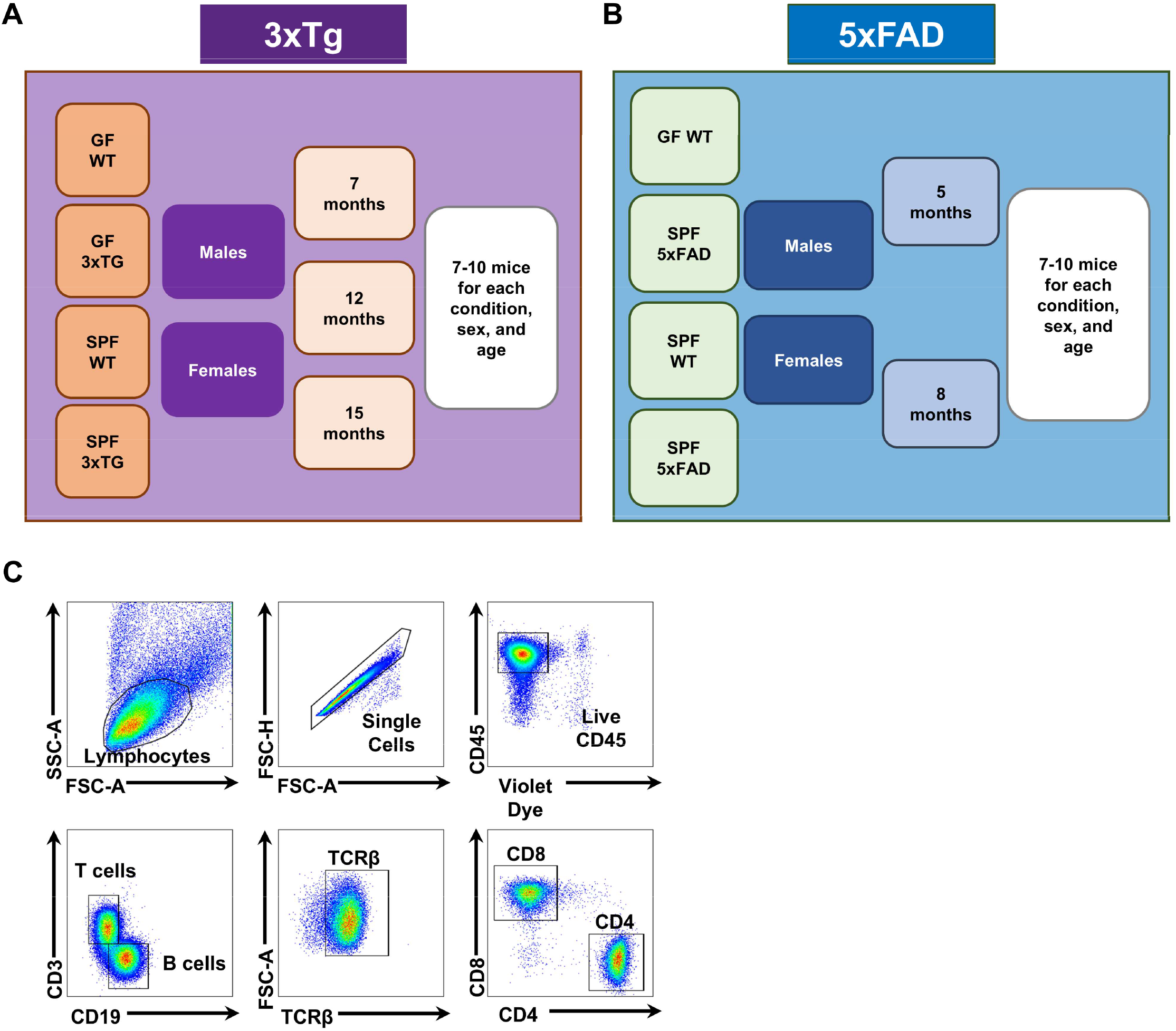
**Experimental design and flow cytometry gating.** (A) Illustration of experimental design for genotypes, housing conditions, ages, and sexes for 3xTg and (B) 5xFAD mice. (C) Flow cytometry gating strategy for T and B cell analysis.

**Supplementary Figure 2.**
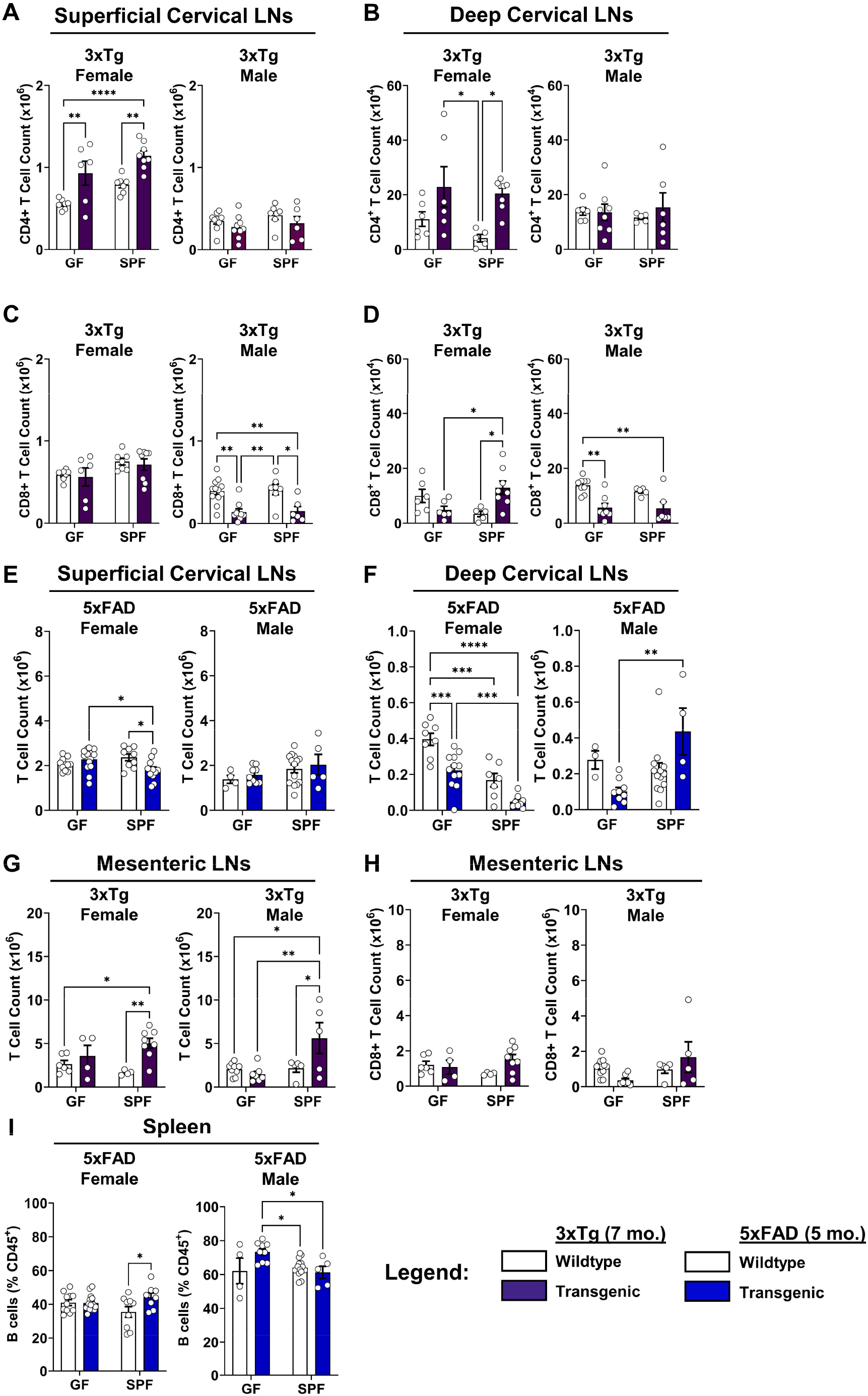
T and B cell immune responses in the 3xTg and 5xFAD models. (A) Quantification of CD4^+^ T cell counts in the superficial cervical lymph nodes of female (Left) and male (Right) 3xTg mice compared to wildtype controls in germ-free (GF) and specific pathogen-free (SPF) conditions at 7 months of age. (B) Deep cervical lymph nodes. (C) Quantification of CD8^+^ T cell counts in the superficial cervical lymph nodes of female (Left) and male (Right) 3xTg mice at 7 months of age. (D) Deep cervical lymph nodes. Data are pooled from 2 independent experiments (n = 5 to 14 per group); two-way ANOVA. (E) Quantification of T cell counts in the superficial cervical lymph nodes of female (Left) and male (Right) 5xFAD mice at 5 months age. (F) Deep cervical lymph nodes. Data are pooled from 2 independent experiments (n = 3 to 14 per group); two-way ANOVA. (G) Quantification of T cell counts in the mesenteric lymph nodes of female (Left) and male (Right) 3xTg mice at 7 months of age. (H) Quantification of CD8^+^ T cell counts. Data are pooled from 2 independent experiments (n = 4 to 11 per group); two-way ANOVA. (I) Quantification of B cell frequencies in the spleens of female (Right) and male (Left) 5xFAD mice at 5 months of age. Data are pooled from 2 independent experiments (n = 4 to 14 per group); two-way ANOVA. Error bars indicate SEM. *P < 0.05, **P < 0.01, ***P < 0.001, ****P < 0.0001.

**Supplementary Figure 3.**
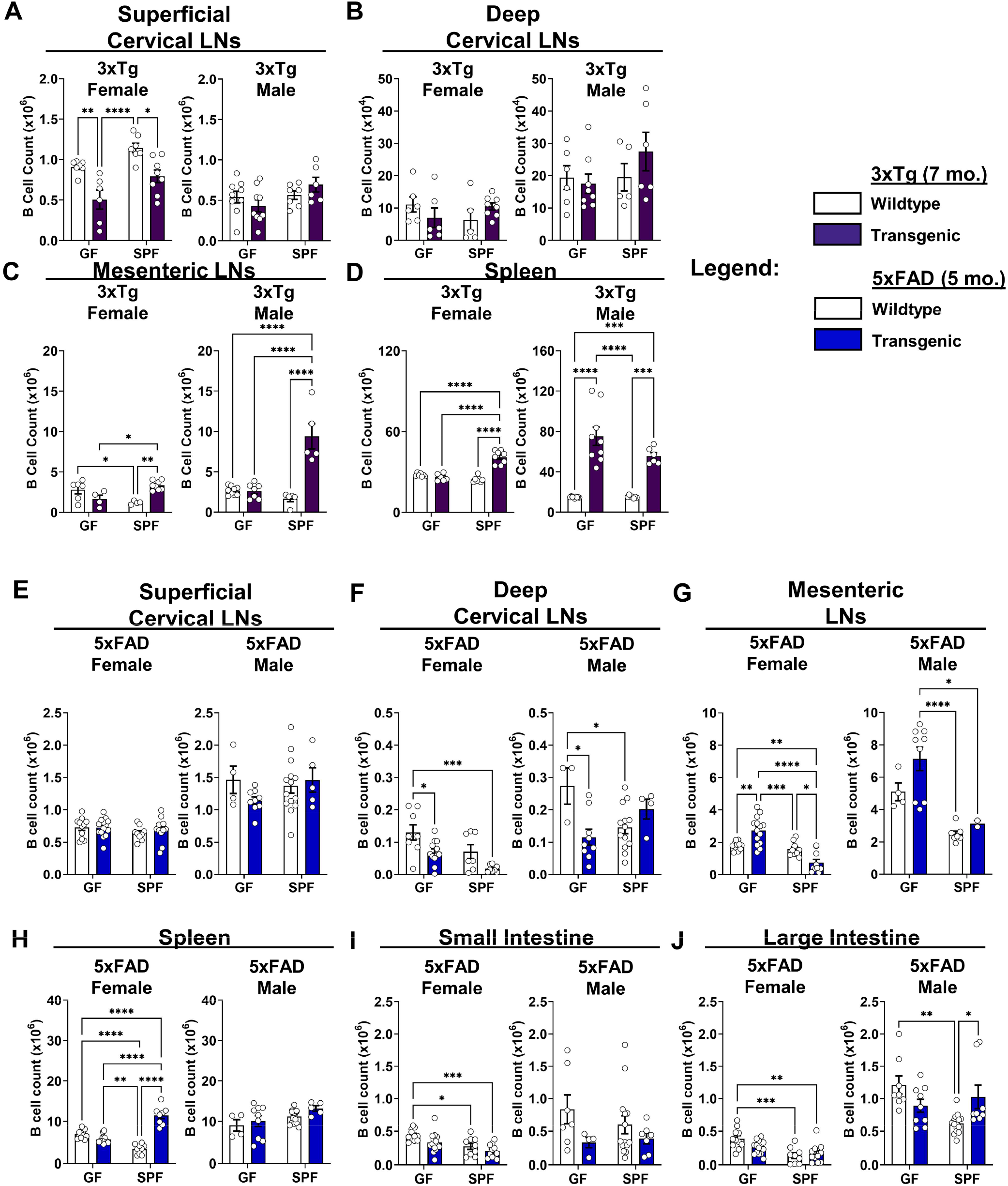
B cell counts in 7-month-old 3xTg and 5-month-old 5xFAD mice. (A) Quantification of B cell counts in the superficial cervical lymph nodes of female (Left) and male (Right) 3xTg mice compared to wildtype controls in germ-free (GF) and specific pathogen-free (SPF) conditions at 7 months of age. (B) Deep cervical lymph nodes. (C) Mesenteric lymph nodes. (D) Spleen. Data are pooled from 2 independent experiments (n = 4 to 14 per group); two-way ANOVA. (E) Quantification of B cell counts in the superficial cervical lymph nodes of female (Left) and male (Right) 5xFAD mice compared to wildtype controls in germ-free (GF) and specific pathogen-free (SPF) conditions at 5 months of age. (F) Deep cervical lymph nodes. (G) Mesenteric lymph nodes. (H) Spleen. (I) Small intestine. (J) Large intestine. Data are pooled from 2 independent experiments (n = 2 to 15 per group); two-way ANOVA. Error bars indicate SEM. *P < 0.05, **P < 0.01, ***P < 0.001, ****P < 0.0001.

**Supplementary Figure 4.**
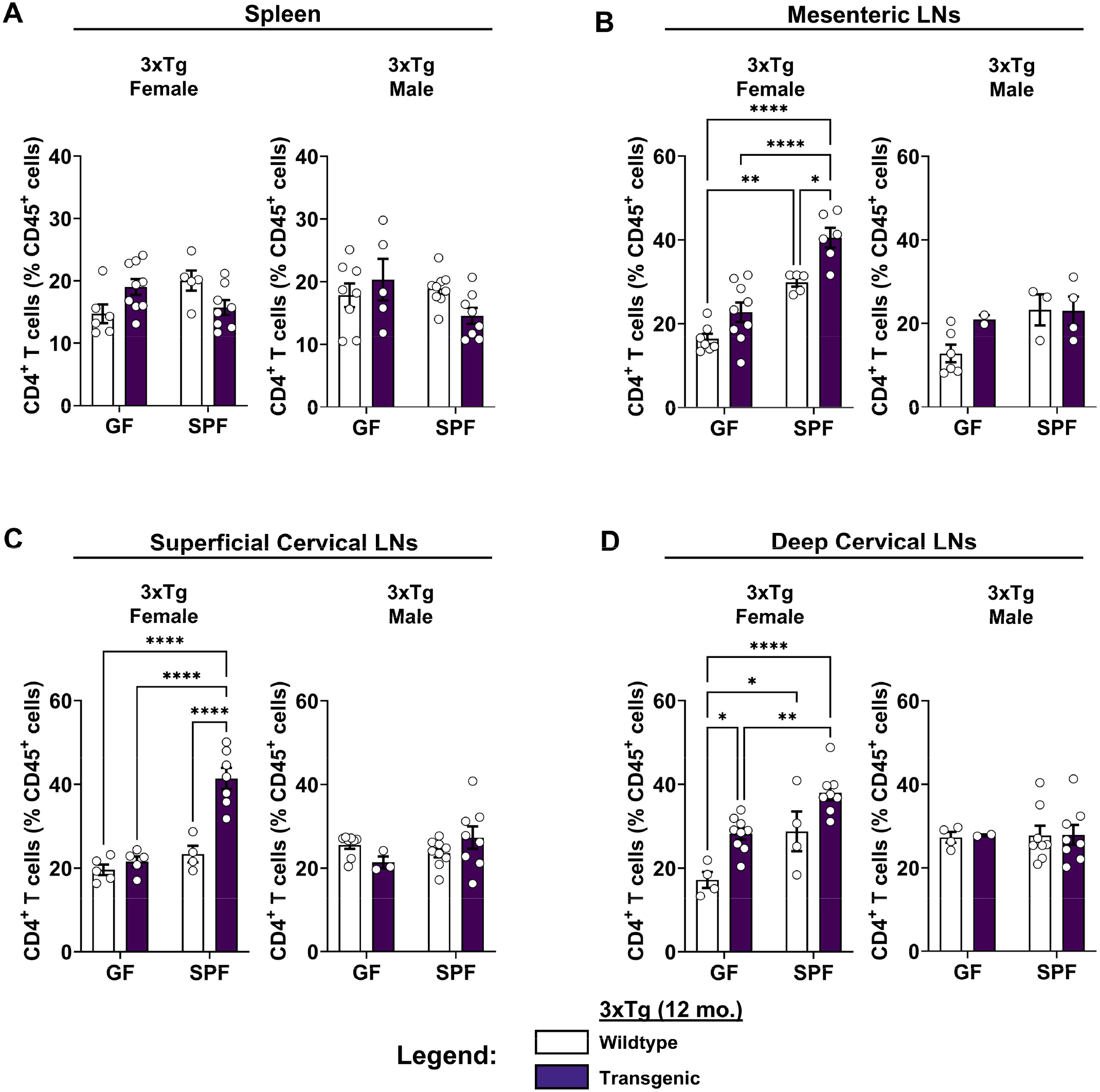
Quantification of CD4^+^ T cell frequencies in 12-month-old 3xTg mice. (A) Quantification of CD4^+^ T cell frequencies in the spleens of female (Left) and male (Right) 3xTg mice compared to wildtype controls in germ-free (GF) and specific pathogen-free (SPF) conditions at 12 months of age. (B) Mesenteric lymph nodes. (C) Superficial cervical lymph nodes. (D) Deep cervical lymph nodes. Data are pooled from 2 independent experiments (n = 2 to 9 per group); two-way ANOVA. Error bars indicate SEM. *P < 0.05, **P < 0.01, ***P < 0.001, ****P < 0.0001.

**Supplementary Figure 5.**
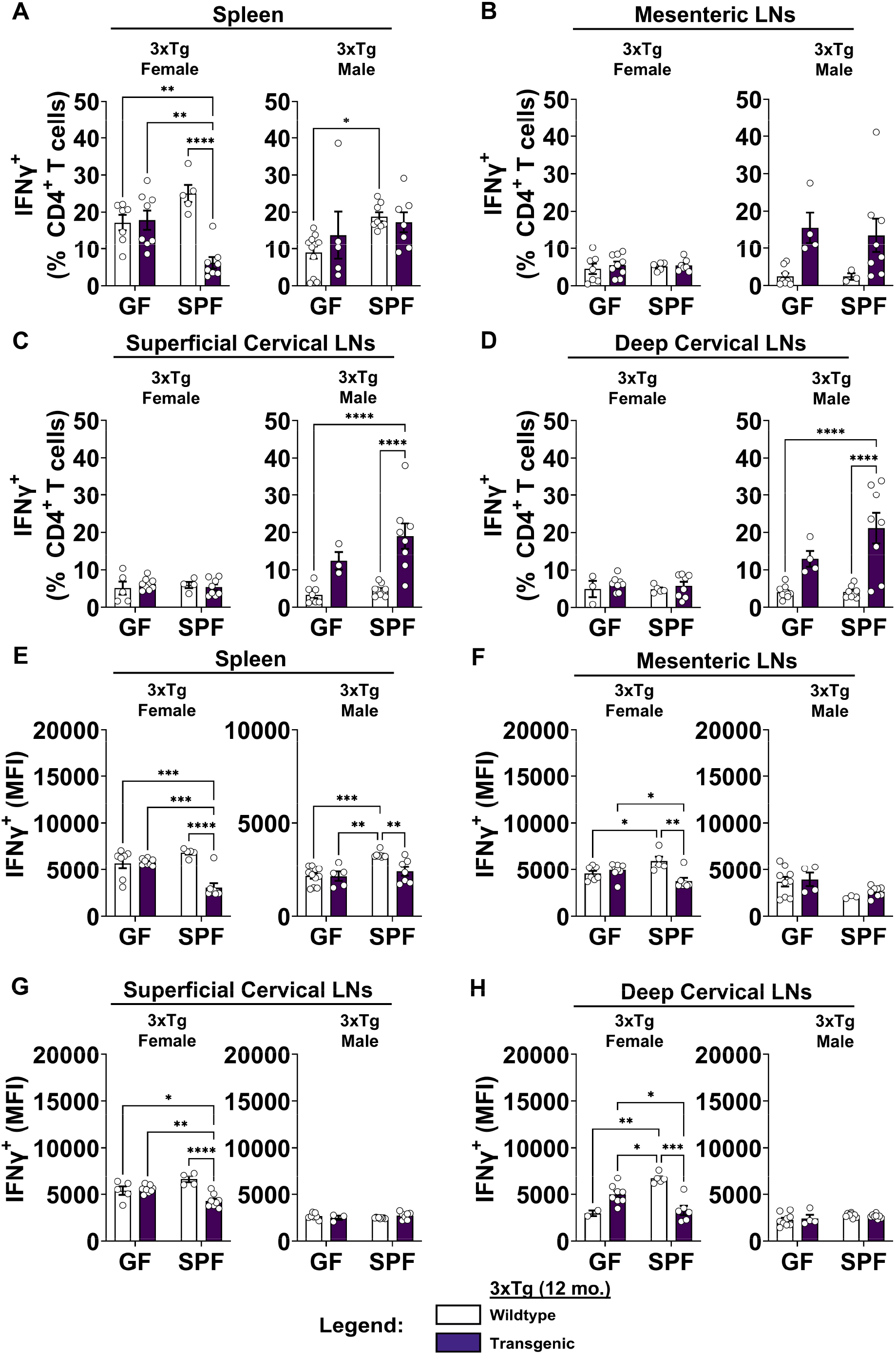
Quantification of IFNγ^+^CD4^+^ T cells in 12-month-old 3xTg mice. (A) Quantification of IFNγ^+^ T cell frequencies in the spleens of female (Left) and male (Right) 3xTg mice compared to wildtype controls in germ-free (GF) and specific pathogen-free (SPF) conditions at 12 months of age. (B) Mesenteric lymph nodes. (C) Superficial cervical lymph nodes. (D) Deep cervical lymph nodes. Data are pooled from 2 independent experiments (n = 3 to 10 per group); two-way ANOVA. (E) Quantification of IFNγ mean fluorescence intensity (MFI) expressed by CD4^+^ T cells in the spleens of female (Left) and male (Right) 3xTg mice compared to wildtype controls in GF and SPF conditions at 12 months of age. (F) Mesenteric lymph nodes. (G) Superficial cervical lymph nodes. (H) Deep cervical lymph nodes. Data are pooled from 2 independent experiments (n = 3 to 10 per group); two-way ANOVA. Error bars indicate SEM. *P < 0.05, **P < 0.01, ***P < 0.001, ****P < 0.0001.

**Supplementary Figure 6.**
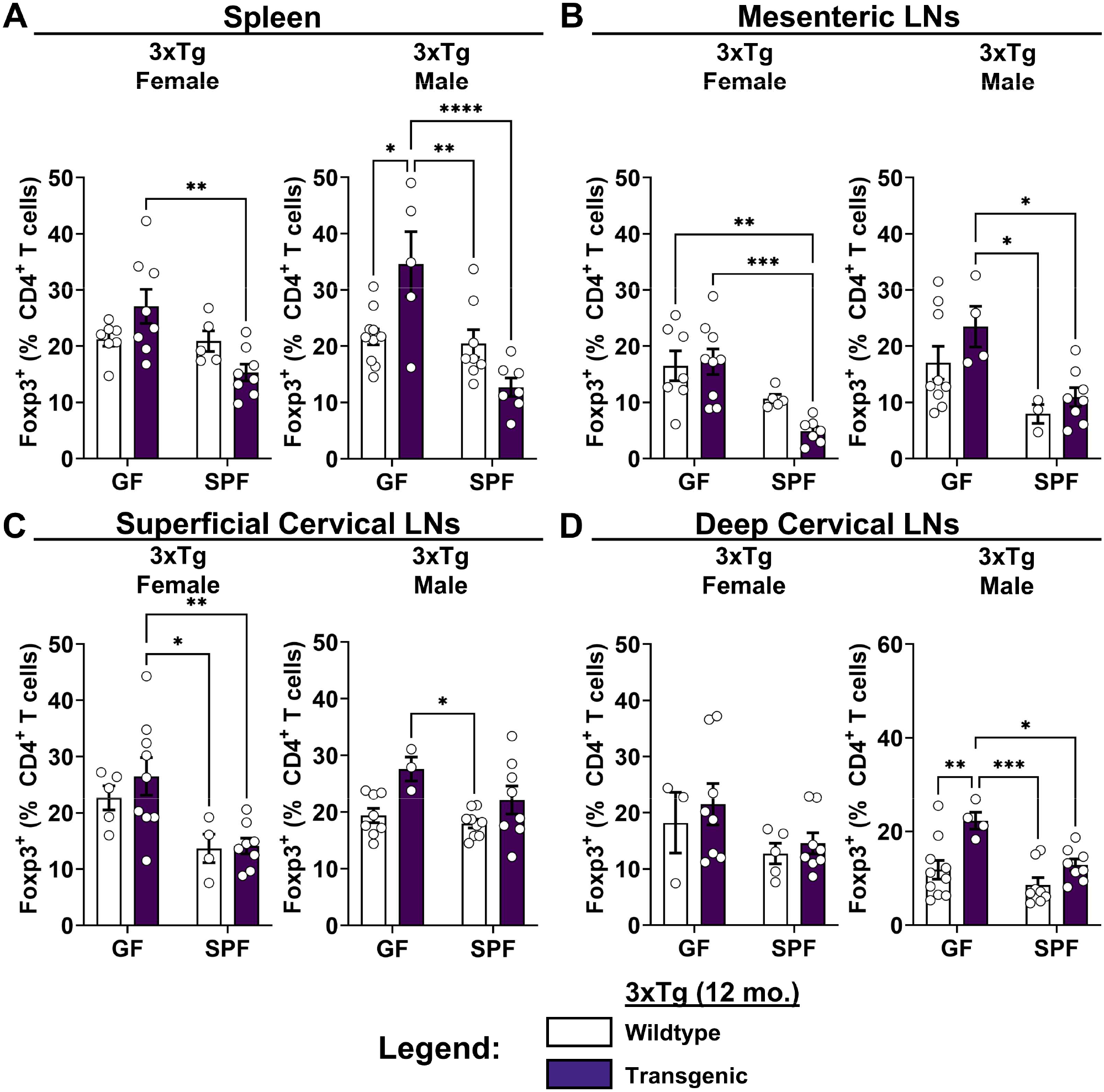
Quantification of Foxp3^+^ T cell frequencies in 12-month-old 3xTg mice. (A) Quantification of Foxp3^+^ T cell frequencies in the spleens of female (Left) and male (Right) 3xTg mice compared to wildtype controls in germ-free (GF) and specific pathogen-free (SPF) conditions at 12 months of age. (B) Mesenteric lymph nodes. (C) Superficial cervical lymph nodes. (D) Deep cervical lymph nodes. Data are pooled from 2 independent experiments (n = 2 to 11 per group); two-way ANOVA. Error bars indicate SEM. *P < 0.05, **P < 0.01, ***P < 0.001, ****P < 0.0001.

**Supplementary Figure 7.**
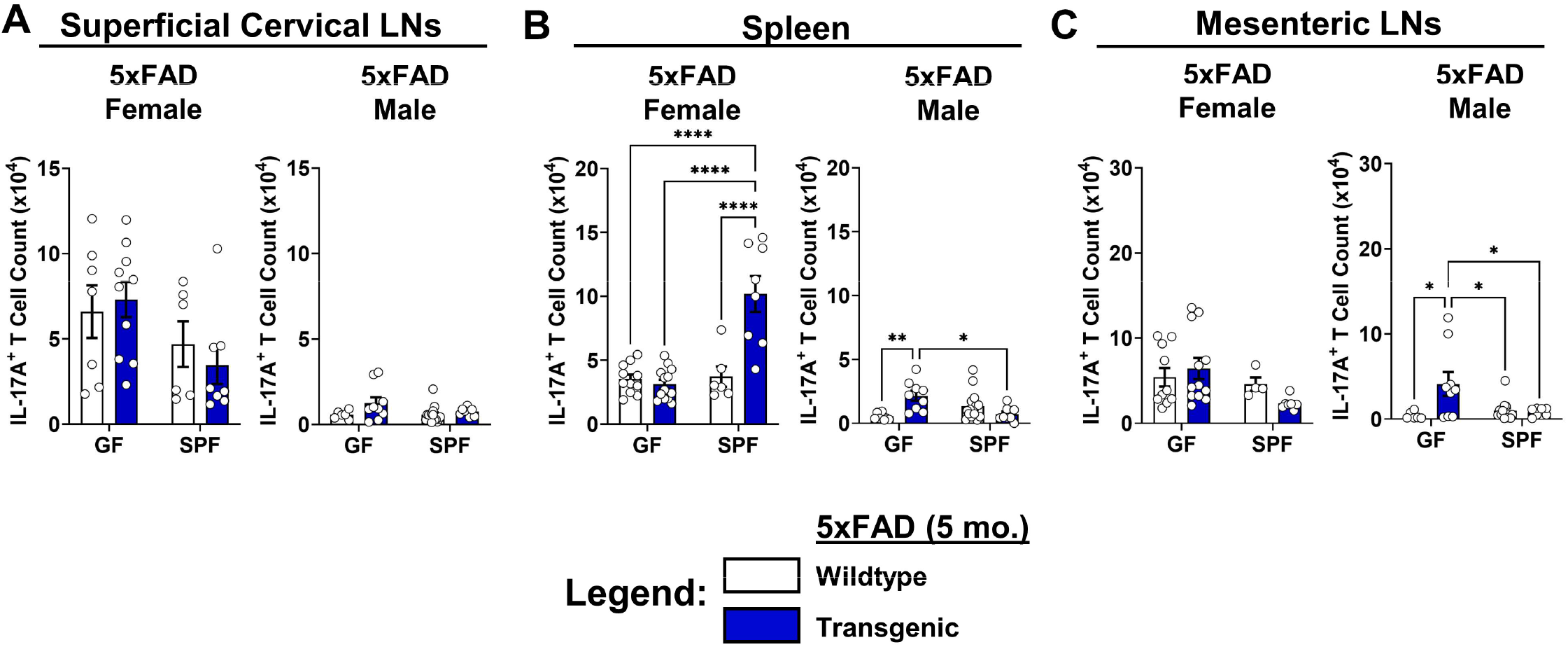
IL-17A^+^ T cell counts in 5xFAD mice. (A) Quantification of IL-17A^+^ T cell counts in the superficial cervical lymph nodes of female (Left) and male (Right) 5xFAD mice compared to wildtype controls in germ-free (GF) and specific pathogen-free (SPF) conditions at 5 months of age. (B) Spleen. (C) Mesenteric lymph nodes. Data are pooled from 2 independent experiments (n = 4 to 15 per group); two-way ANOVA. Error bars indicate SEM. *P < 0.05, **P < 0.01, ***P < 0.001, ****P < 0.0001.

**Supplementary Figure 8.**
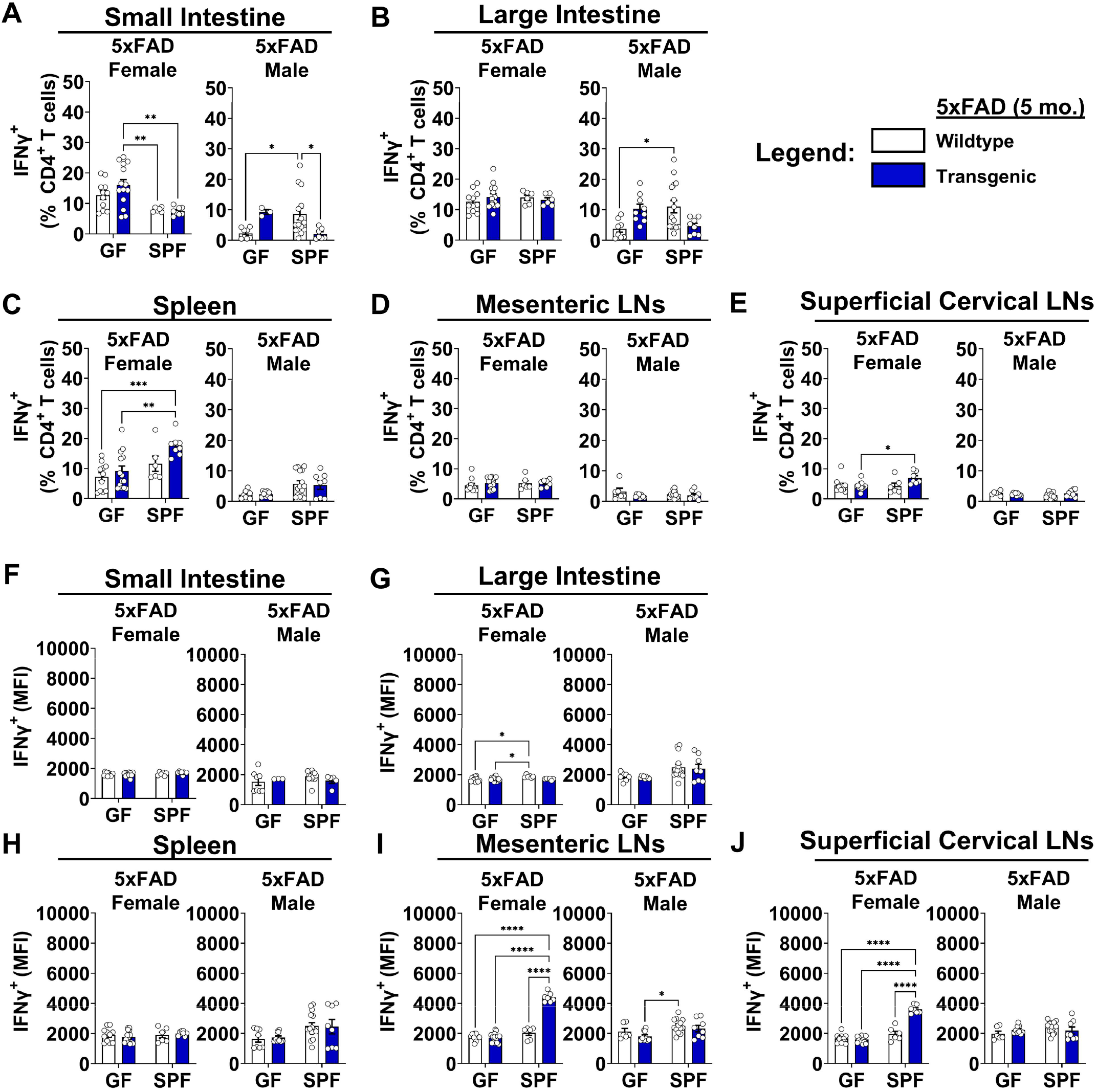
Quantification of IFNγ^+^CD4_+_ T cells in 5-month-old 5xFAD mice. (A) Quantification of IFNγ^+^ T cell frequencies in the small intestines of female (Left) and male (Right) 5xFAD mice compared to wildtype controls in germ-free (GF) and specific pathogen-free (SPF) conditions at 5 months of age. (B) Large intestine. (C) Spleen. (D) Mesenteric lymph nodes. (E) Superficial cervical lymph nodes. (F) Quantification of IFNγ mean fluorescence intensity (MFI) expressed by CD4^+^ T cells in the small intestine. (G) Large intestine. (H) Spleen. (I) Mesenteric lymph nodes. (J) Superficial cervical lymph nodes. Data are pooled from 2 independent experiments (n = 4 to 14 per group); two-way ANOVA. Error bars indicate SEM. *P < 0.05, **P < 0.01, ***P < 0.001, ****P < 0.0001.

**Supplementary Figure 9.**
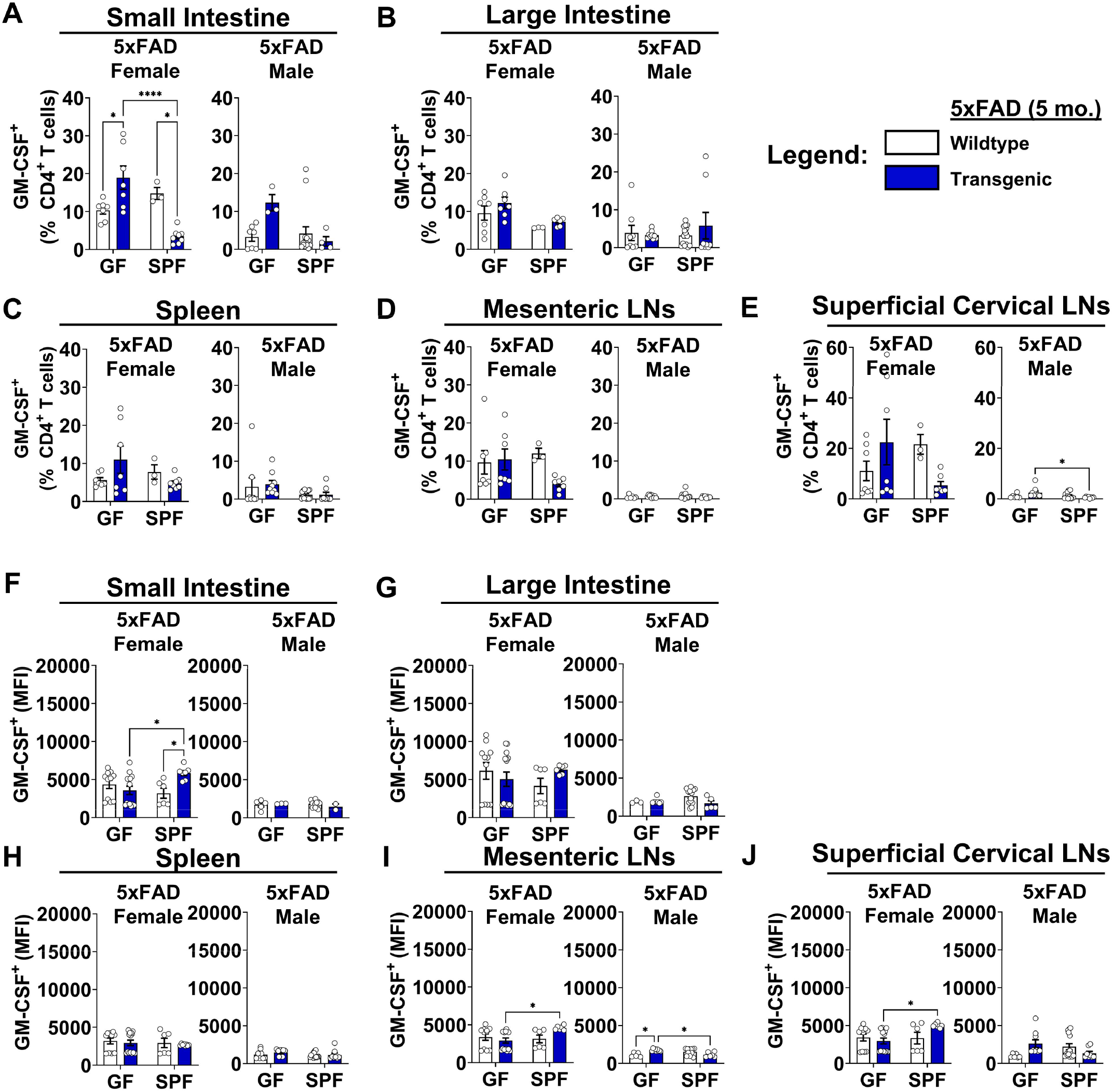
**Quantification of GM-CSF^+^CD4^+^ T cells in 5-month-old 5xFAD mice.** (A) Quantification of GM-CSF^+^ T cell frequencies in the small intestines of female (Left) and male (Right) 5xFAD mice compared to wildtype controls in germ-free (GF) and specific pathogen-free (SPF) conditions at 5 months of age. (B) Large intestine. (C) Spleen. (D) Mesenteric lymph nodes. (E) Superficial cervical lymph nodes. (F) Quantification of GM-CSF mean fluorescence intensity (MFI) expressed by CD4^+^ T cells in the small intestine. (G) Large intestine. (H) Spleen. (I) Mesenteric lymph nodes. (J) Superficial cervical lymph nodes. Data are pooled from 2 independent experiments (n = 4 to 14 per group); two-way ANOVA. Error bars indicate SEM. *P < 0.05, **P < 0.01, ***P < 0.001, ****P < 0.0001.

**Supplementary Figure 10.**
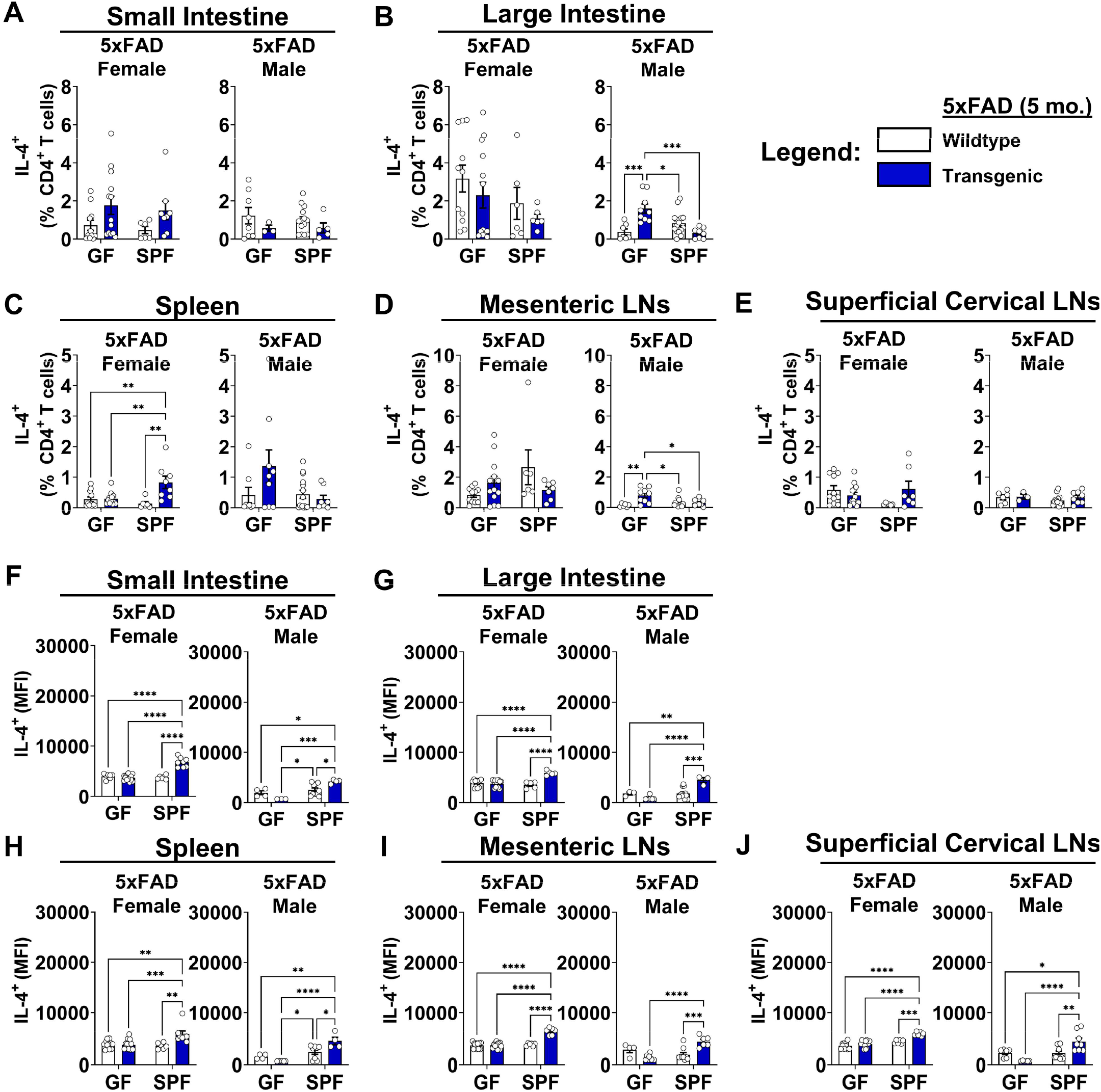
**Quantification of IL-4^+^CD4^+^ T cells in 5-month-old 5xFAD mice.** (A) Quantification of IL-4^+^ T cell frequencies in the small intestines of female (Left) and male (Right) 5xFAD mice compared to wildtype controls in germ-free (GF) and specific pathogen-free (SPF) conditions at 5 months of age. (B) Large intestine. (C) Spleen. (D) Mesenteric lymph nodes. (E) Superficial cervical lymph nodes. (F) Quantification of IL-4 mean fluorescence intensity (MFI) expressed by CD4^+^ T cells in the small intestine. (G) Large intestine. (H) Spleen. (I) Mesenteric lymph nodes. (J) Superficial cervical lymph nodes. Data are pooled from 2 independent experiments (n = 4 to 14 per group); two-way ANOVA. Error bars indicate SEM. *P < 0.05, **P < 0.01, ***P < 0.001, ****P < 0.0001.

**Supplementary Figure 11.**
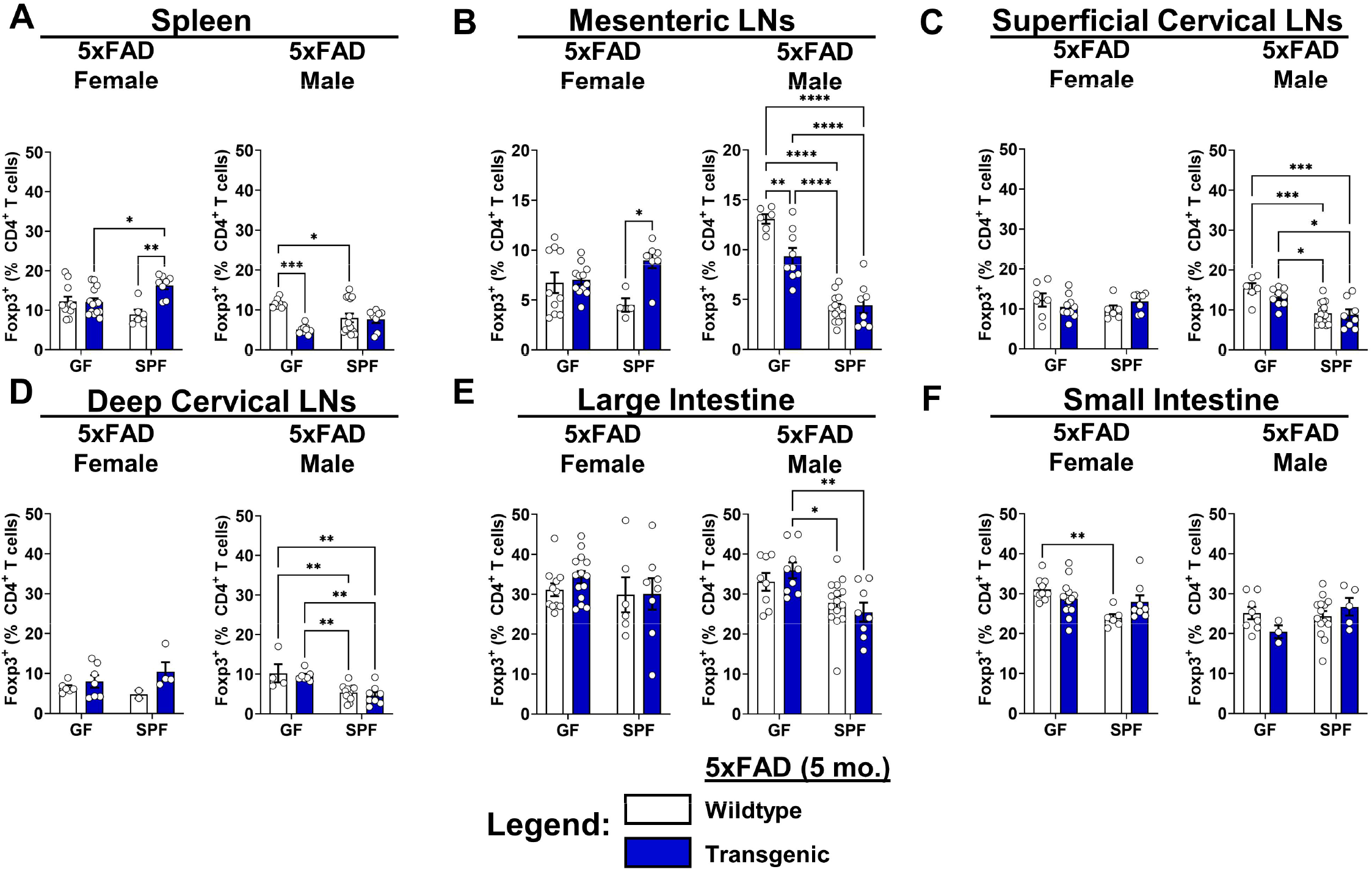
Quantification of Foxp3^+^ T cell frequencies in 5-month-old 5xFAD mice. (A) Quantification of Foxp3^+^ T cell frequencies in the spleens of female (Left) and male (Right) 5xFAD mice compared to wildtype controls in germ-free (GF) and specific pathogen-free (SPF) conditions at 5 months of age. (B) Mesenteric lymph nodes. (C) Superficial cervical lymph nodes. (D) Deep cervical lymph nodes. (E) Large intestine. (F) Small intestine. Data are pooled from 2 independent experiments (n = 2 to 15 per group); two-way ANOVA. Error bars indicate SEM. *P < 0.05, **P < 0.01, ***P < 0.001, ****P < 0.0001.

**Supplementary Figure 12.**
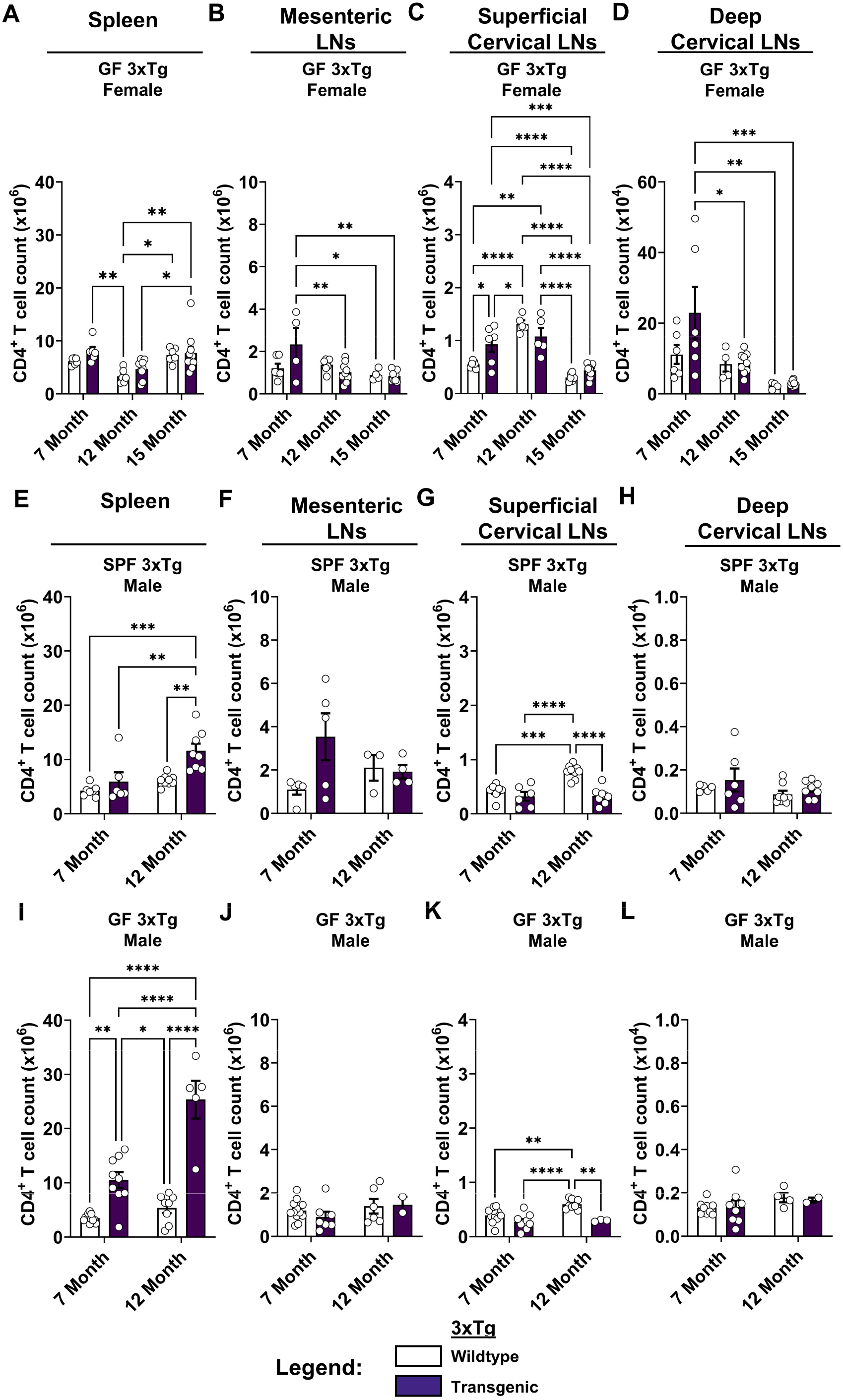
Longitudinal CD4^+^ T cell responses in 3xTg mice. (A) Quantification of CD4^+^ T cell counts in the spleens of female 3xTg mice compared to wildtype controls in germ free (GF) conditions at 7-, 12-, and 15-month ages. (B) Mesenteric lymph nodes. (C) Superficial cervical lymph nodes. (D) Deep cervical lymph nodes. Data are pooled from 5 independent experiments (n = 4 to 10 per group); two-way ANOVA. (E) Quantification of CD4^+^ T cell counts in the spleens of specific pathogen-free (SPF) male 3xTg mice at 7-, 12-, and 15-month ages. (F) Mesenteric lymph nodes. (G) Superficial cervical lymph nodes. (H) Deep cervical lymph nodes. Data are pooled from 5 independent experiments (n = 3 to 9 per group); two-way ANOVA. (I) Quantification of CD4^+^ T cell counts in the spleens of GF male 3xTg mice at 7-, 12-, and 15-month ages. (J) Mesenteric lymph nodes. (K) Superficial cervical lymph nodes. (L) Deep cervical lymph nodes. Data are pooled from 5 independent experiments (n = 2 to 9 per group); two-way ANOVA. Error bars indicate SEM. *P < 0.05, **P < 0.01, ***P < 0.001, ****P < 0.0001.

**Supplementary Figure 13.**
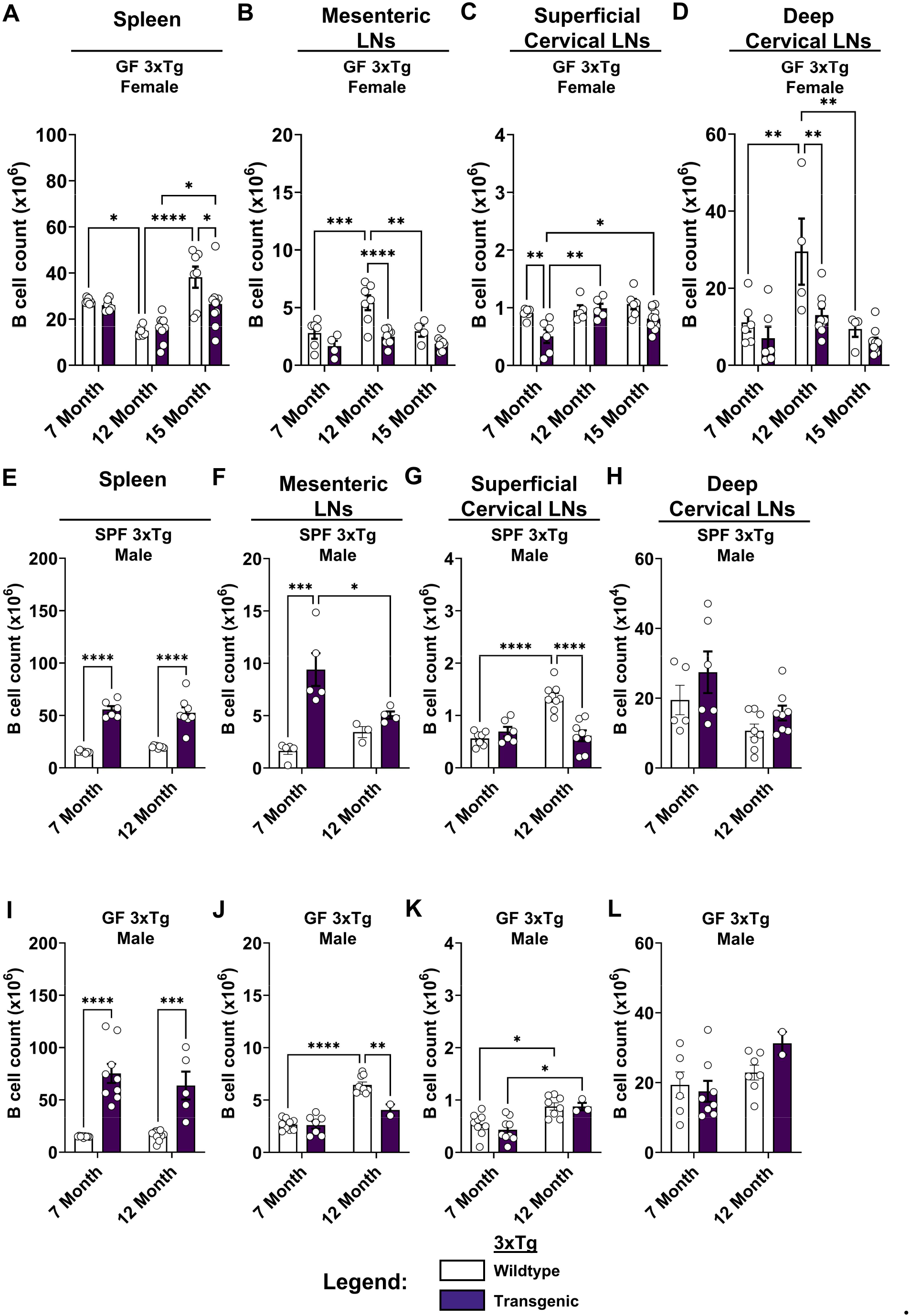
Longitudinal B cell responses in 3xTg mice. (A) Quantification of B cell counts in the spleens of female 3xTg mice compared to wildtype controls in germ free (GF) conditions at 7-, 12-, and 15-month ages. (B) Mesenteric lymph nodes. (C) Superficial cervical lymph nodes. (D) Deep cervical lymph nodes. Data are pooled from 5 independent experiments (n = 4 to 10 per group); two-way ANOVA. (E) Quantification of B cell counts in the spleens of specific pathogen-free (SPF) male 3xTg mice at 7-, 12-, and 15-month ages. (F) Mesenteric lymph nodes. (G) Superficial cervical lymph nodes. (H) Deep cervical lymph nodes. Data are pooled from 5 independent experiments (n = 3 to 9 per group); two-way ANOVA. (I) Quantification of B cell counts in the spleens of GF male 3xTg mice at 7-, 12-, and 15- month ages. (J) Mesenteric lymph nodes. (K) Superficial cervical lymph nodes. (L) Deep cervical lymph nodes. Data are pooled from 5 independent experiments (n = 2 to 11 per group); two-way ANOVA. For clarity, only significant comparisons of the same mouse group between ages or between transgenic and control mice at the same age are shown. Error bars indicate SEM. *P < 0.05, **P < 0.01, ***P < 0.001, ****P < 0.0001.

**Supplementary Figure 14.**
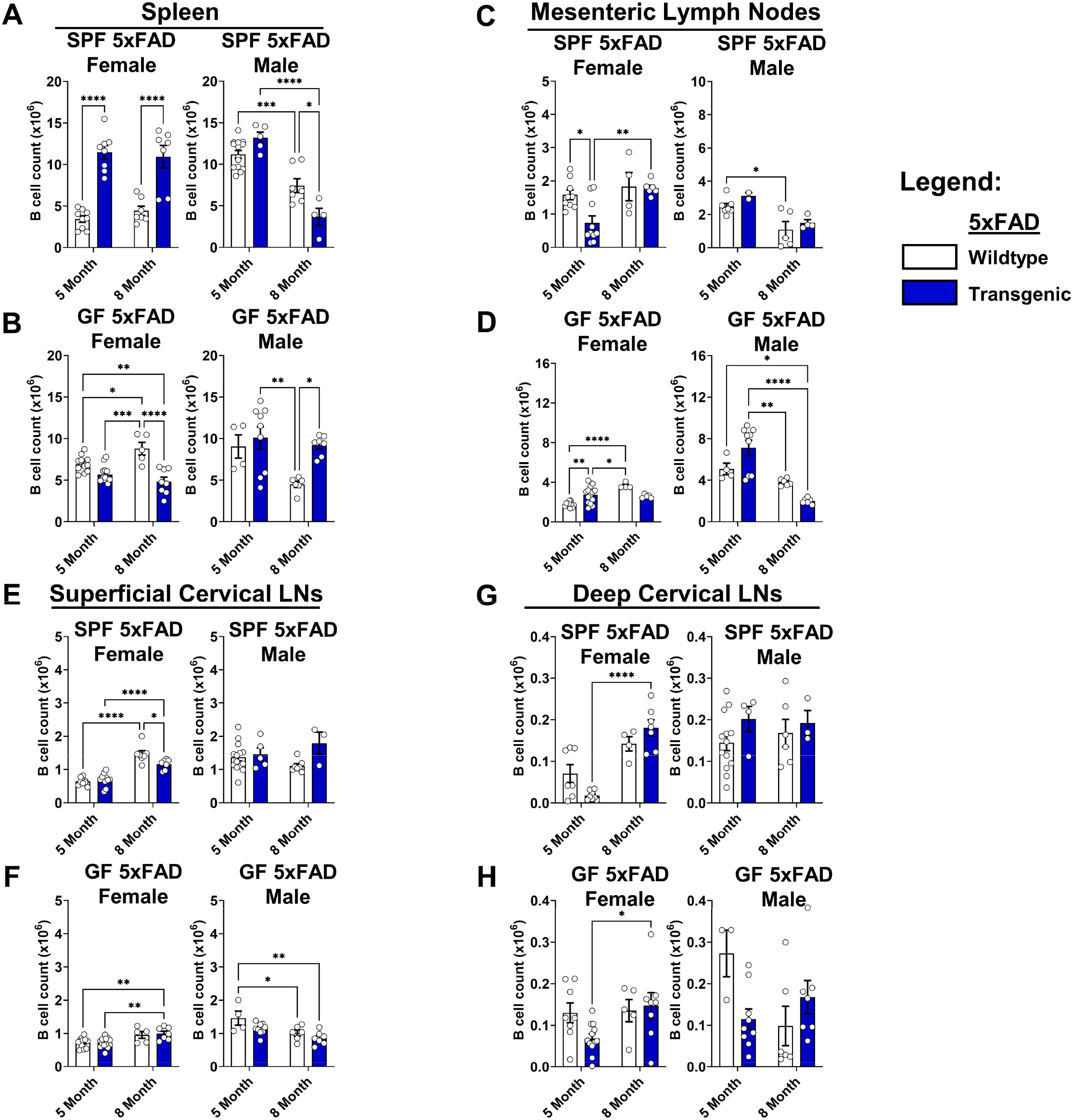
Longitudinal B cell responses in 5xFAD mice. (A) Quantification of B cell counts in the spleens of female (Left) and male (Right) 5xFAD mice compared to wildtype controls in specific pathogen-free (SPF) and (B) germ-free (GF) conditions at 5 and 8 months of age. (C) Mesenteric lymph nodes in SPF and (D) GF conditions. (E) Superficial cervical lymph nodes in SPF and (F) GF conditions. (G) Deep cervical lymph nodes in SPF and (H) GF conditions. Data are pooled from 2 independent experiments (n = 2 to 15 per group); two-way ANOVA. For clarity, only significant comparisons of the same mouse group between ages or between transgenic and control mice at the same age are shown. Error bars indicate SEM. *P < 0.05, **P < 0.01, ***P < 0.001, ****P < 0.0001.

**Supplementary Figure 15.**
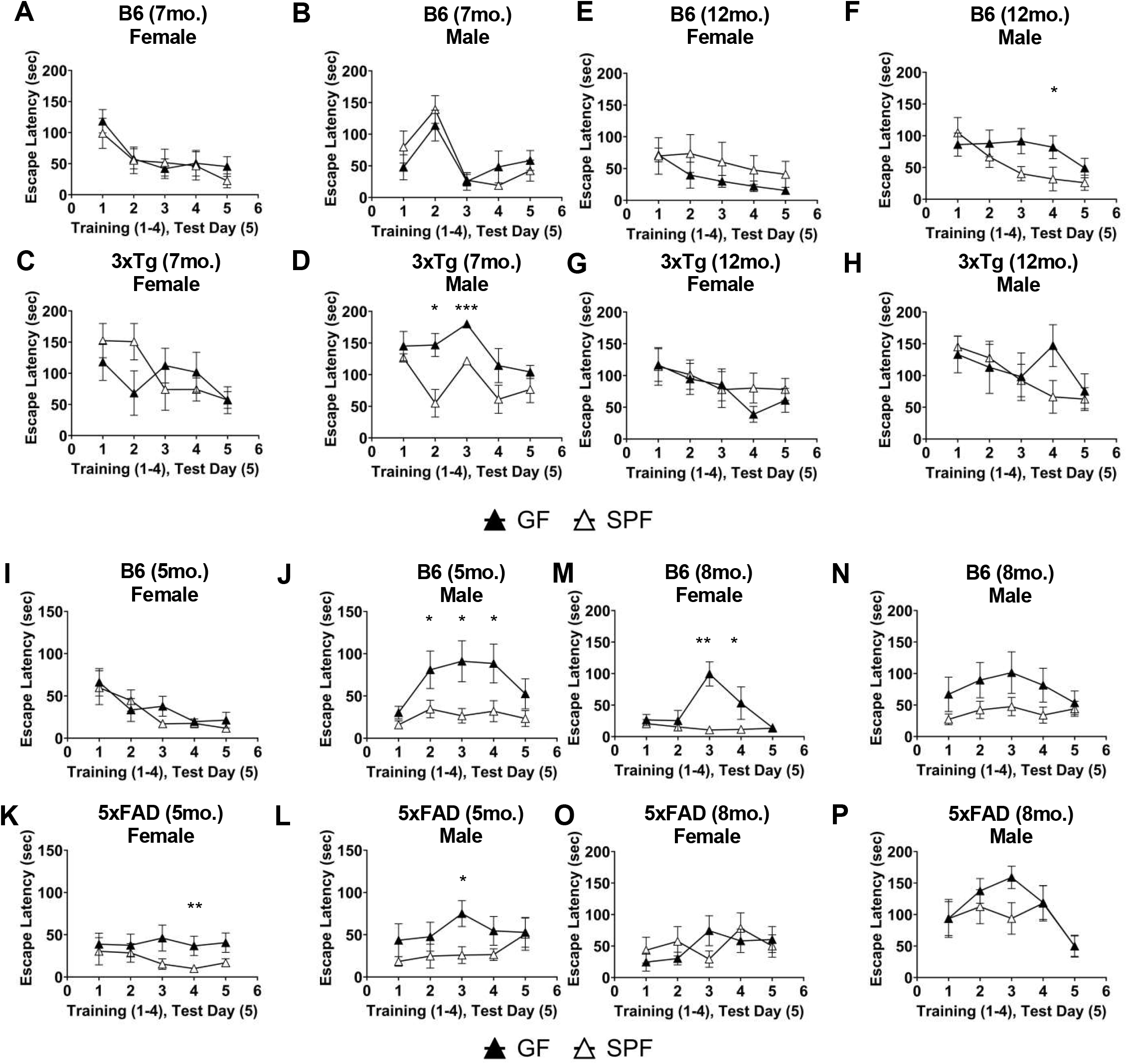
Cognitive deficits in Barnes Maze testing of female and male 7-, 12- and 15- month-old 3xTg and 5- and 8-month-old 5xFAD mice [GF vs. SPF comparison]. (A) Training (1-4) and test (5) day mean escape latency in Barnes Maze for 7-month-old female wildtype (B6) mice in specific pathogen-free (SPF) compared to germ-free (GF) conditions. (B) Male B6 mice. (C) Female 3xTg mice. (D) Male 3xTg mice. Data are pooled from 2 independent experiments (n = 6 to 11 per group); Mann-Whitney U test comparing SPF to GF conditions. (E) 12-month-old female B6 mice in SPF compared to GF conditions. (F) Male B6 mice. (G) Female 3xTg mice. (H) Male 3xTg mice. Data are pooled from 2 independent experiments (n = 5 to 11 per group). (I) 5-month-old female B6 mice in SPF compared to GF conditions. (J) Male B6 mice. (K) Female 5xFAD mice. (L) Male 5xFAD mice. Data are pooled from 2 independent experiments (n = 8 to 14 per group); Mann-Whitney U test comparing SPF to GF conditions. (M) 8-month-old female B6 mice in SPF compared to GF conditions. (N) Male B6 mice. (O) Female 5xFAD mice. (P) Male 5xFAD mice. Data are pooled from 2 independent experiments (n = 5 to 13 per group); Mann-Whitney U test comparing SPF to GF conditions. Error bars indicate SEM. *P < 0.05, **P < 0.01, ***P < 0.001, ****P < 0.0001.

**Supplementary Figure 16.**
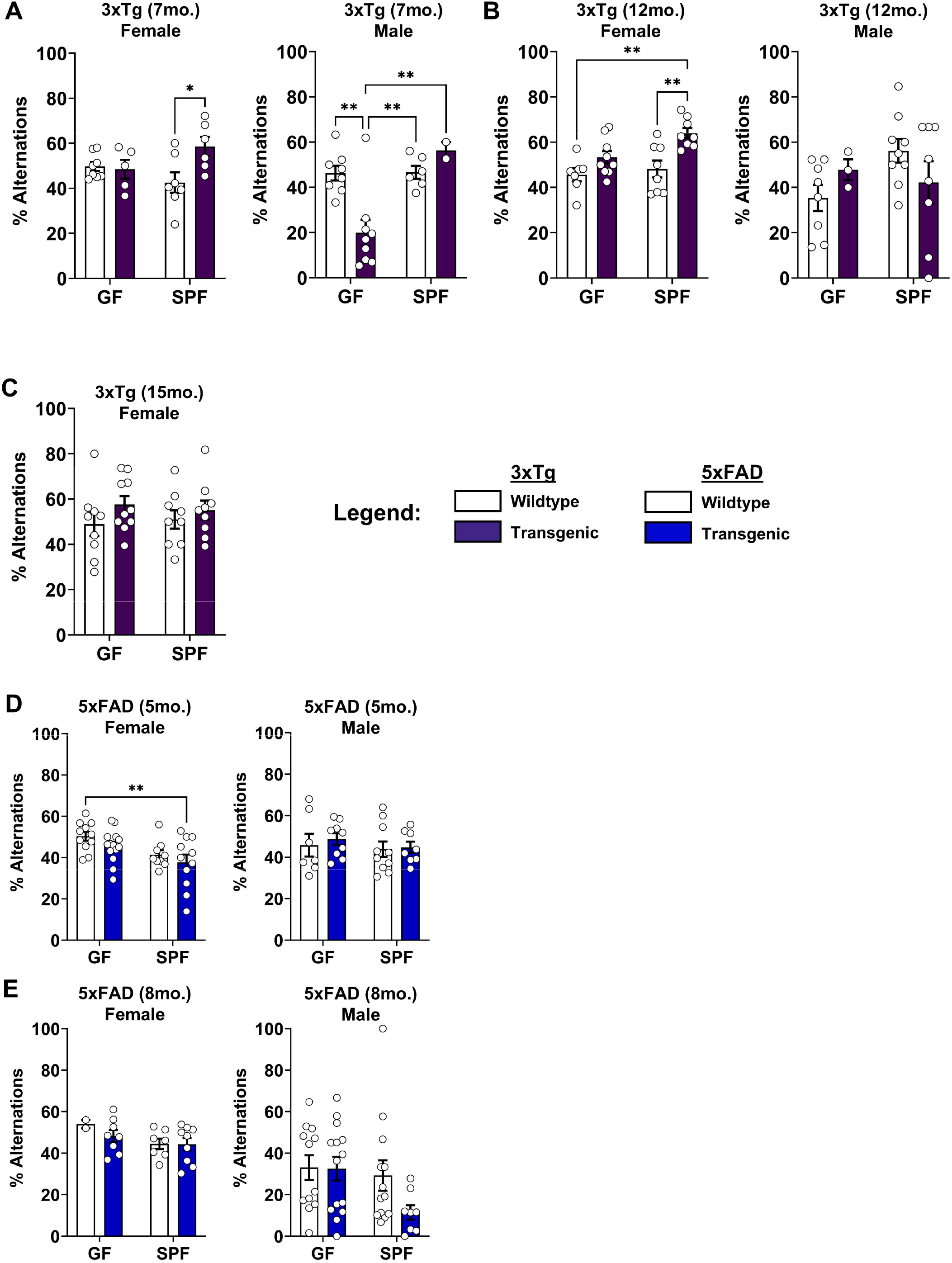
Cognitive deficits in Y-maze testing of female and male 7-, 12- and 15-month-old 3xTg and 5- and 8-month-old 5xFAD mice. (A) Alternations (%) in Y-maze testing of 7-month-old female (Left) and male (Right) 3xTg mice compared to wildtype controls in germ free (GF) or specific pathogen-free (SPF) conditions. (B) 12-month-old female (Left) and male (Right) 3xTg mice. (C) 15-month-old female 3xTg mice. Data are pooled from 2 independent experiments (n = 2 to 14 per group); two-way ANOVA. (D) 5-month-old female (Left) and male (Right) 5xFAD mice. (E) 8-month-old female (Left) and male (Right) 5xFAD mice. Data are pooled from 2 independent experiments (n = 5 to 16 per group); two-way ANOVA. Error bars indicate SEM. *P < 0.05, **P < 0.01, ***P < 0.001, ****P < 0.0001.

**Supplementary Figure 17.**
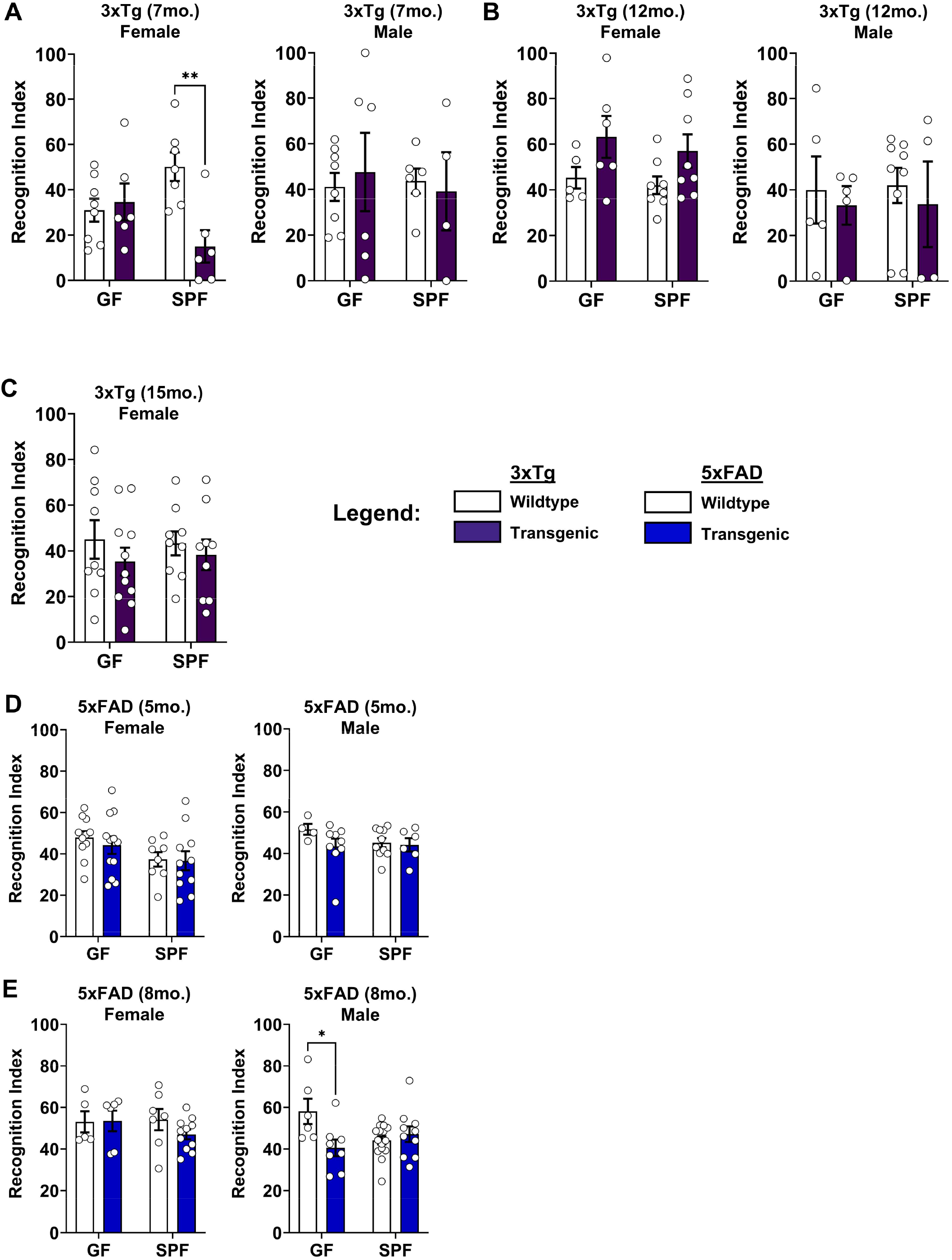
Cognitive deficits in Novel Object Recognition testing of female and male 7-, 12- and 15-month-old 3xTg and 5- and 8-month-old 5xFAD mice. (A) Recognition index in novel object recognition testing for 7-month-old female (Left) and male (Right) 3xTg mice compared to wildtype controls in germ free (GF) or specific pathogen-free (SPF) conditions. (B) 12- month-old female (Left) and male (Right) 3xTg mice. (C) 15-month-old female 3xTg mice. Data are pooled from 2 independent experiments (n = 4 to 14 per group); two-way ANOVA. (D) 5-month-old female (Left) and male (Right) 5xFAD mice. (E) 8-month-old female (Left) and male (Right) 5xFAD mice. Data are pooled from 2 independent experiments (n = 5 to 16 per group); two-way ANOVA. Error bars indicate SEM. *P < 0.05, **P < 0.01, ***P < 0.001, ****P < 0.0001.

