## Supplementary Representative Flow Cytometry Plots for "The microbiome shapes immunity in a sex-specific manner in mouse models of Alzheimer’s disease"

**Abbreviations:**

| | |
|---|---|
| 5M | 5 months |
| 7M | 7 months |
| 8M | 8 months |
| 12M | 12 months |
| 15M | 15 months |
| GF | Germ- free |
| DC | Deep cervical lymph nodes |
| LI | Large Intestine |
| MLN | Mesenteric lymph nodes |
| SC | Superficial cervical lymph nodes |
| SI | Small intestine |
| SP | Spleen |
| SPF | Specific pathogen- free |

### Representative Flow Cytometry Plots for Fig. 1, S2: $CD4^+$ and $CD8^+$ T cells [3xTg]

**3xTg 7M Females: SP**
**$CD45^+CD3^+TCR\beta^+$**

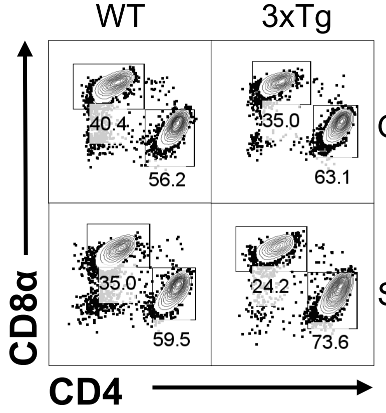


**3xTg 7M Females: MLN**
**$CD45^+CD3^+TCR\beta^+$**

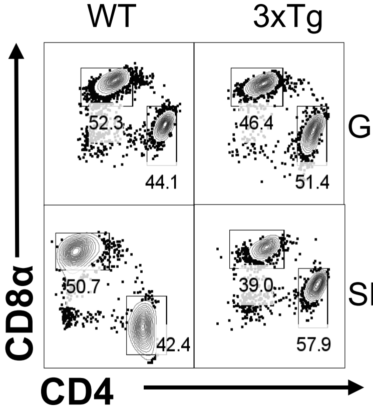


**3xTg 7M Females: SC**
**$CD45^+CD3^+TCR\beta^+$**

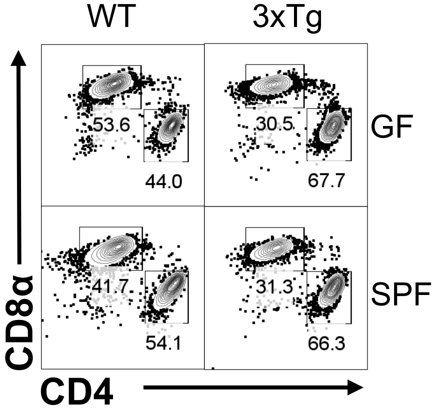


**3xTg 7M Females: DC**
**$CD45^+CD3^+TCR\beta^+$**

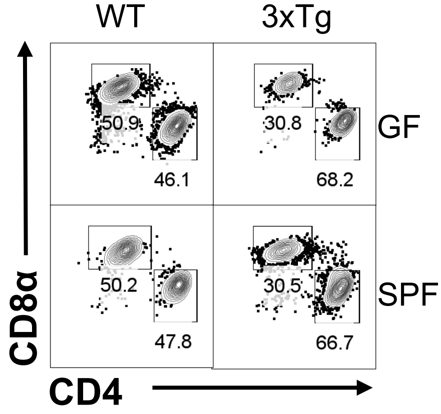


**3xTg 7M Males: SP**
**$CD45^+CD3^+TCR\beta^+$**

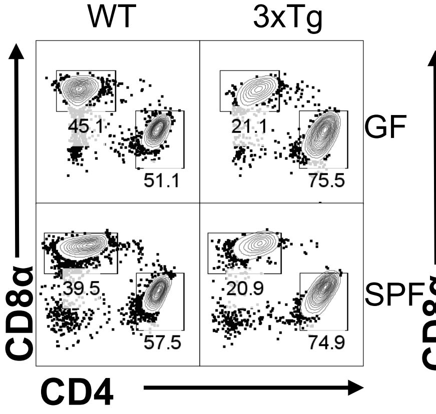


**3xTg 7M Males: MLN**
**$CD45^+CD3^+TCR\beta^+$**

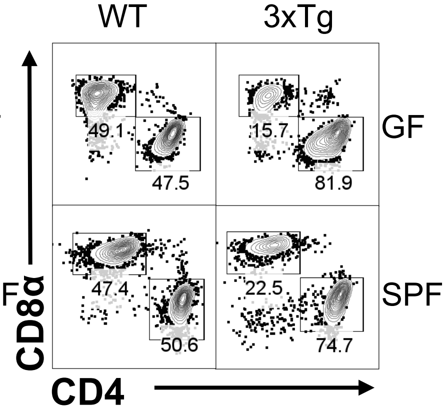


**3xTg 7M Males: SC**
**$CD45^+CD3^+TCR\beta^+$**

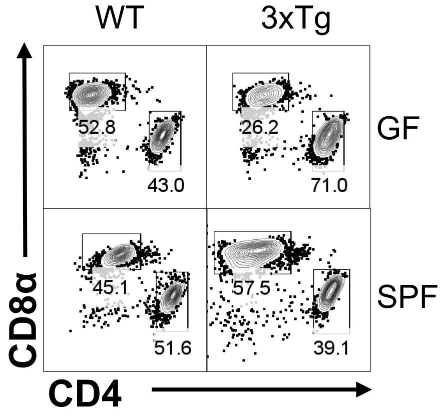


**3xTg 7M Males: DC**
**$CD45^+CD3^+TCR\beta^+$**

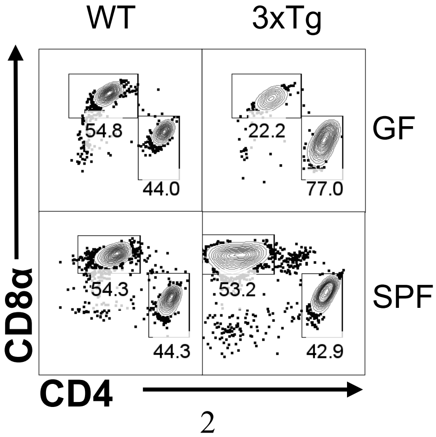

### Representative Flow Cytometry Plots for Fig. 1: CD4$^+$ and CD8$^+$ T cells [5xFAD]

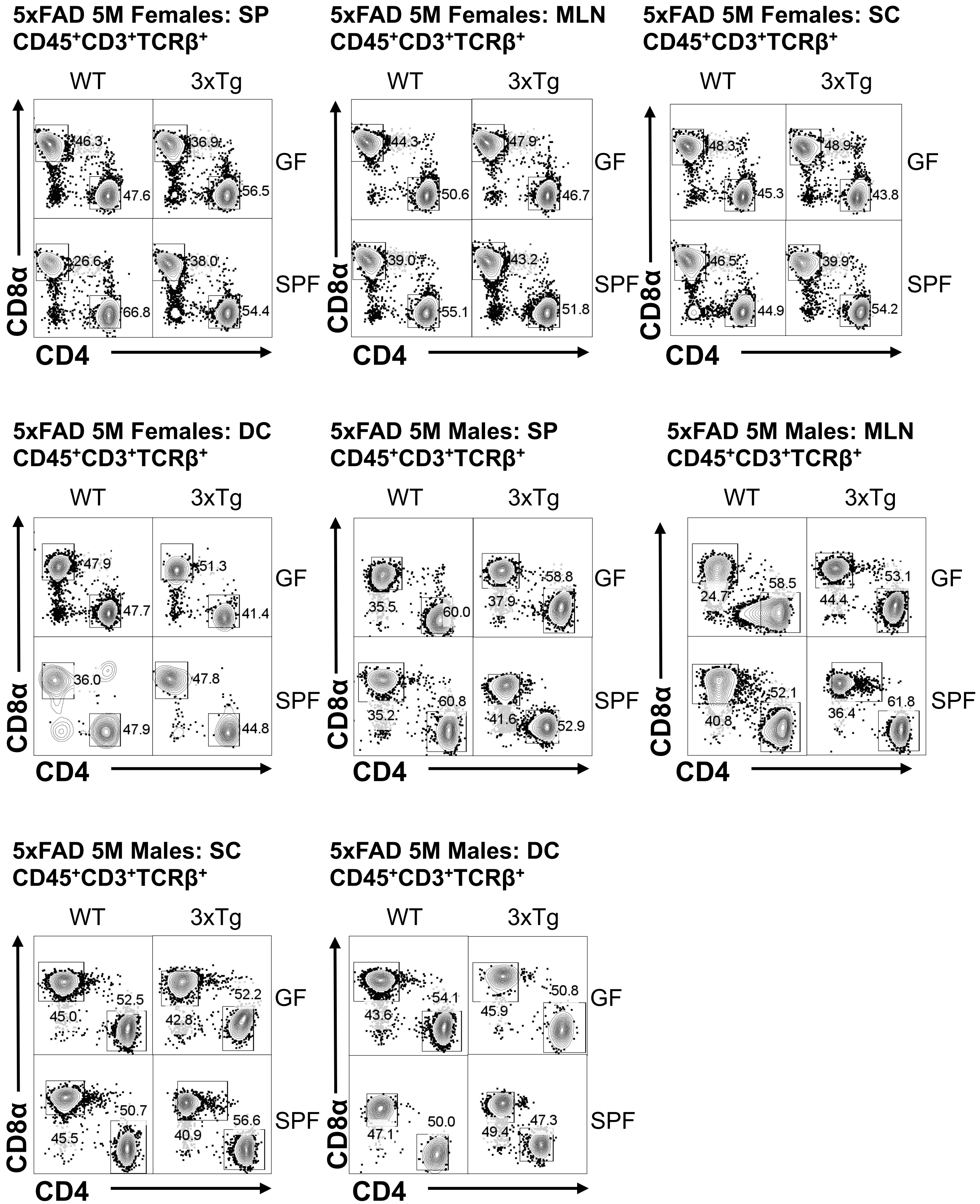

### Representative Flow Cytometry Plots for Fig. 2, S4: IFNγ+/IL-17A+ T cells [3xTg]

**3xTg 12M Females: SP**
**CD45+CD4+TCRβ+Foxp3−**

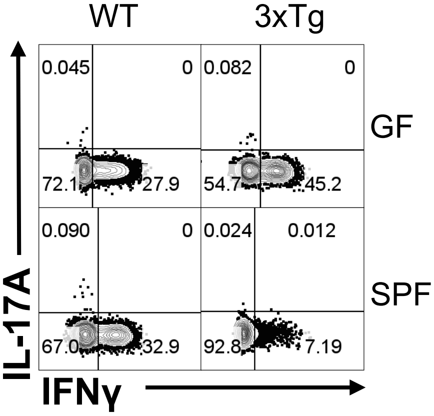


**3xTg 12M Females: MLN**
**CD45+CD4+TCRβ+Foxp3−**

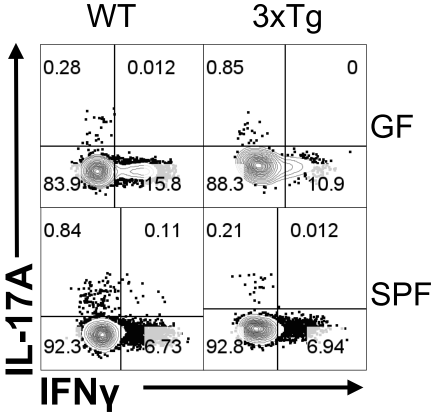


**3xTg 12M Females: SC**
**CD45+CD4+TCRβ+Foxp3−**

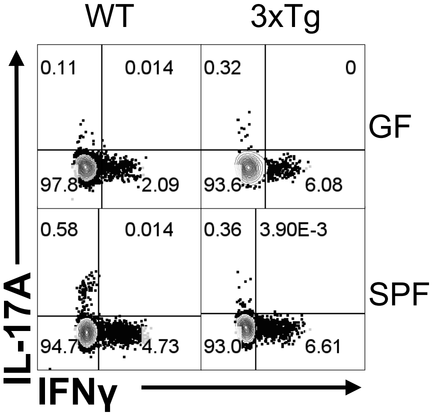


**3xTg 12M Females: DC**
**CD45+CD4+TCRβ+Foxp3−**

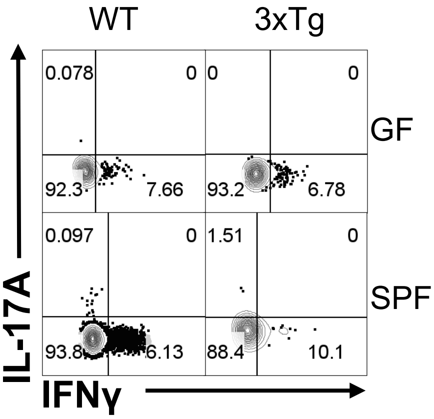


**3xTg 12M Males: SP**
**CD45+CD4+TCRβ+Foxp3−**

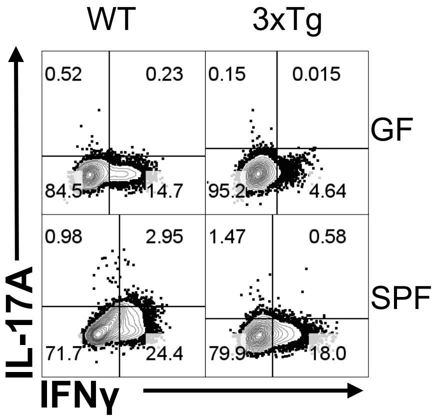


**3xTg 12M Males: MLN**
**CD45+CD4+TCRβ+Foxp3−**

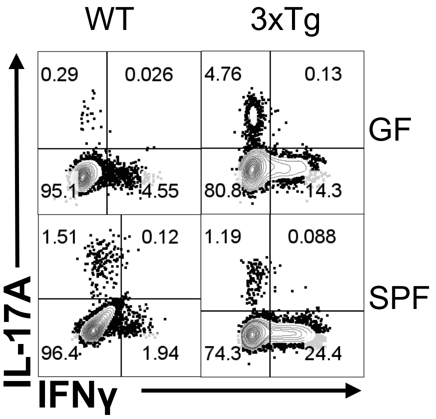


**3xTg 7M Males: SC**
**CD45+CD4+TCRβ+Foxp3−**

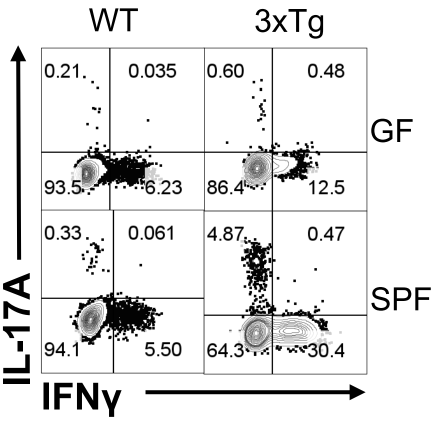


**3xTg 7M Males: DC**
**CD45+CD4+TCRβ+Foxp3−**

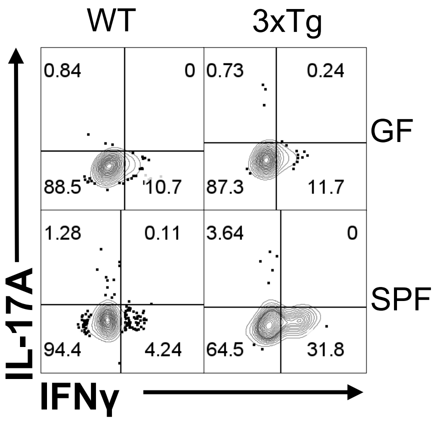

### Representative Flow Cytometry Plots for Fig. 2: IL-17A MFI [3xTg]

**3xTg 12M Females: SP**
**CD45⁺CD4⁺TCRβ⁺Foxp3⁻**

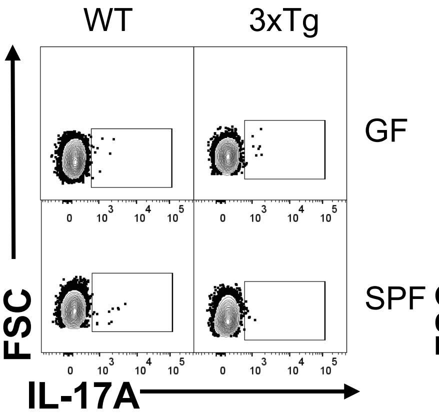


**3xTg 12M Females: MLN**
**CD45⁺CD4⁺TCRβ⁺Foxp3⁻**

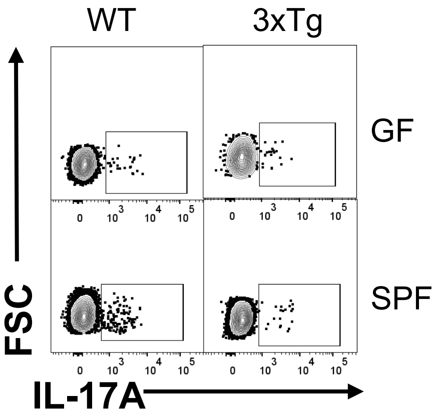


**3xTg 12M Females: SC**
**CD45⁺CD4⁺TCRβ⁺Foxp3⁻**

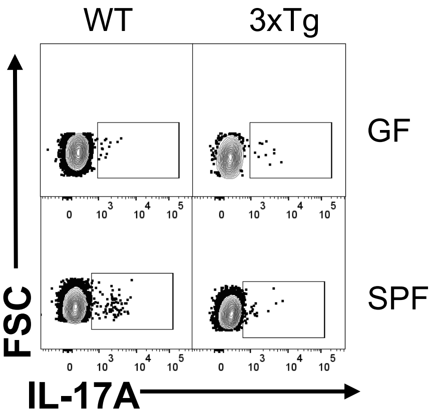


**3xTg 12M Females: DC**
**CD45⁺CD4⁺TCRβ⁺Foxp3⁻**

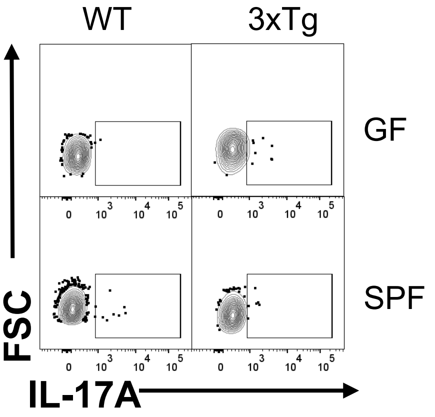


**3xTg 12M Males: SP**
**CD45⁺CD4⁺TCRβ⁺Foxp3⁻**

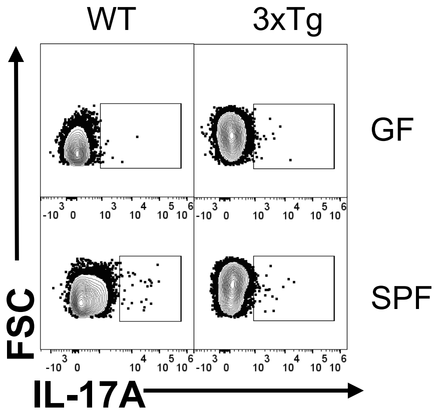


**3xTg 12M Males: MLN**
**CD45⁺CD4⁺TCRβ⁺Foxp3⁻**

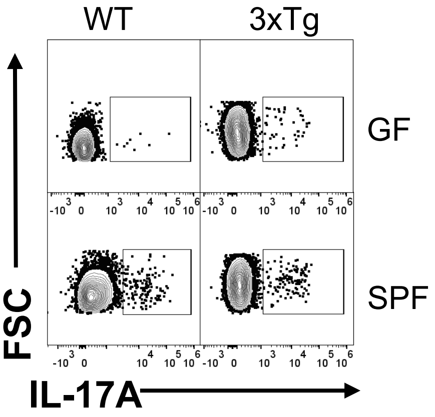


**3xTg 7M Males: SC**
**CD45⁺CD4⁺TCRβ⁺Foxp3⁻**

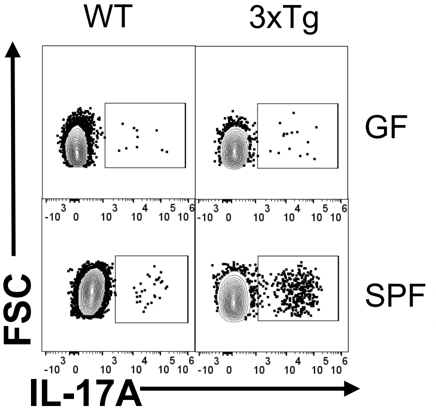


**3xTg 7M Males: DC**
**CD45⁺CD4⁺TCRβ⁺Foxp3⁻**

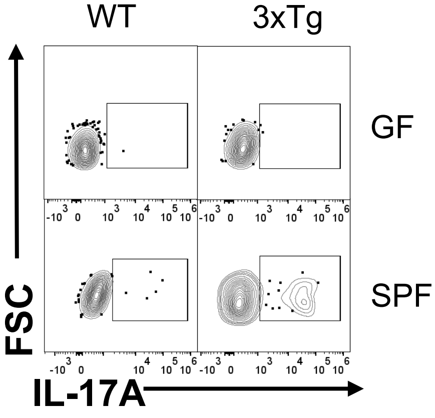

### Representative Flow Cytometry Plots for Fig. 3, S7: IL-17A$^+$ T cells and MFI [5xFAD Females]

**5xFAD 5M Females: SI**
**CD45$^+$CD4$^+$TCRβ$^+$Foxp3$^-$**

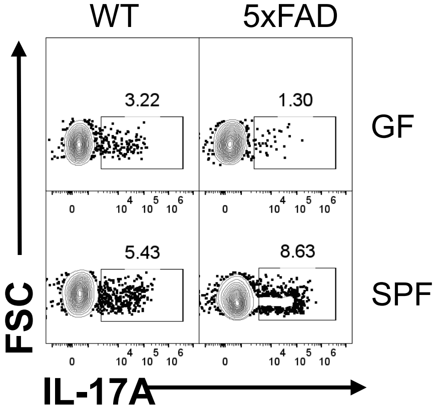


**5xFAD 5M Females: LI**
**CD45$^+$CD4$^+$TCRβ$^+$Foxp3$^-$**

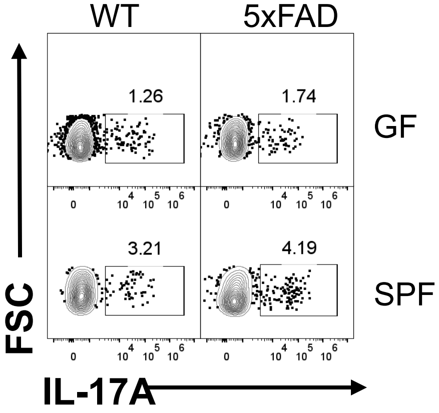


**5xFAD 5M Females: SP**
**CD45$^+$CD4$^+$TCRβ$^+$Foxp3$^-$**

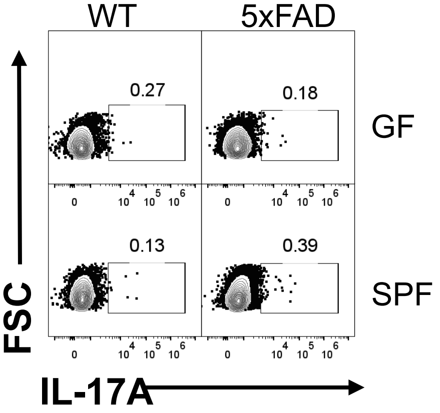


**5xFAD 5M Females: MLN**
**CD45$^+$CD4$^+$TCRβ$^+$Foxp3$^-$**

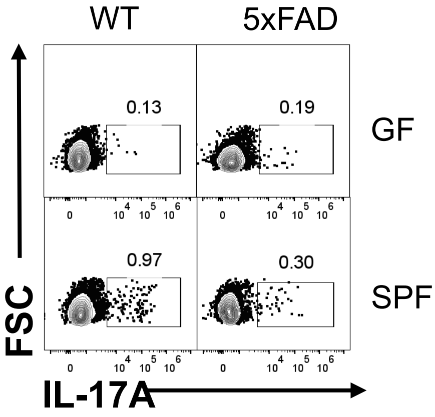


**5xFAD 5M Females: SC**
**CD45$^+$CD4$^+$TCRβ$^+$Foxp3$^-$**

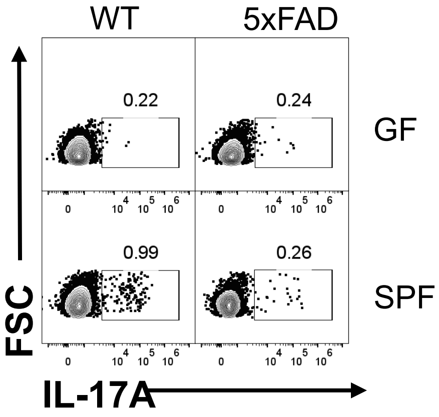

### Representative Flow Cytometry Plots for Fig. 3, S7: IL-17A⁺ T cells and MFI [5xFAD Males]

**5xFAD 5M Males: SI**
**CD45⁺CD4⁺TCRβ⁺Foxp3⁻**

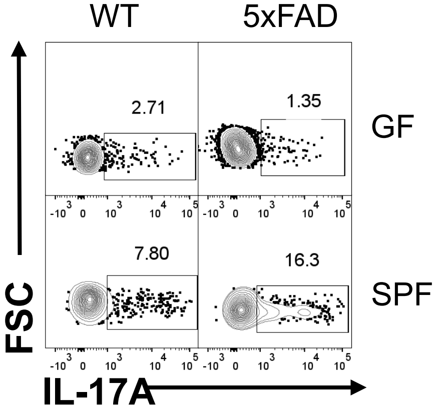


**5xFAD 5M Males: LI**
**CD45⁺CD4⁺TCRβ⁺Foxp3⁻**

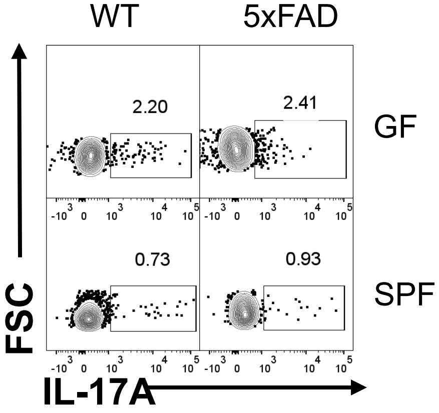


**5xFAD 5M Males: SP**
**CD45⁺CD4⁺TCRβ⁺Foxp3⁻**

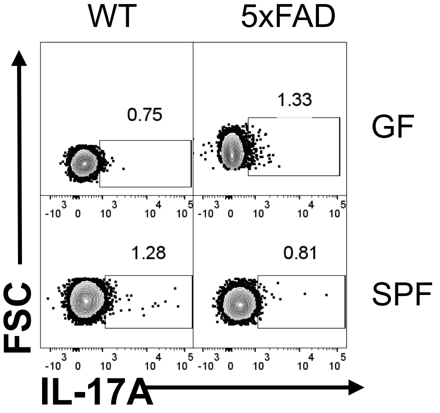


**5xFAD 5M Males: MLN**
**CD45⁺CD4⁺TCRβ⁺Foxp3⁻**

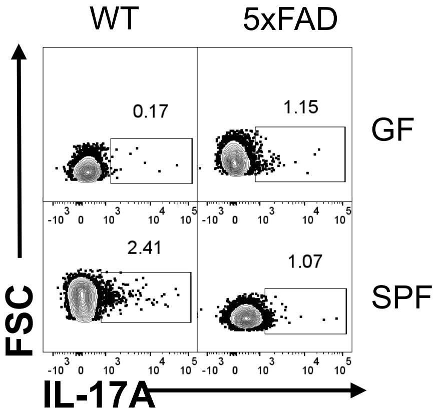


**5xFAD 5M Males: SC**
**CD45⁺CD4⁺TCRβ⁺Foxp3⁻**

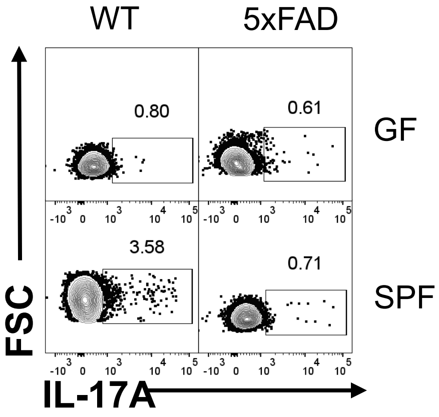

### Representative Flow Cytometry Plots for Fig. 4, S5, S12: CD4$^+$ and CD8$^+$ T cells [3xTg SPF Females]

**3xTg SPF Females: SP**
**CD45$^+$CD3$^+$TCRβ$^+$**

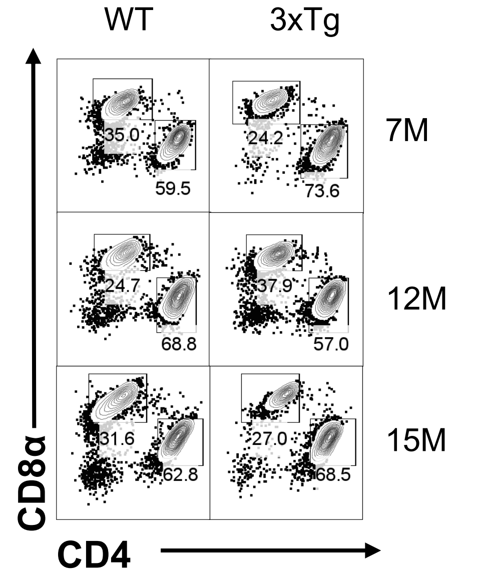


**3xTg SPF Females: MLN**
**CD45$^+$CD3$^+$TCRβ$^+$**

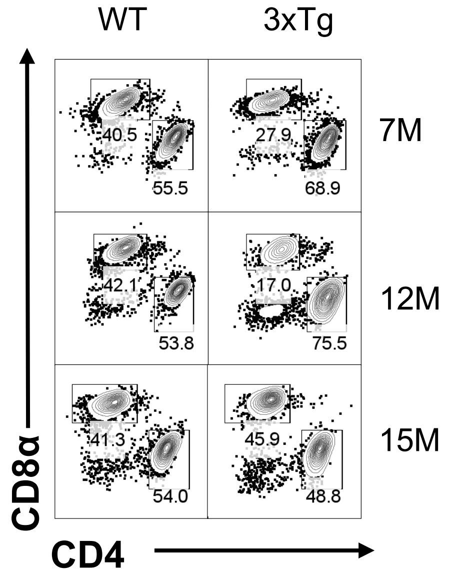


**3xTg SPF Females: SC**
**CD45$^+$CD3$^+$TCRβ$^+$**

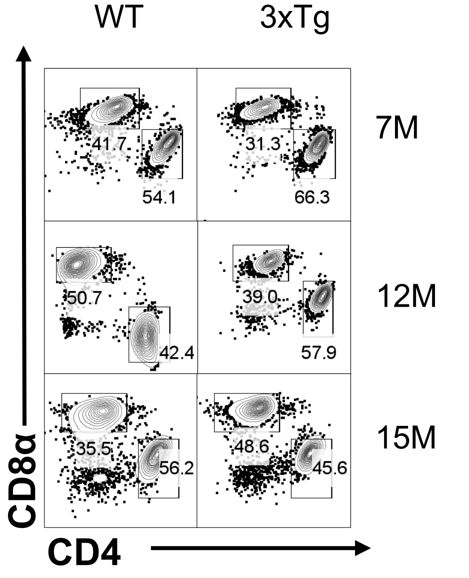


**3xTg SPF Females: DC**
**CD45$^+$CD3$^+$TCRβ$^+$**

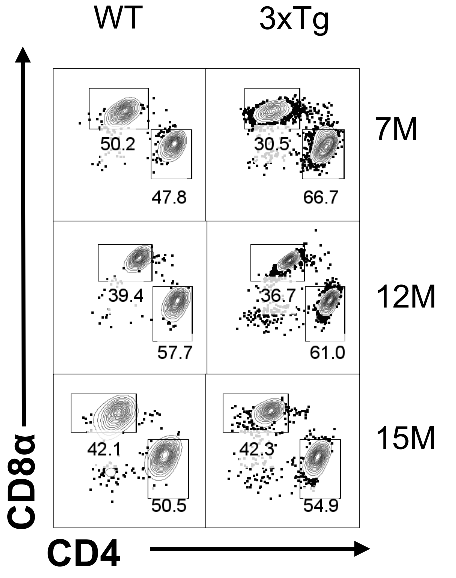

### Representative Flow Cytometry Plots for Fig. 4, S13, S14: T and B cells [3xTg SPF Females]

**3xTg SPF Females: SP**
**CD45⁺**

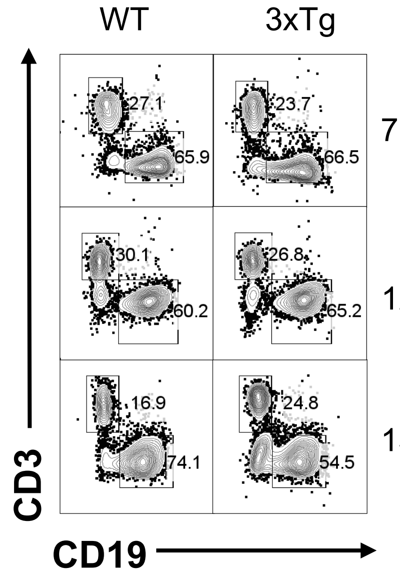


**3xTg SPF Females: MLN**
**CD45⁺**

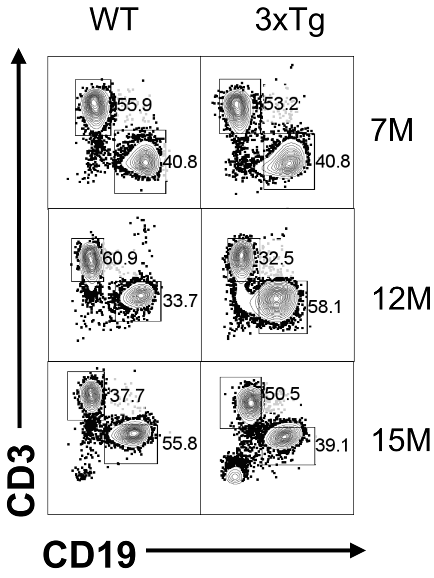


**3xTg SPF Females: SC**
**CD45⁺**

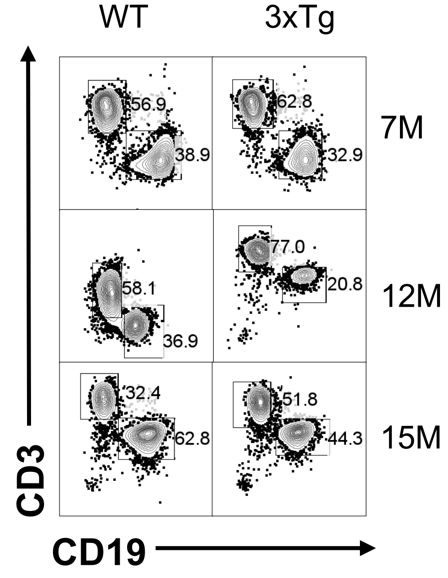


**3xTg SPF Females: DC**
**CD45⁺**

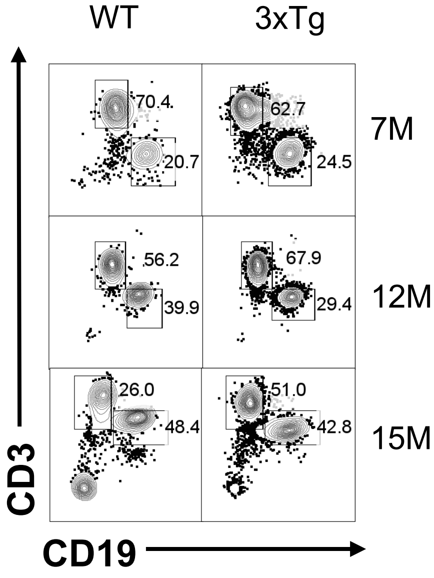

### Representative Flow Cytometry Plots for Fig. 4: IL-17A+ T cells and MFI [5xFAD Females]

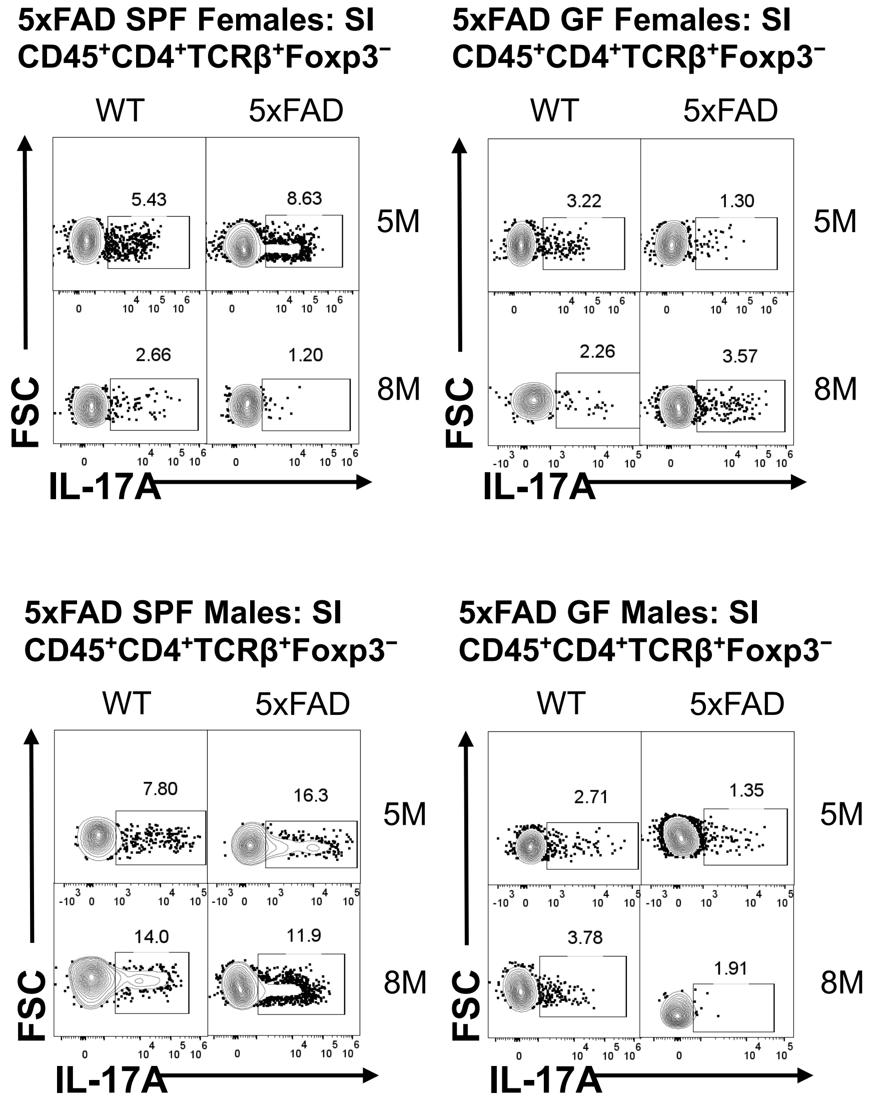

### Representative Flow Cytometry Plots for Fig. S2, S3, S14: T and B cells [5xFAD 5M]

**5xFAD 5M Females: SP CD45+**

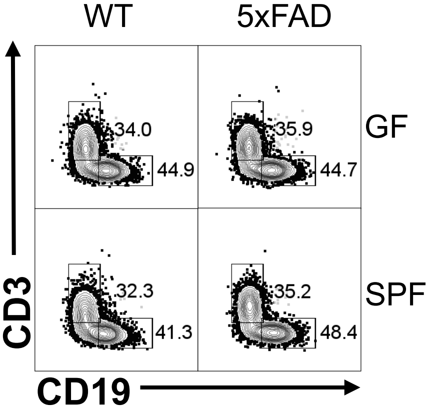


**5xFAD 5M Females: MLN CD45+**

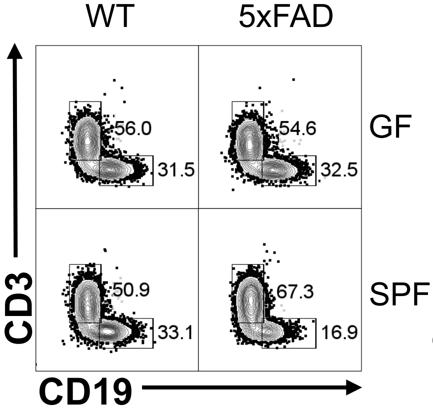


**5xFAD 5M Females: SC CD45+**

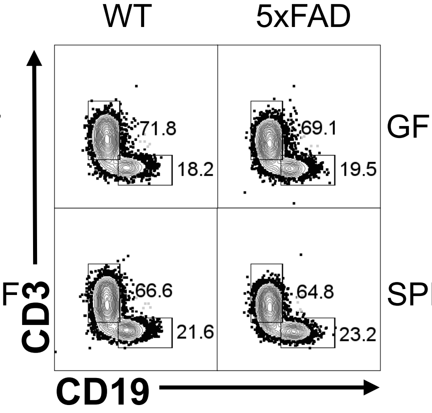


**5xFAD 5M Females: DC CD45+**

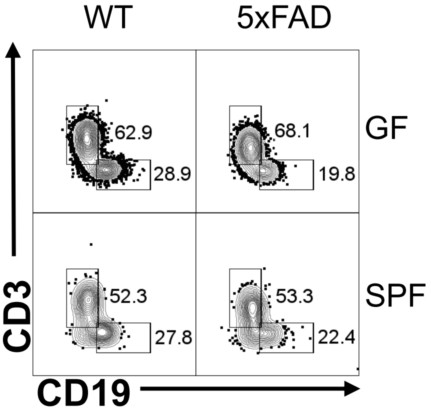


**5xFAD 5M Males: SP CD45+**

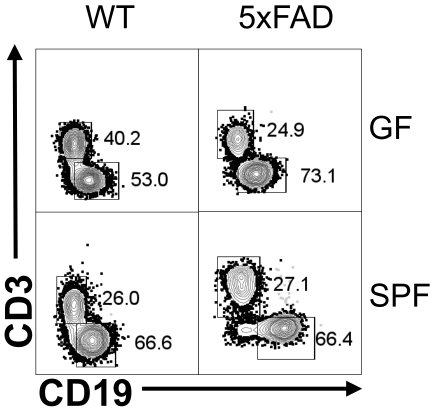


**5xFAD 5M Males: MLN CD45+**

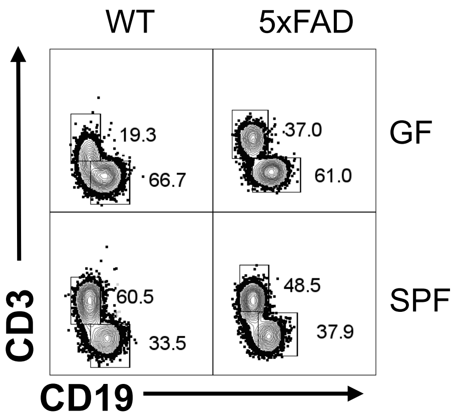


**5xFAD 5M Males: SC CD45+**

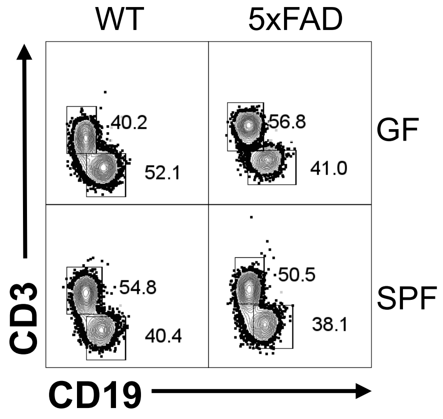


**5xFAD 5M Males: DC CD45+**

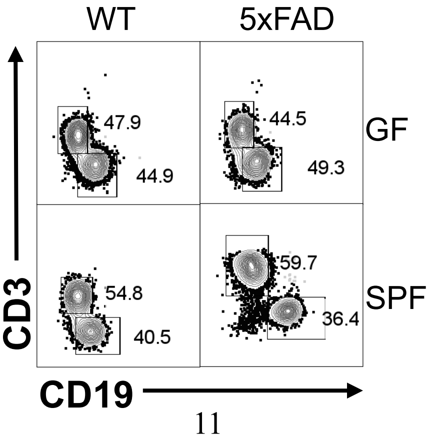

### Representative Flow Cytometry Plots for Fig. S3: T and B cells [5xFAD 5M]

**5xFAD 5M Females: SI**
**CD45+**

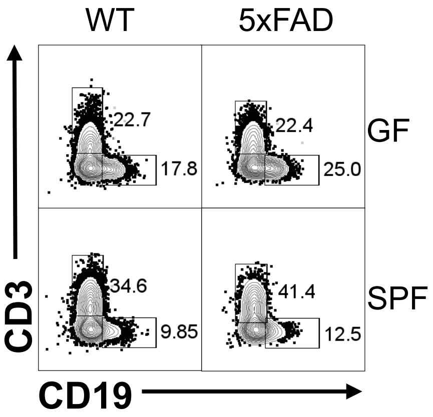


**5xFAD 5M Females: LI**
**CD45+**

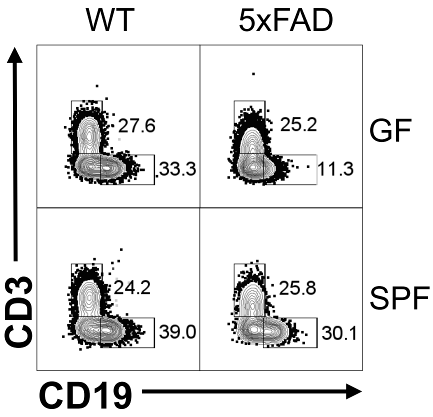


**5xFAD 5M Males: SI**
**CD45+**

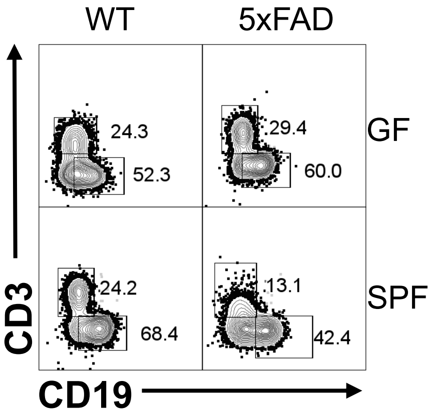


**5xFAD 5M Males: LI**
**CD45+**

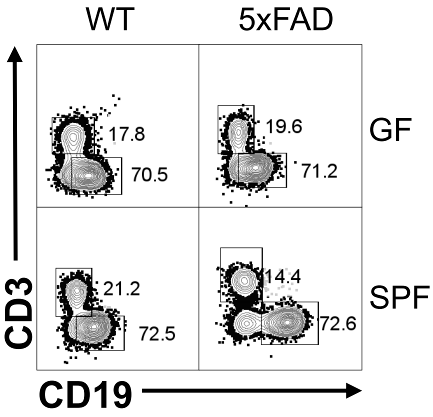

### Representative Flow Cytometry Plots for Fig. S4, S12: CD4$^+$ and CD8$^+$ T cells [3xTg SPF Males]

**3xTg SPF Males: SP**
**CD45$^+$CD3$^+$TCRβ$^+$**

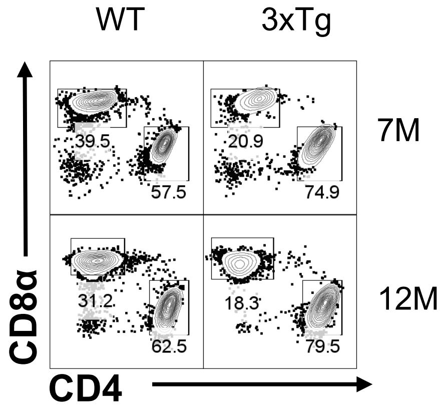


**3xTg SPF Males: MLN**
**CD45$^+$CD3$^+$TCRβ$^+$**

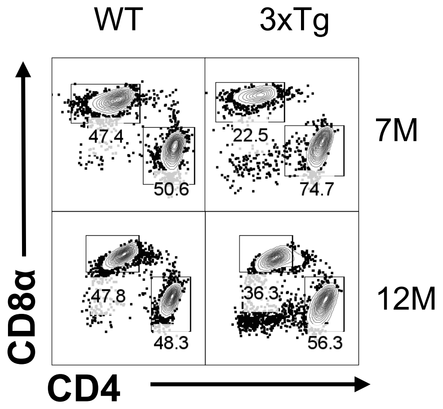


**3xTg SPF Males: SC**
**CD45$^+$CD3$^+$TCRβ$^+$**

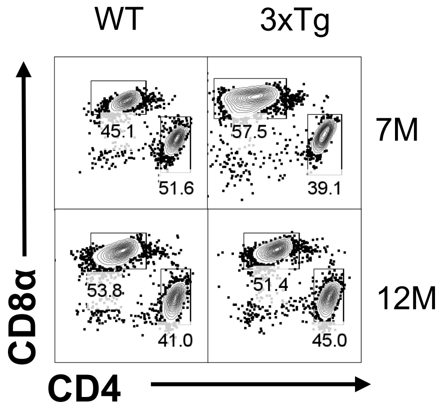


**3xTg SPF Males: DC**
**CD45$^+$CD3$^+$TCRβ$^+$**

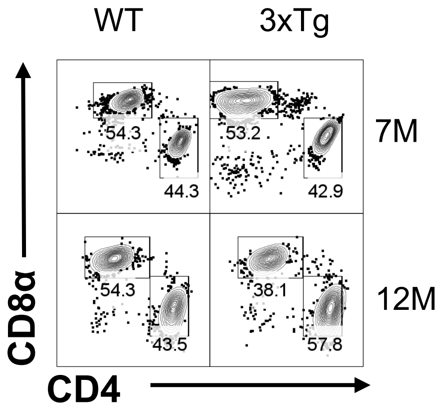

### Representative Flow Cytometry Plots for Fig. S4, S12: CD4$^+$ and CD8$^+$ T cells [3xTg GF Females]

**3xTg GF Females: SP**
**CD45$^+$CD3$^+$TCRβ$^+$**

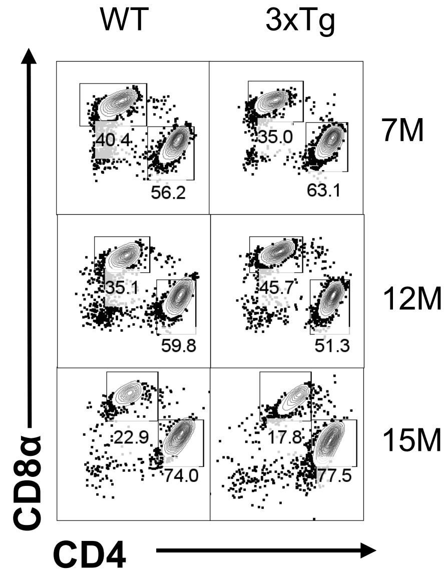


**3xTg GF Females: MLN**
**CD45$^+$CD3$^+$TCRβ$^+$**

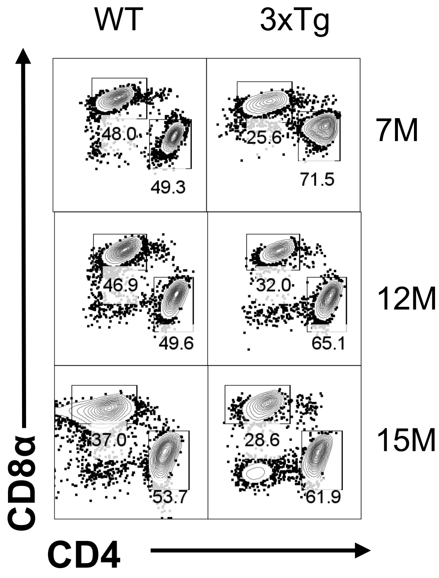


**3xTg GF Females: SC**
**CD45$^+$CD3$^+$TCRβ$^+$**

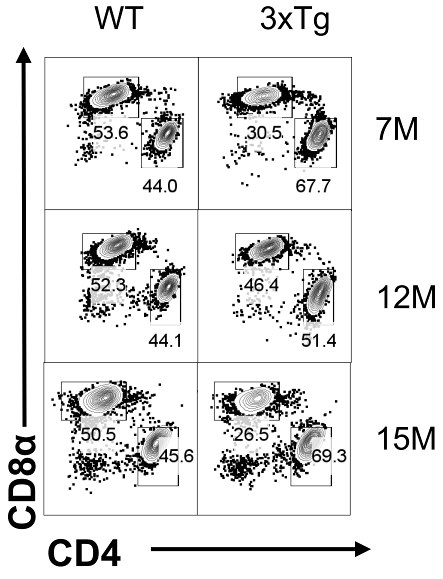


**3xTg GF Females: DC**
**CD45$^+$CD3$^+$TCRβ$^+$**

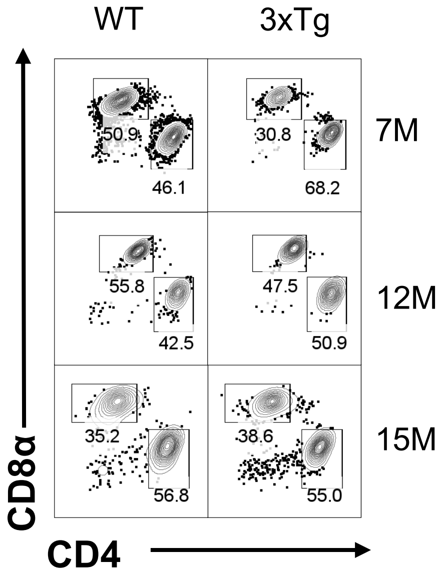

### Representative Flow Cytometry Plots for Fig. S4, S12: $CD4^{+}$ and $CD8^{+}$ T cells [3xTg GF Males]

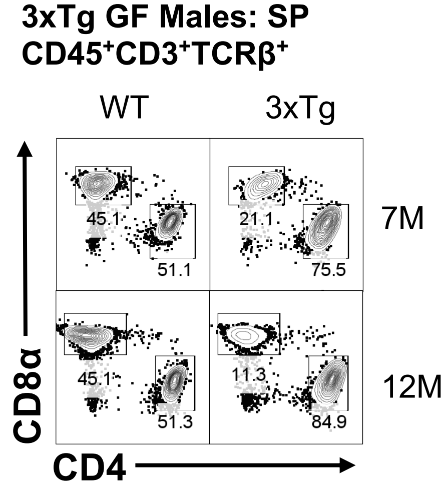


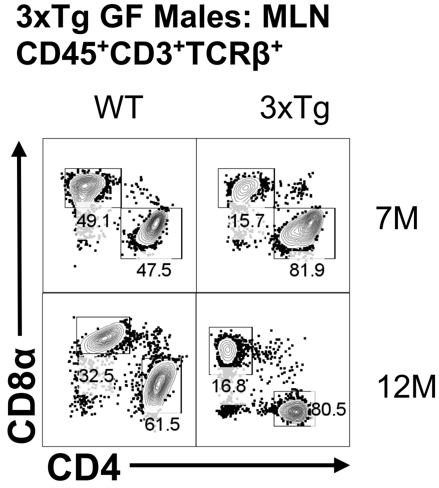


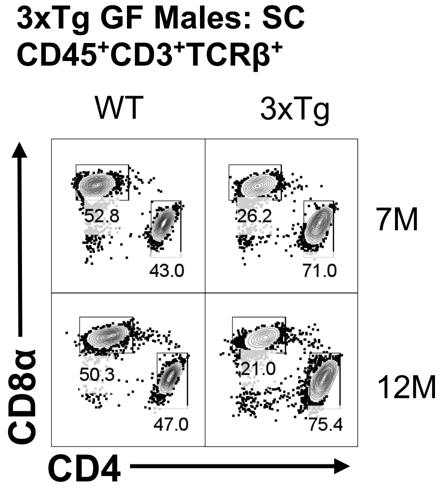


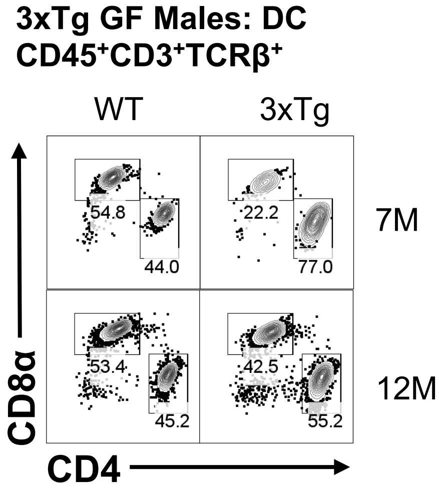

### Representative Flow Cytometry Plots for Fig. S5: IFNγ [3xTg]

**3xTg 12M Females: SP**
**CD45$^+$CD4$^+$TCRβ$^+$Foxp3$^-$**

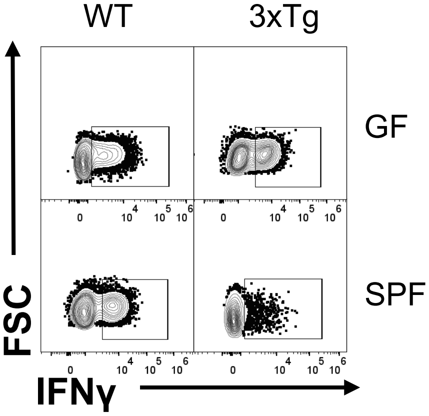


**3xTg 12M Females: MLN**
**CD45$^+$CD4$^+$TCRβ$^+$Foxp3$^-$**

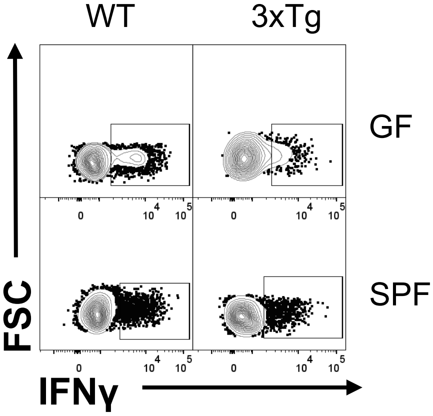


**3xTg 12M Females: SC**
**CD45$^+$CD4$^+$TCRβ$^+$Foxp3$^-$**

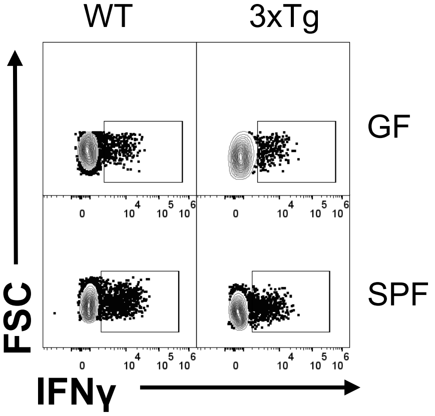


**3xTg 12M Females: DC**
**CD45$^+$CD4$^+$TCRβ$^+$Foxp3$^-$**

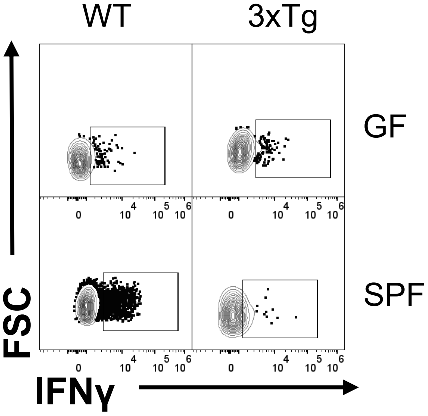


**3xTg 12M Males: SP**
**CD45$^+$CD4$^+$TCRβ$^+$Foxp3$^-$**

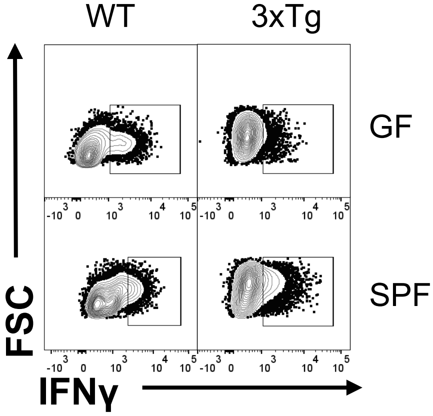


**3xTg 12M Males: MLN**
**CD45$^+$CD4$^+$TCRβ$^+$Foxp3$^-$**

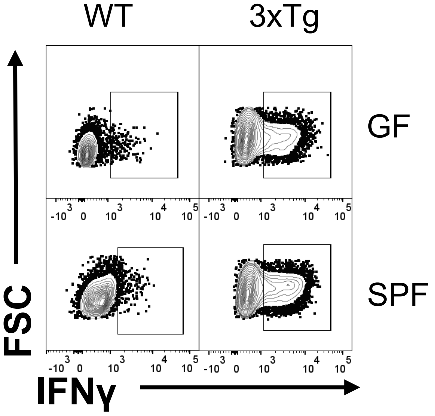


**3xTg 7M Males: SC**
**CD45$^+$CD4$^+$TCRβ$^+$Foxp3$^-$**

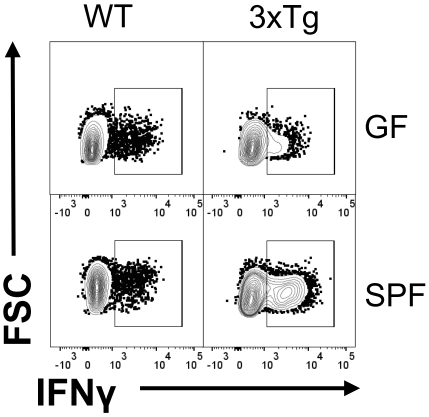


**3xTg 7M Males: DC**
**CD45$^+$CD4$^+$TCRβ$^+$Foxp3$^-$**

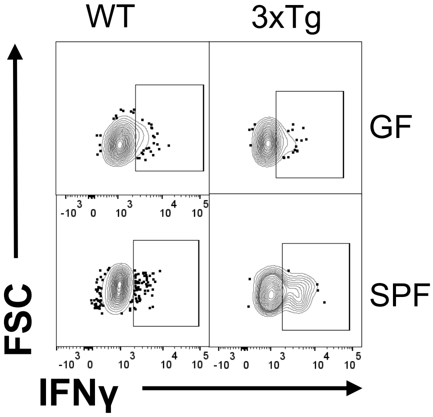

### Representative Flow Cytometry Plots for Fig. S6: Foxp3⁺ T cells [3xTg]

**3xTg 12M Females: SP**
**CD45⁺CD4⁺TCRβ⁺**

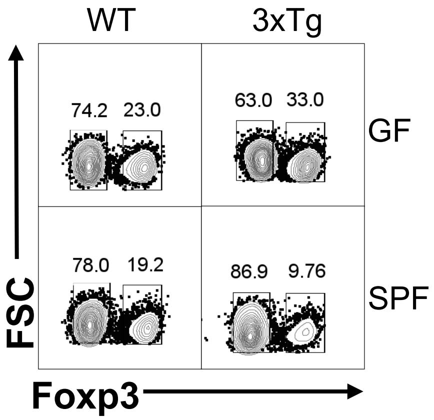


**3xTg 12M Females: MLN**
**CD45⁺CD4⁺TCRβ⁺**

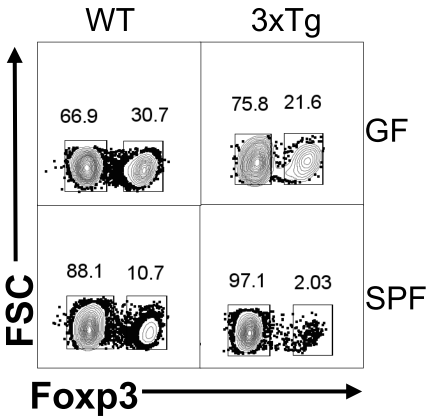


**3xTg 12M Females: SC**
**CD45⁺CD4⁺TCRβ⁺**

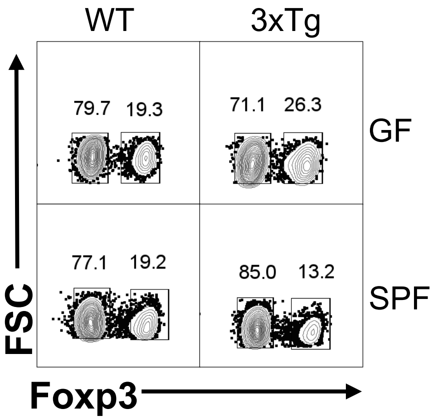


**3xTg 12M Females: DC**
**CD45⁺CD4⁺TCRβ⁺**

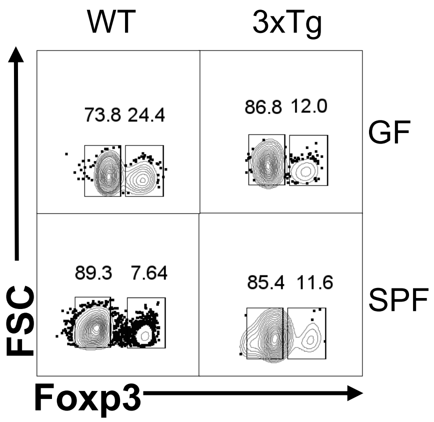


**3xTg 12M Males: SP**
**CD45⁺CD4⁺TCRβ⁺**

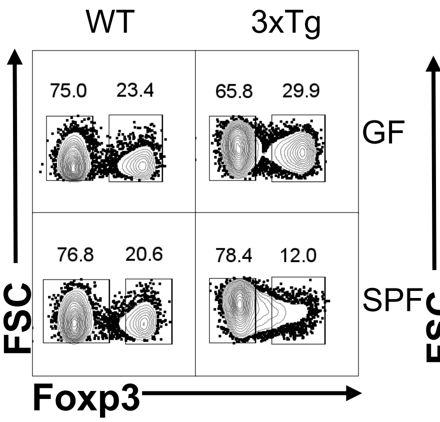


**3xTg 12M Males: MLN**
**CD45⁺CD4⁺TCRβ⁺**

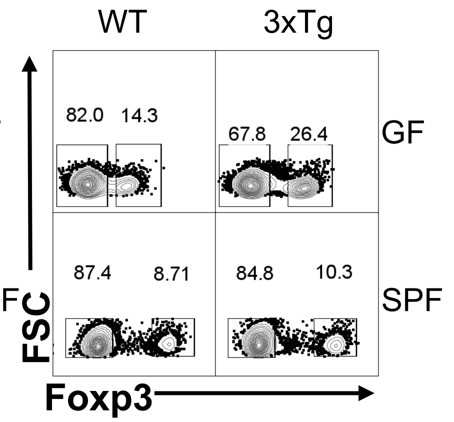


**3xTg 12M Males: SC**
**CD45⁺CD4⁺TCRβ⁺**

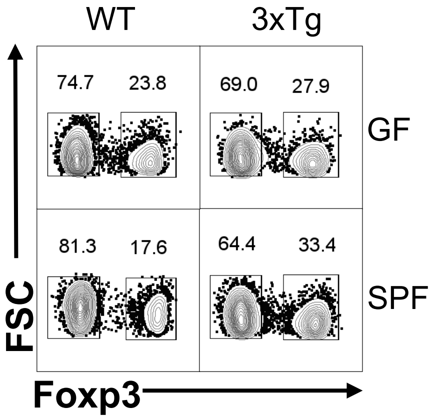


**3xTg 12M Males: DC**
**CD45⁺CD4⁺TCRβ⁺**

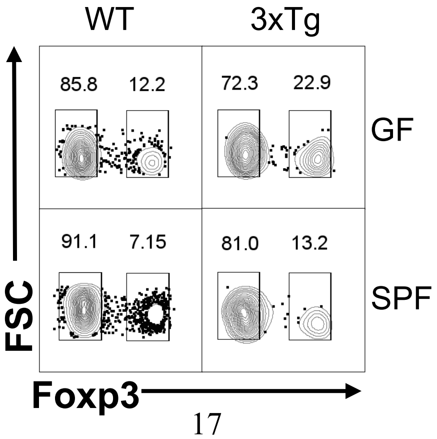

### Representative Flow Cytometry Plots for Fig. S8, S9: IFNγ+/GM-CSF+ T cells [5xFAD Females]

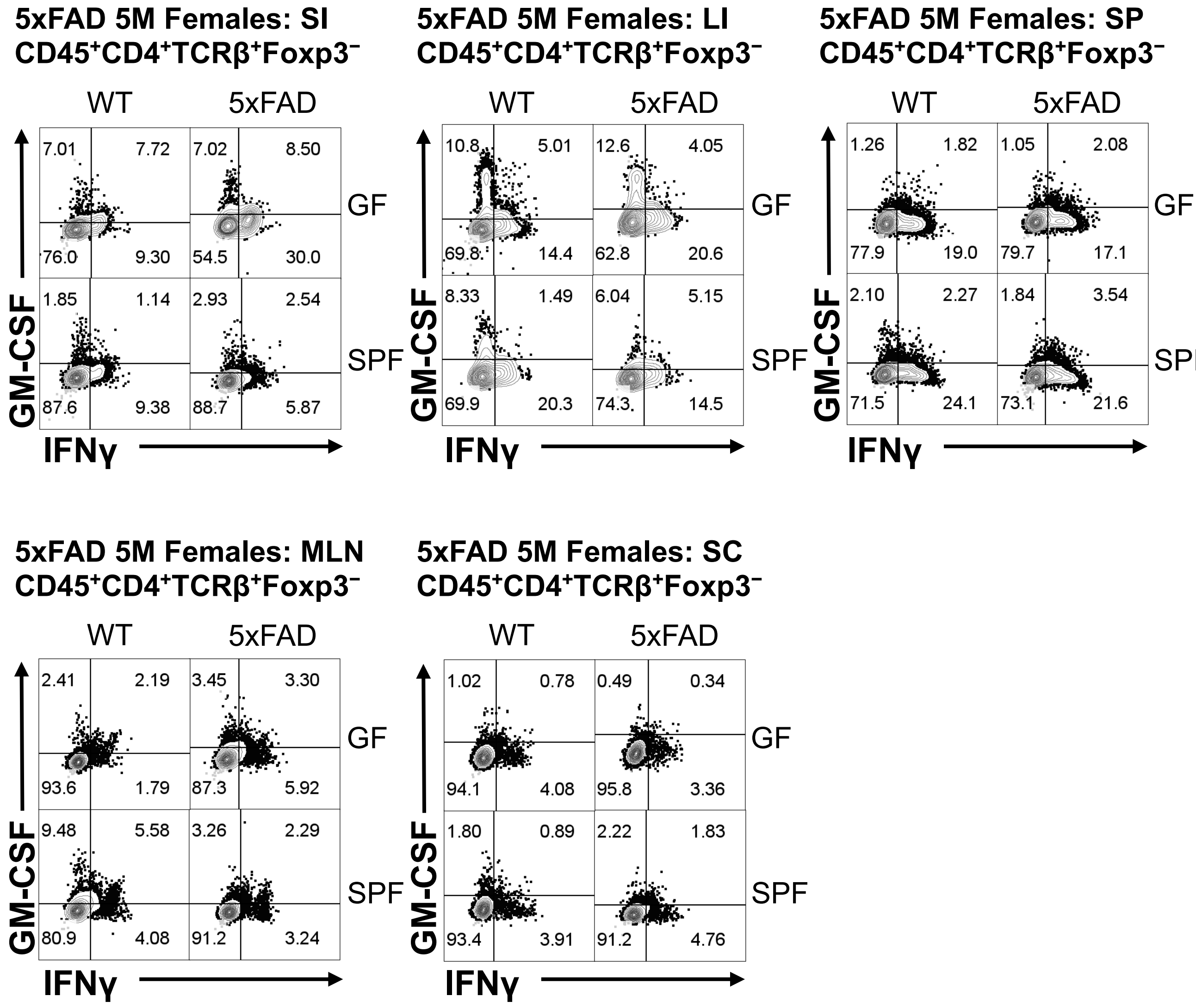

### Representative Flow Cytometry Plots for Fig. S8, S9: IFNγ⁺/GM-CSF⁺ T cells [5xFAD Males]

**5xFAD 5M Males: SI**
**CD45⁺CD4⁺TCRβ⁺Foxp3⁻**

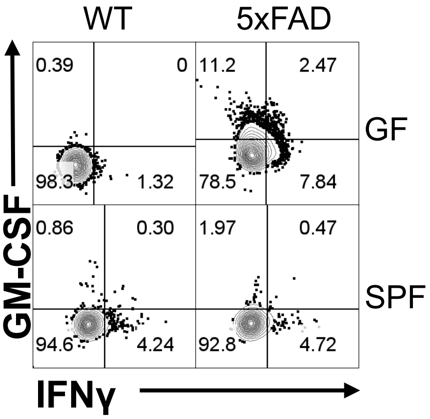


**5xFAD 5M Males: LI**
**CD45⁺CD4⁺TCRβ⁺Foxp3⁻**

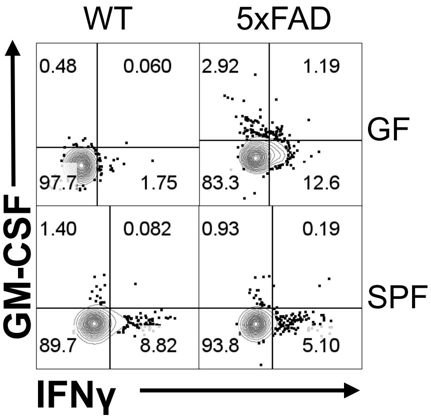


**5xFAD 5M Males: SP**
**CD45⁺CD4⁺TCRβ⁺Foxp3⁻**

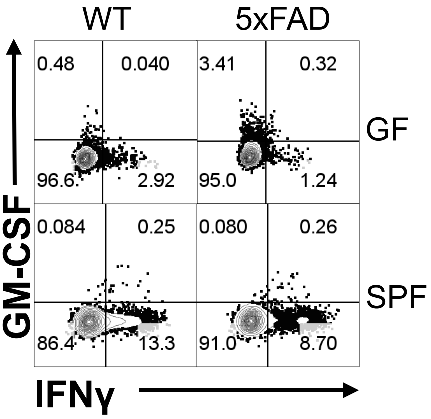


**5xFAD 5M Males: MLN**
**CD45⁺CD4⁺TCRβ⁺Foxp3⁻**

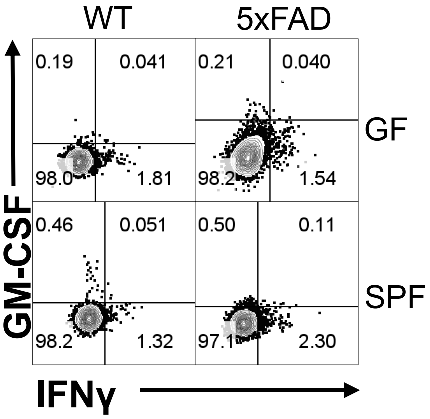


**5xFAD 5M Males: SC**
**CD45⁺CD4⁺TCRβ⁺Foxp3⁻**

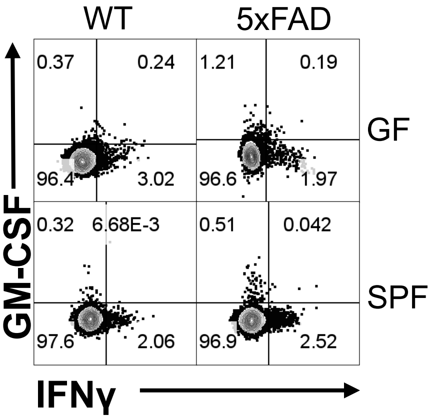

### Representative Flow Cytometry Plots for Fig. S8: IFNγ⁺ MFI [5xFAD Females]

**5xFAD 5M Females: SI**
**CD45⁺CD4⁺TCRβ⁺Foxp3⁻**

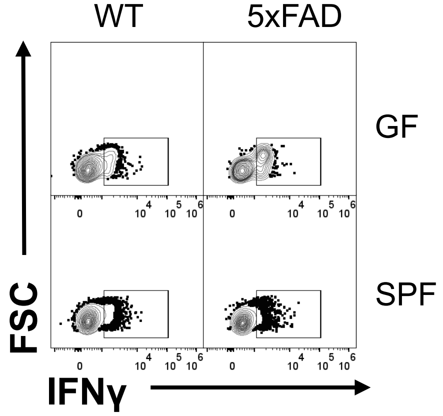


**5xFAD 5M Females: LI**
**CD45⁺CD4⁺TCRβ⁺Foxp3⁻**

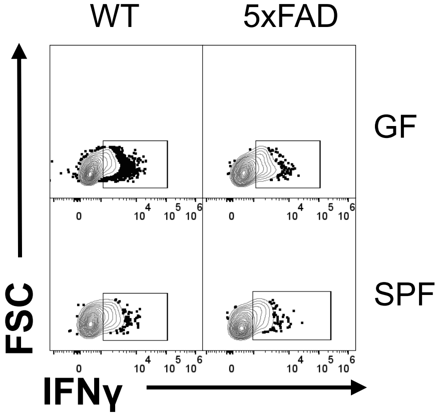


**5xFAD 5M Females: SP**
**CD45⁺CD4⁺TCRβ⁺Foxp3⁻**

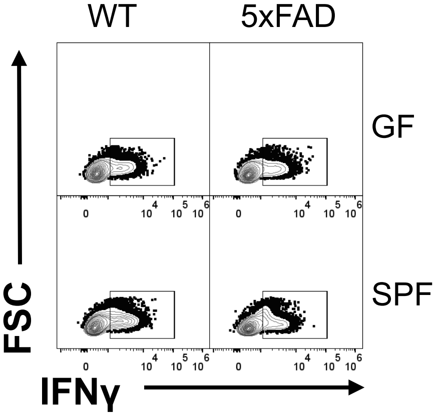


**5xFAD 5M Females: MLN**
**CD45⁺CD4⁺TCRβ⁺Foxp3⁻**

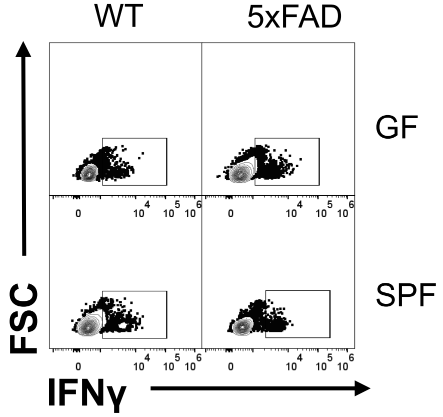


**5xFAD 5M Females: SC**
**CD45⁺CD4⁺TCRβ⁺Foxp3⁻**

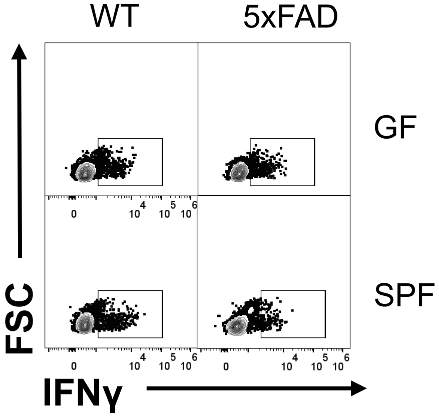

### Representative Flow Cytometry Plots for Fig. S8: IFNγ⁺ MFI [5xFAD Males]

**5xFAD 5M Males: SI**
**$CD45^+CD4^+TCR\beta^+Foxp3^-$**

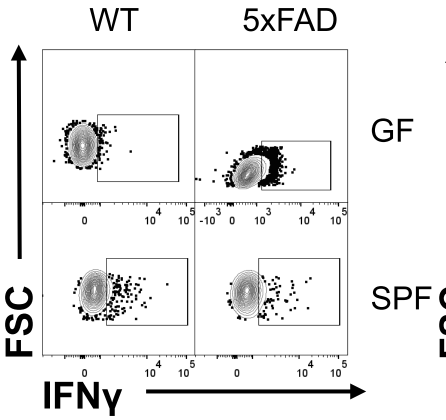


**5xFAD 5M Males: LI**
**$CD45^+CD4^+TCR\beta^+Foxp3^-$**

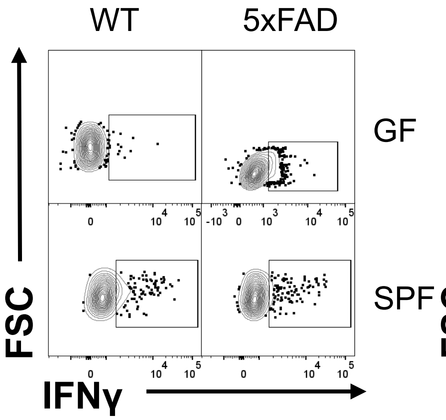


**5xFAD 5M Males: SP**
**$CD45^+CD4^+TCR\beta^+Foxp3^-$**

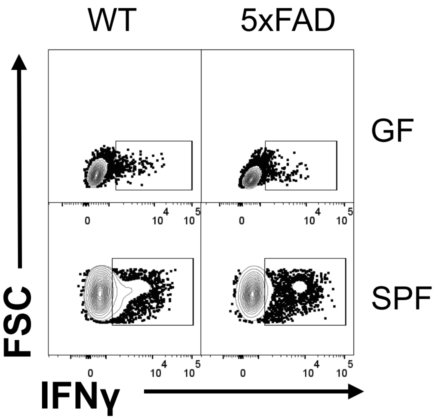


**5xFAD 5M Males: MLN**
**$CD45^+CD4^+TCR\beta^+Foxp3^-$**

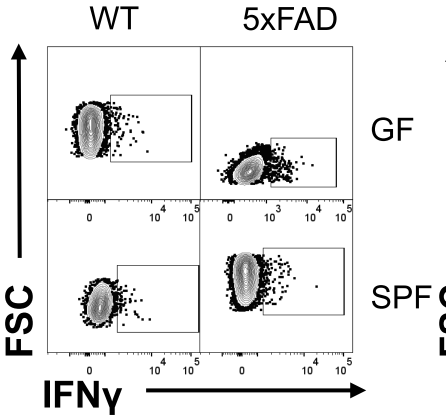


**5xFAD 5M Males: SC**
**$CD45^+CD4^+TCR\beta^+Foxp3^-$**

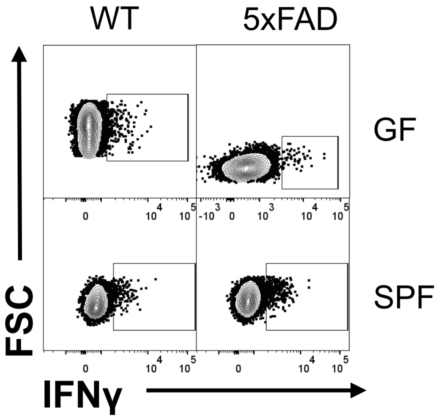

### Representative Flow Cytometry Plots for Fig. S9: GM-CSF$^{+}$ MFI [5xFAD Females]

**5xFAD 5M Females: SI**
**CD45$^{+}$CD4$^{+}$TCRβ$^{+}$Foxp3$^{-}$**

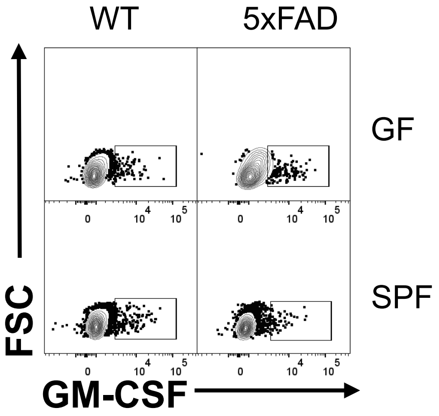


**5xFAD 5M Females: LI**
**CD45$^{+}$CD4$^{+}$TCRβ$^{+}$Foxp3$^{-}$**

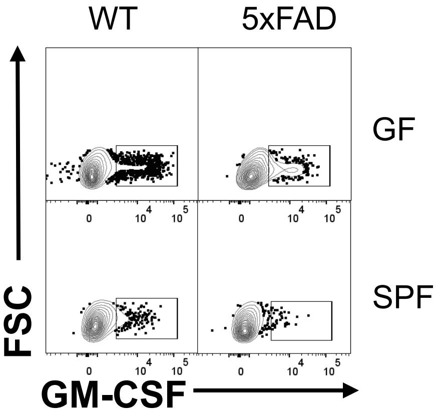


**5xFAD 5M Females: SP**
**CD45$^{+}$CD4$^{+}$TCRβ$^{+}$Foxp3$^{-}$**

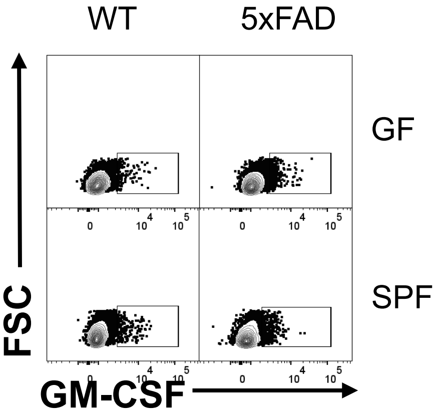


**5xFAD 5M Females: MLN**
**CD45$^{+}$CD4$^{+}$TCRβ$^{+}$Foxp3$^{-}$**

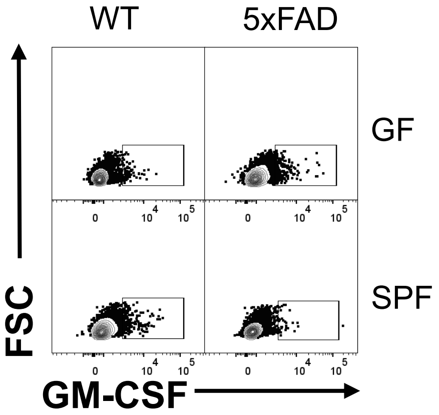


**5xFAD 5M Females: SC**
**CD45$^{+}$CD4$^{+}$TCRβ$^{+}$Foxp3$^{-}$**

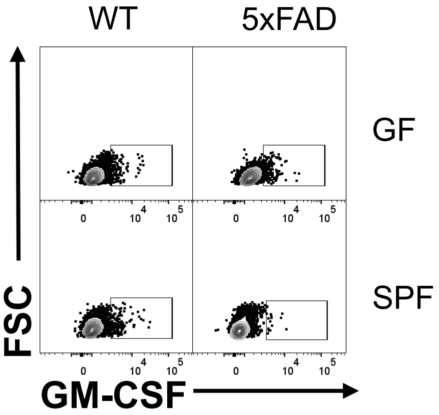

### Representative Flow Cytometry Plots for Fig. S9: GM-CSF+ MFI [5xFAD Males]

**5xFAD 5M Males: SI**
**$CD45^+CD4^+TCR\beta^+Foxp3^-$**

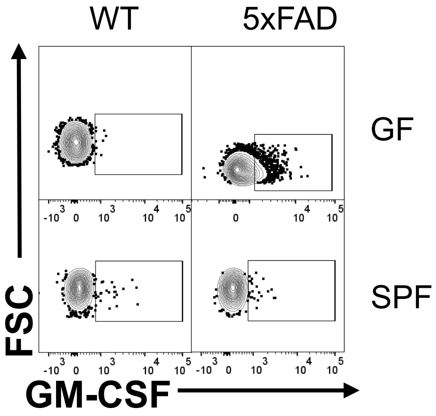


**5xFAD 5M Males: LI**
**$CD45^+CD4^+TCR\beta^+Foxp3^-$**

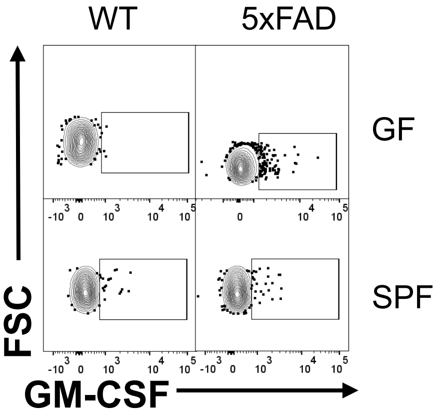


**5xFAD 5M Males: SP**
**$CD45^+CD4^+TCR\beta^+Foxp3^-$**

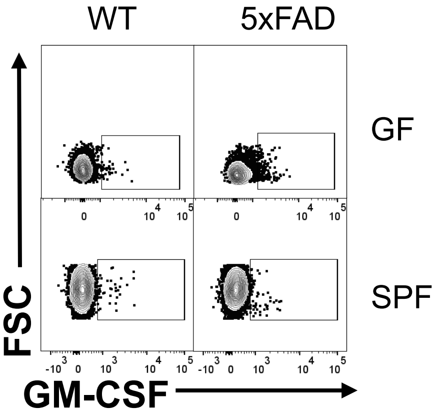


**5xFAD 5M Males: MLN**
**$CD45^+CD4^+TCR\beta^+Foxp3^-$**

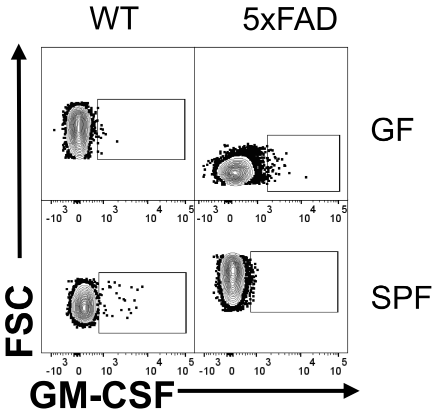


**5xFAD 5M Males: SC**
**$CD45^+CD4^+TCR\beta^+Foxp3^-$**

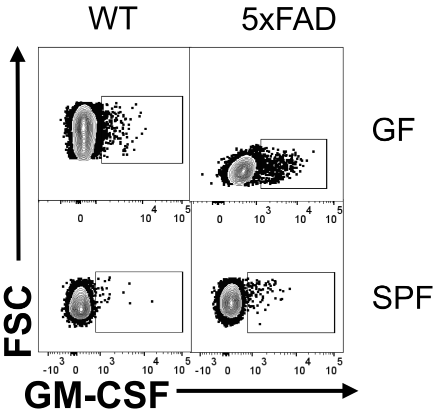

### Representative Flow Cytometry Plots for Fig. S10: IFNγ+/IL-4+ T cells [5xFAD Females]

**5xFAD 5M Females: SI**
**CD45+CD4+TCRβ+Foxp3−**

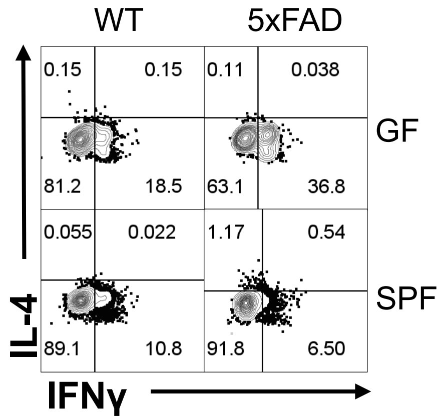


**5xFAD 5M Females: LI**
**CD45+CD4+TCRβ+Foxp3−**

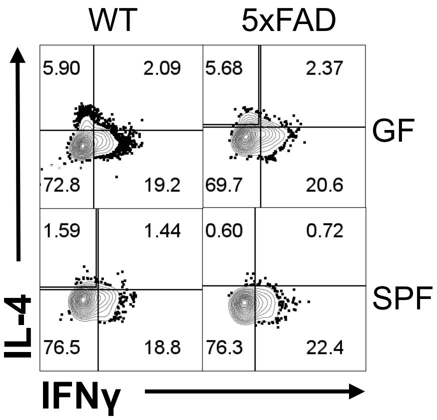


**5xFAD 5M Females: SP**
**CD45+CD4+TCRβ+Foxp3−**

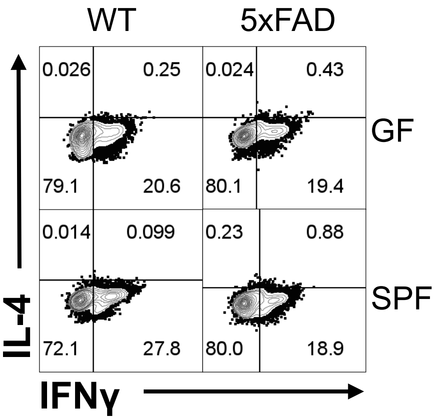


**5xFAD 5M Females: MLN**
**CD45+CD4+TCRβ+Foxp3−**

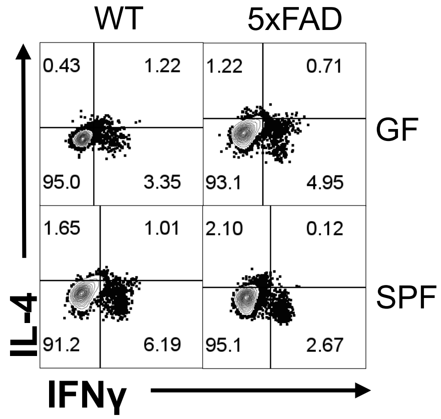


**5xFAD 5M Females: SC**
**CD45+CD4+TCRβ+Foxp3−**

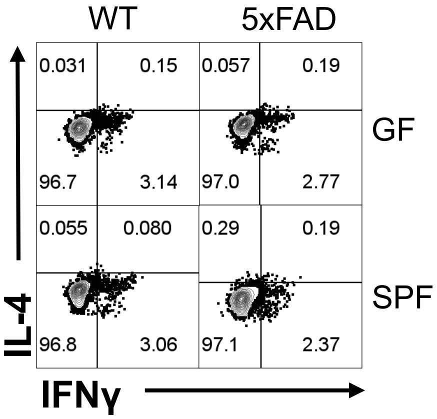

### Representative Flow Cytometry Plots for Fig. S10: IFNγ+/IL-4+ T cells [5xFAD Males]

**5xFAD 5M Males: SI**
**CD45+CD4+TCRβ+Foxp3–**

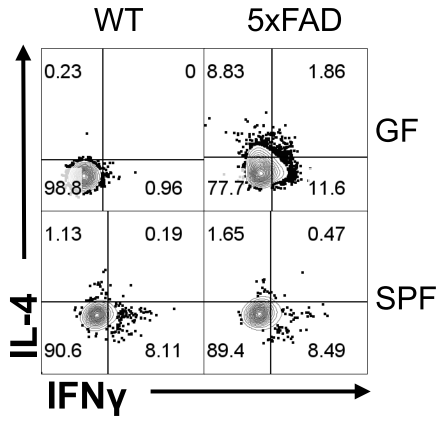


**5xFAD 5M Males: LI**
**CD45+CD4+TCRβ+Foxp3–**

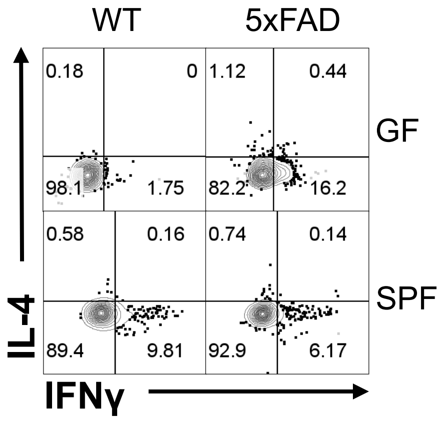


**5xFAD 5M Males: SP**
**CD45+CD4+TCRβ+Foxp3–**

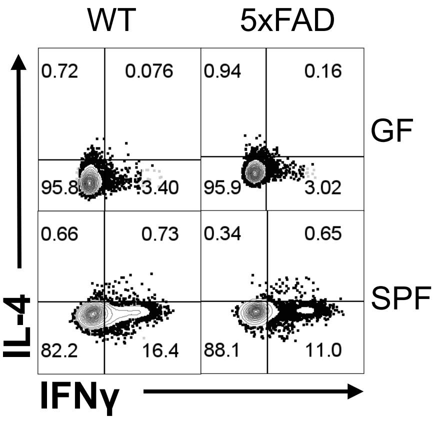


**5xFAD 5M Males: MLN**
**CD45+CD4+TCRβ+Foxp3–**

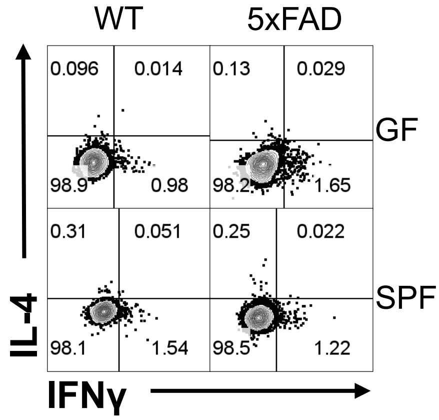


**5xFAD 5M Males: SC**
**CD45+CD4+TCRβ+Foxp3–**

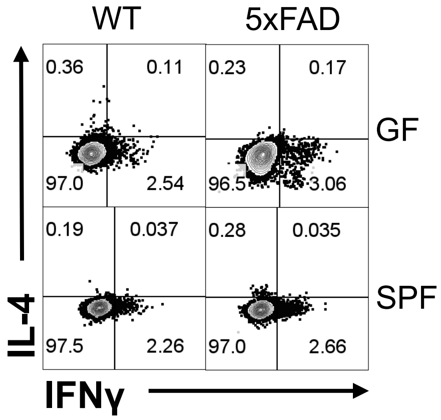

### Representative Flow Cytometry Plots for Fig. S10: IL-4⁺ MFI [5xFAD Females]

**5xFAD 5M Females: SI**
**CD45⁺CD4⁺TCRβ⁺Foxp3⁻**

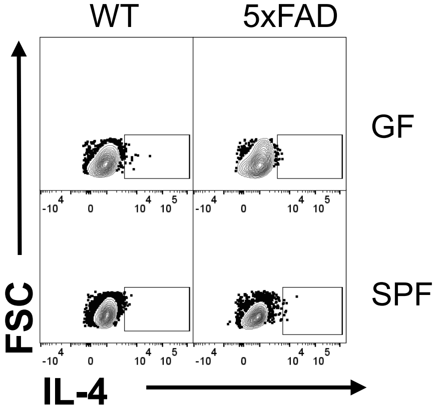


**5xFAD 5M Females: LI**
**CD45⁺CD4⁺TCRβ⁺Foxp3⁻**

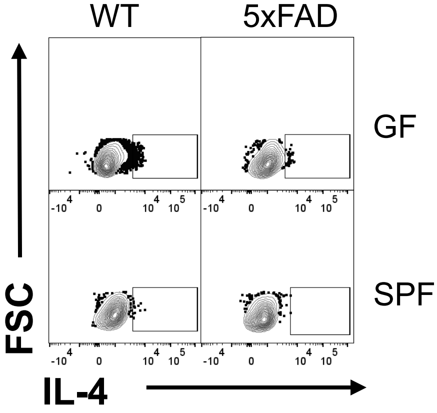


**5xFAD 5M Females: SP**
**CD45⁺CD4⁺TCRβ⁺Foxp3⁻**

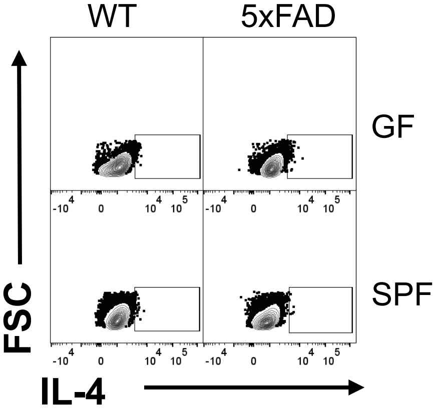


**5xFAD 5M Females: MLN**
**CD45⁺CD4⁺TCRβ⁺Foxp3⁻**

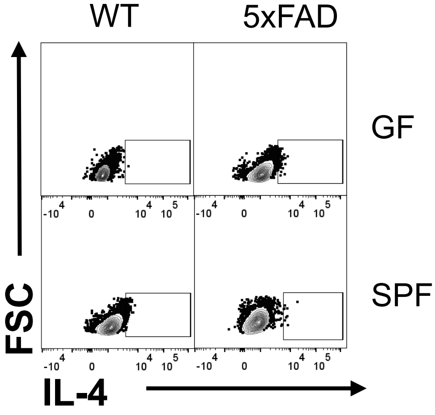


**5xFAD 5M Females: SC**
**CD45⁺CD4⁺TCRβ⁺Foxp3⁻**

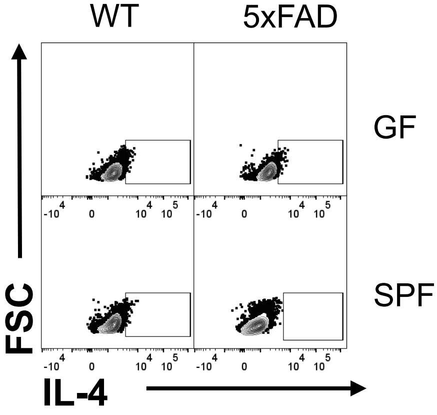

### Representative Flow Cytometry Plots for Fig. S10: IL-4$^+$ MFI [5xFAD Males]

**5xFAD 5M Males: SI**
**CD45$^+$CD4$^+$TCRβ$^+$Foxp3$^-$**

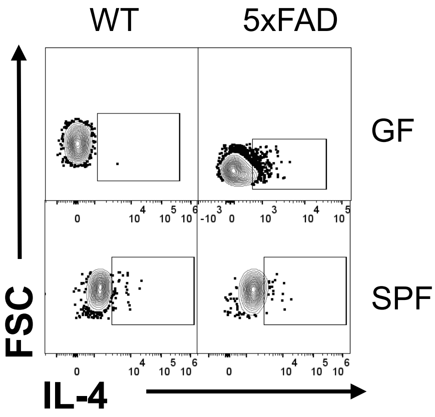


**5xFAD 5M Males: LI**
**CD45$^+$CD4$^+$TCRβ$^+$Foxp3$^-$**

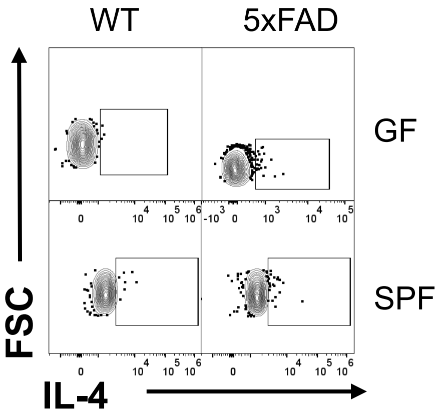


**5xFAD 5M Males: SP**
**CD45$^+$CD4$^+$TCRβ$^+$Foxp3$^-$**

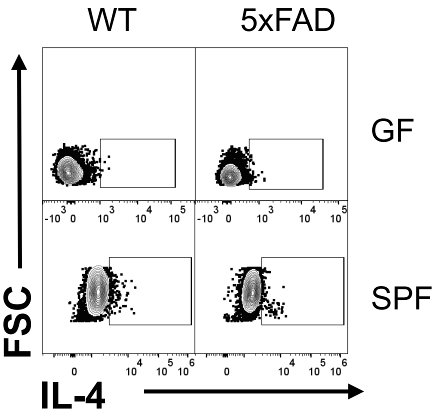


**5xFAD 5M Males: MLN**
**CD45$^+$CD4$^+$TCRβ$^+$Foxp3$^-$**

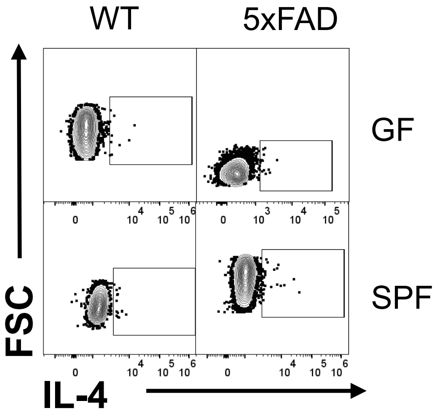


**5xFAD 5M Males: SC**
**CD45$^+$CD4$^+$TCRβ$^+$Foxp3$^-$**

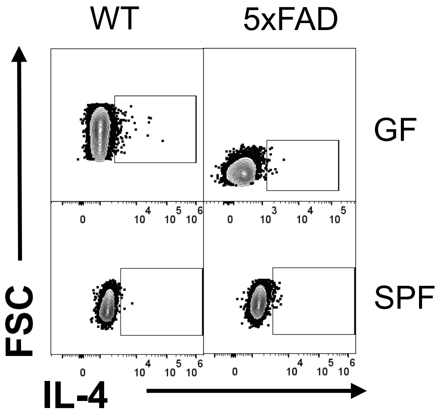

### Representative Flow Cytometry Plots for Fig. S11: Foxp3$^+$ T cells [5xFAD Females]

**5xFAD 5M Females: SI**
**CD45$^+$CD4$^+$TCRβ$^+$**

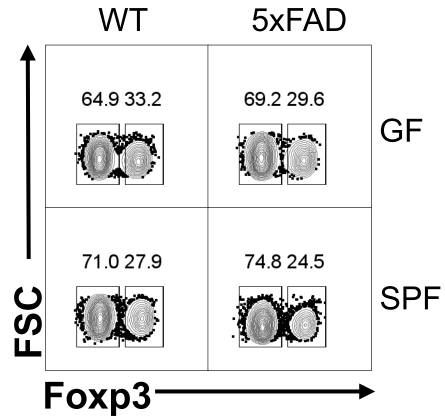


**5xFAD 5M Females: LI**
**CD45$^+$CD4$^+$TCRβ$^+$**

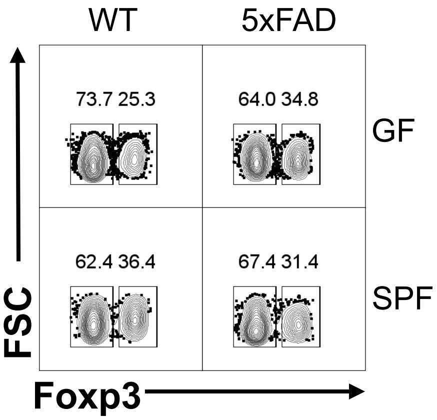


**5xFAD 5M Females: SP**
**CD45$^+$CD4$^+$TCRβ$^+$**

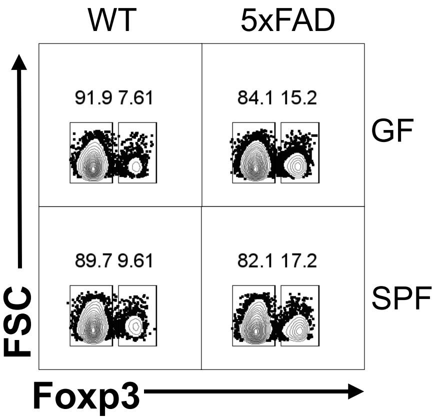


**5xFAD 5M Females: MLN**
**CD45$^+$CD4$^+$TCRβ$^+$**

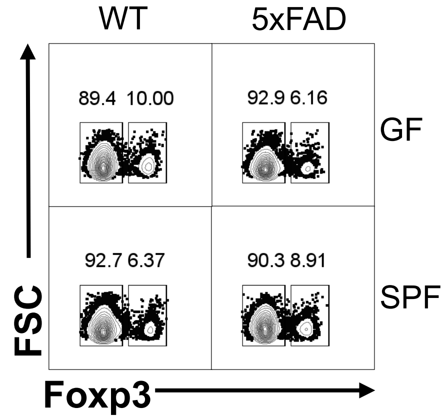


**5xFAD 5M Females: SC**
**CD45$^+$CD4$^+$TCRβ$^+$**

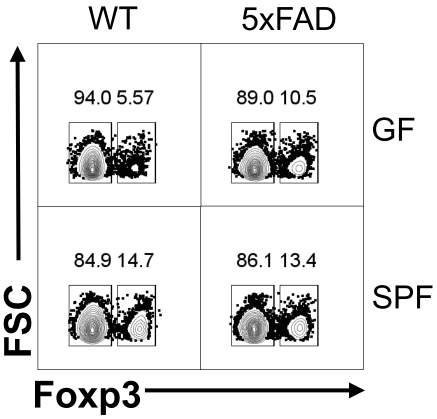

### Representative Flow Cytometry Plots for Fig. S11: Foxp3$^{+}$ T cells [5xFAD Males]

**5xFAD 5M Males: SI**
**CD45$^{+}$CD4$^{+}$TCRβ$^{+}$**

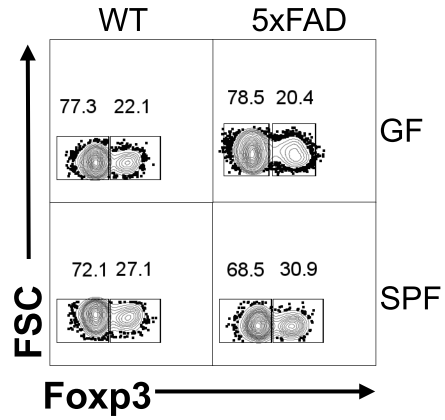


**5xFAD 5M Males: LI**
**CD45$^{+}$CD4$^{+}$TCRβ$^{+-}$**

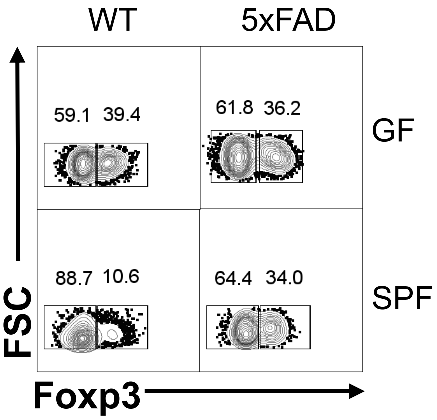


**5xFAD 5M Males: SP**
**CD45$^{+}$CD4$^{+}$TCRβ$^{+}$**

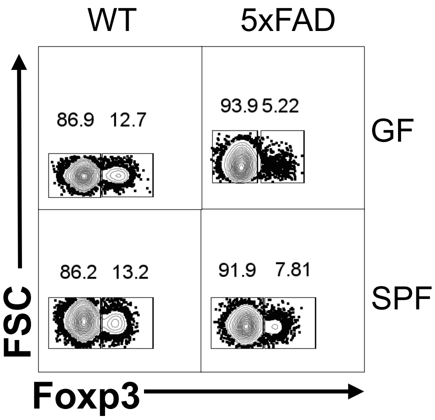


**5xFAD 5M Males: MLN**
**CD45$^{+}$CD4$^{+}$TCRβ$^{+}$**

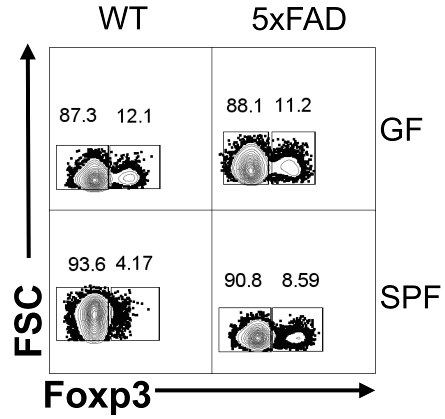


**5xFAD 5M Males: SC**
**CD45$^{+}$CD4$^{+}$TCRβ$^{+}$**

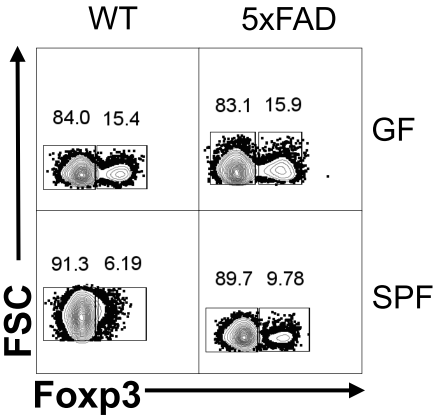

### Representative Flow Cytometry Plots for Fig. S13: T and B cells [3xTg SPF Males]

**3xTg SPF Males: SP**
**CD45$^{+}$**

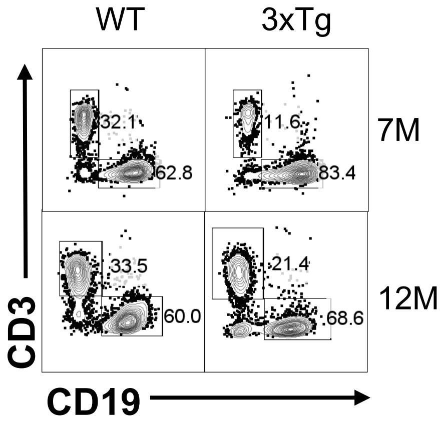


**3xTg SPF Males: MLN**
**CD45$^{+}$**

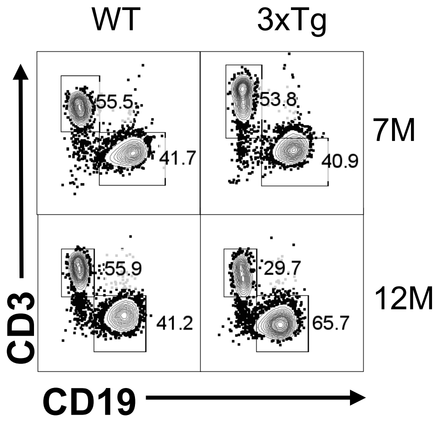


**3xTg SPF Males: SC**
**CD45$^{+}$**

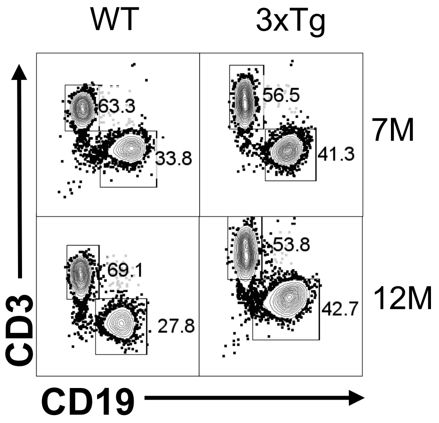


**3xTg SPF Males: DC**
**CD45$^{+}$**

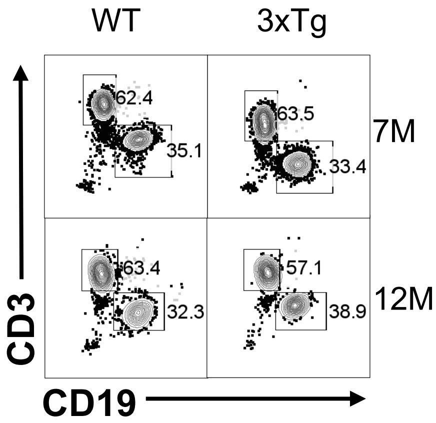

### Representative Flow Cytometry Plots for Fig. S13: T and B cells [3xTg GF Females]

**3xTg GF Females: SP**
**$CD45^+$**

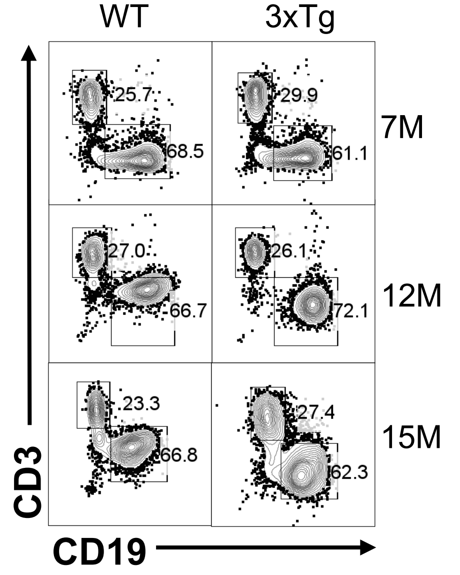


**3xTg GF Females: MLN**
**$CD45^+$**

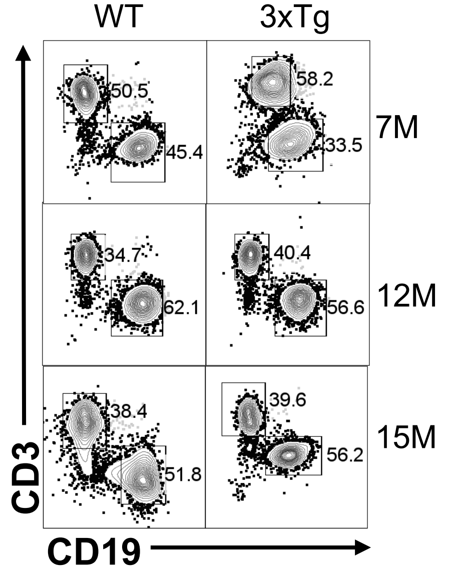


**3xTg GF Females: SC**
**$CD45^+$**

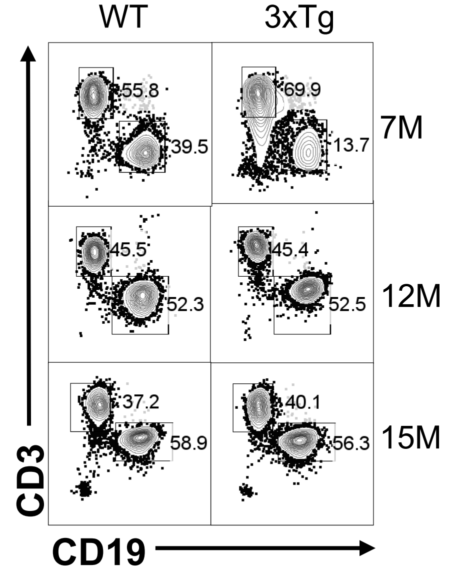


**3xTg GF Females: DC**
**$CD45^+$**

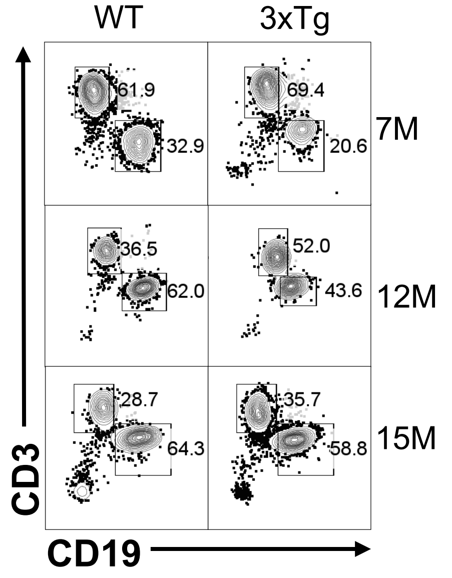

### Representative Flow Cytometry Plots for Fig. S13: T and B cells [3xTg GF Males]

**3xTg GF Males: SP**
**CD45$^+$**

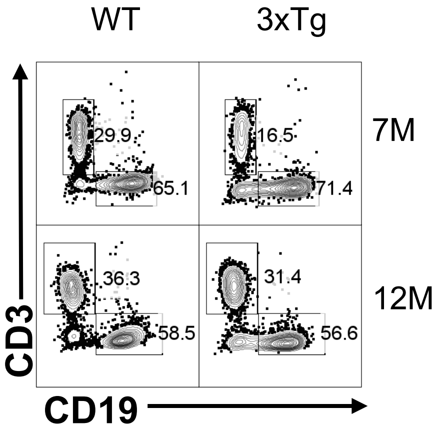


**3xTg GF Males: MLN**
**CD45$^+$**

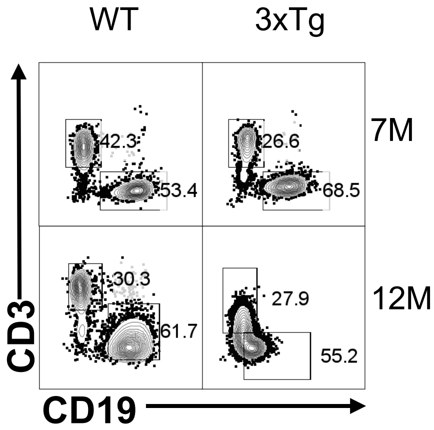


**3xTg GF Males: SC**
**CD45$^+$**

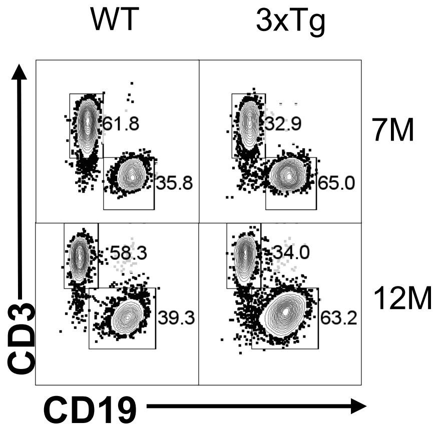


**3xTg GF Males: DC**
**CD45$^+$**

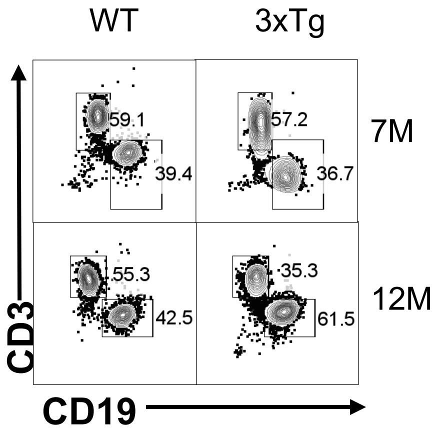

### Representative Flow Cytometry Plots for Fig. S14: T and B cells [5xFAD 8M]

**5xFAD 8M Females: SP**
**CD45+**

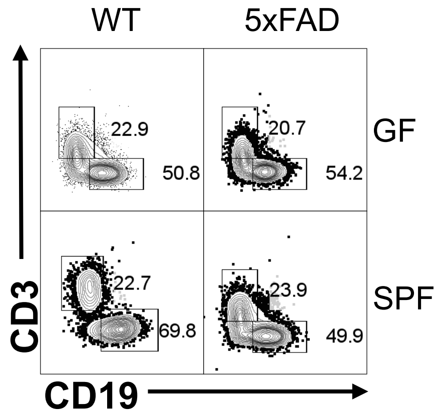


**5xFAD 8M Females: MLN**
**CD45+**

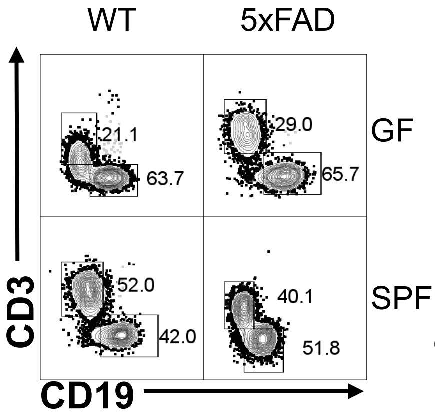


**5xFAD 8M Females: SC**
**CD45+**

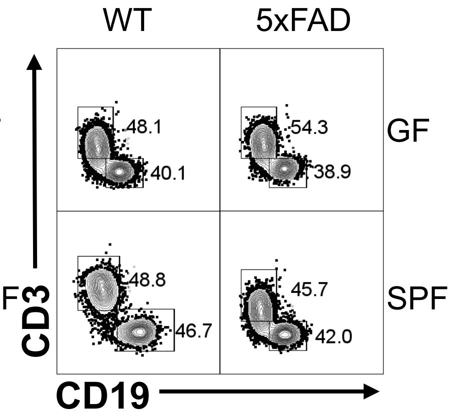


**5xFAD 8M Females: DC**
**CD45+**

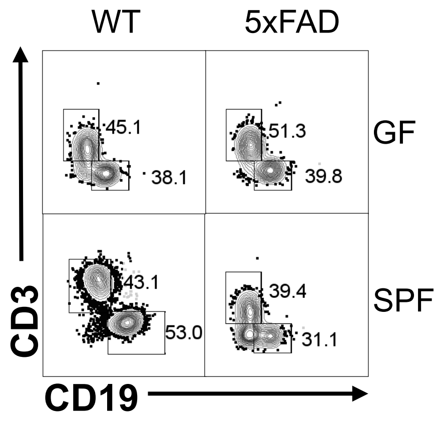


**5xFAD 8M Males: SP**
**CD45+**

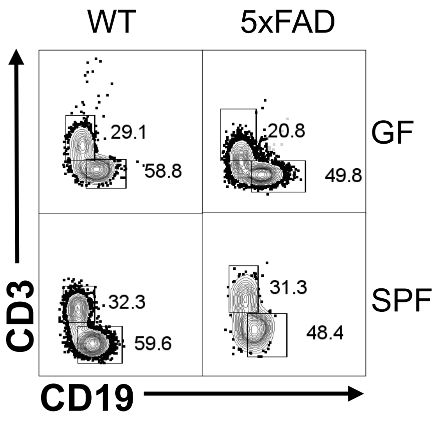


**5xFAD 8M Males: MLN**
**CD45+**

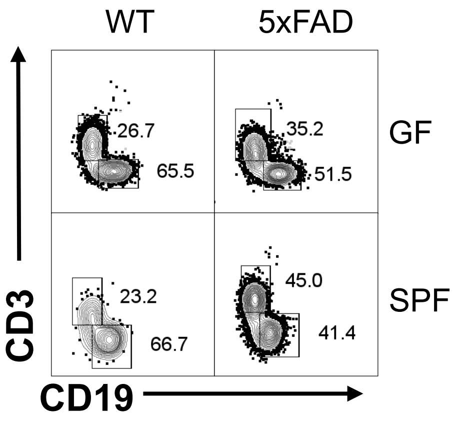


**5xFAD 8M Males: SC**
**CD45+**

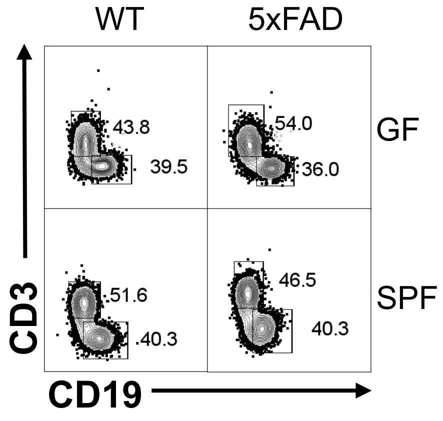


**5xFAD 8M Males: DC**
**CD45+**

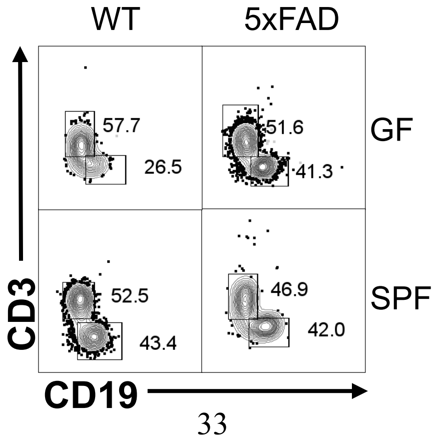
